# The ultrasonic vocalization (USV) syllable profile during neonatal opioid withdrawal and a kappa opioid receptor component to increased USV emissions in female mice

**DOI:** 10.1101/2024.07.02.601766

**Authors:** Kelly K. Wingfield, Teodora Misic, Kaahini Jain, Carly S. McDermott, Nalia M. Abney, Kayla T. Richardson, Mia B. Rubman, Jacob A. Beierle, Sophia A. Miracle, Emma J. Sandago, Britahny M. Baskin, William B. Lynch, Kristyn N. Borrelli, Emily J. Yao, Elisha M. Wachman, Camron D. Bryant

**Author notes:** Corresponding Author, Camron D. Bryant, Ph.D., Laboratory of Addiction Genetics, Department for Pharmaceutical Sciences, Center for Drug Discovery, Northeastern University, 140 The Fenway, X138 Boston, MA USA 02115, E. **FUNDING** U01DA050243 (C.D.B.). U01DA55299 (C.D.B.), T32DA055553 (B.M.B.).

## Abstract

**Rationale:** Opioid use during pregnancy can lead to negative infant health outcomes, including neonatal opioid withdrawal syndrome (NOWS). NOWS comprises gastrointestinal, autonomic nervous system, and neurological dysfunction that manifest during spontaneous withdrawal. Variability in NOWS severity necessitates a more individualized treatment approach. Ultrasonic vocalizations (USVs) in neonatal mice are emitted in isolation as a stress response and are increased during opioid withdrawal, thus modeling a negative affective state that can be utilized to test new treatments.

**Objectives:** We sought to identify the behavioral and USV profile, brainstem transcriptomic adaptations, and role of kappa opioid receptors in USVs during neonatal opioid withdrawal.

**Methods:** We employed a third trimester-approximate opioid exposure model, where neonatal inbred FVB/NJ pups were injected twice-daily with morphine (10mg/kg, s.c.) or saline (0.9%, 20 ul/g, s.c.) from postnatal day(P) 1 to P14. This protocol induces reduced weight gain, hypothermia, thermal hyperalgesia, and increased USVs during spontaneous morphine withdrawal.

**Results:** On P14, there were increased USV emissions and altered USV syllables during withdrawal, including an increase in Complex 3 syllables in FVB/NJ females (but not males). Brainstem bulk mRNA sequencing revealed an upregulation of the kappa opioid receptor (*Oprk1)*, which contributes to withdrawal-induced dysphoria. The kappa opioid receptor (KOR) antagonist, nor-BNI (30 mg/kg, s.c.), significantly reduced USVs in FVB/NJ females, but not males during spontaneous morphine withdrawal. Furthermore, the KOR agonist, U50,488h (0.625 mg/kg, s.c.), was sufficient to increase USVs on P10 (both sexes) and P14 (females only) in FVB/NJ mice.

**Conclusions:** We identified an elevated USV syllable, Complex 3, and a female-specific recruitment of the dynorphin/KOR system in increased USVs associated with neonatal opioid withdrawal severity.

## INTRODUCTION

Opioid use during pregnancy is a major public health concern (Goetz et al., 2021; Haight, 2018; Hales et al., 2020; Ko et al., 2020; VanHouten et al., 2019). Exposure to opioids during gestation can lead to negative health outcomes for infants, such as preterm birth, growth restriction, and neonatal opioid withdrawal syndrome (NOWS). Approximately 59 infants in the United States are diagnosed with NOWS daily and are at an increased risk for admission to the neonatal intensive care unit (NICU), longer hospital stays, and greater hospital costs compared to other infants (Haight, 2018; Milliren et al., 2018; Strahan et al., 2020; Tolia et al., 2015; Wachman & Werler, 2019; Winkelman et al., 2018). NOWS consists of symptoms induced by spontaneous cessation of in-utero opioid exposure and is characterized by gastrointestinal irregularities (weight loss, poor feeding, diarrhea, vomiting), disrupted autonomous system (sweating, respiratory difficulties, sneezing) and neurological dysfunction (sleep disruptions, irritability, and excessive, high-pitched crying) (Jansson et al., 2009; Ko et al., 2017; Strahan et al., 2020). Current treatments for NOWS involve non-pharmacologic interventions, such as rooming-in and breastfeeding, as well as mu-opioid receptor agonist therapy (Abdel-Latif et al., 2006; Howard et al., 2017; Kraft et al., 2017; Wachman et al., 2018; Wachman & Werler, 2019). NOWS severity is highly variable, which can be attributed to multiple factors, including genetics, biological sex, type of opioid exposure, maternal polysubstance use, gestational age of exposure, and hospital care model (Liu et al., 2010; O’Connor et al., 2013; Seligman et al., 2008; Wachman et al., 2013, 2015). Thus, it is challenging to investigate the impact of individual factors on NOWS severity in a clinical setting. However, understanding the factors contributing to symptom severity can inform personalized treatment strategies to improve infant health outcomes.

Rodent models for NOWS traits are used to study factors influencing withdrawal symptom onset, duration, and severity (Byrnes & Vassoler, 2018; Ferrante & Blendy, 2024; Richardson et al., 2006). To effectively model NOWS traits in mice, we employ a third trimester-approximate opioid exposure paradigm, which is sufficient to induce a withdrawal state in pups (Borrelli et al., 2021; Dunn et al., 2023; Ferrante et al., 2022; Richardson et al., 2006; S. A. Robinson et al., 2020). P1–P14 in mice largely mirrors the neurodevelopmental events that occur during the third trimester of human pregnancy, including oligodendrocyte maturation, myelination, synaptogenesis, and establishment of the blood-brain barrier (Byrnes & Vassoler, 2018; Craig et al., 2003; Rice & Barone, 2000; Saunders et al., 2012; Semple et al., 2013). Other models include prenatal opioid exposure to the pups and dam; however, these models create challenges for accomplishing precise delivery of doses to individual pups, which is necessary for the study of factors (e.g., genetic) underlying individual differences in neonatal opioid withdrawal presence and severity. Prenatal models also necessitate maternal drug exposure, which, while valid for studying the human condition, complicates the understanding of innate individual differences underlying neonatal opioid withdrawal severity.

Here, we implemented a third trimester-approximate NOWS model (Borrelli et al., 2021; S. A. Robinson et al., 2020)in inbred FVB/NJ and outbred Swiss Webster Carworth Farms White (CFW) mouse pups to investigate the behavioral and transcriptional adaptations of chronic morphine exposure and withdrawal. We first used supervised machine learning to classify ultrasonic vocalization (USV) syllable profiles during spontaneous morphine withdrawal. Neonatal USVs are emitted exclusively in isolation to communicate discomfort and promote maternal attention (Branchi et al., 2001; Caruso et al., 2022; D’Amato et al., 2005; Scattoni et al., 2008). Thus, USVs can serve to model the negative affective state associated with morphine withdrawal. Next, we examined the transcriptomic changes in the brainstem (medulla and pons), a brain region containing multiple nuclei (e.g., locus coeruleus, rostroventral medulla) that contribute to neurobehavioral adaptations during opioid withdrawal, including enhanced nociception, dysphoria, and anhedonia (Basinger & Hogg, 2023; Bruchas et al., 2010; Downs & McElligott, 2022; Land et al., 2008; Shippenberg et al., 2007). We then investigated whether kappa opioid receptors (KOR), which are known to contribute to dysphoria during opioid withdrawal, contribute to USV emissions and syllable profiles during spontaneous morphine withdrawal. Together, our results identify a specific USV syllable, Complex 3, as a potential marker for the severity of withdrawal-associated dysphoria in mice, as well as a female-specific KOR component to increased USVs during spontaneous morphine withdrawal. These results support the extensive body of literature documenting sex differences in the dynorphin/KOR system in responses comprising negative affective-motivational states and emphasize the importance of biological sex in considering treatment strategies for opioid withdrawal (Becker & Chartoff, 2019; Conway et al., 2019; Fox & Sinha, 2009; Madurai et al., 2024).

## METHODS

### Mice

All experiments involving mice were conducted in accordance with the National Institutes of Health *Guide for the Care and Use of Laboratory Animals* and were approved by the Institutional Animal Care and Use Committees at Boston University and Northeastern University. FVB/NJ inbred mice were purchased at 8 weeks old from The Jackson Laboratory (Strain #001800). The FVB/NJ strain was chosen based on their excellent fecundity and the unexplored prospect of potentially exploiting closely related FVB substrains for mapping the genetic basis of NOWS model traits (Bryant et al., 2020). Outbred Cartworth Farm Webster (Swiss Webster) mice (8 weeks old) were purchased from Charles River Laboratories, and their historical dataset (Borrelli et al., 2021) was used to test for generalization of our findings, as they represent a commonly used and a relatively more diverse outbred genetic background. Mice were provided ad libitum laboratory breeding chow (Teklad 18% Protein Diet, Envigo) and tap water and maintained on a 12h light/dark cycle (lights on at 0630h). Breeders were paired in-house after one week of acclimation to the vivarium. All breeder cages contained nestlets. Sires were removed seven days after breeder pairing to strictly control for cage environment and avoid untimed pregnancies. Phenotyping occurred during spontaneous withdrawal (16h) between 0900h and 1100h and was performed by female experimenters to control for the effect of experimenter sex on rodent behavior (Driel & Talling, 2005; Sorge et al., 2014).

### Tattooing of mice

Pup tails were tattooed (ATS-3 General Rodent Tattoo System, AIMS) on postnatal day (P) 7 following behavioral testing and morning injections for identification and were returned to their home cage.

### Morphine administration in FVB/NJ pups from P1 to P15

We employed a third-trimester approximate mouse model of opioid exposure where injections of either morphine sulfate pentahydrate (10mg/kg, 20ul/g, s.c.; Sigma-Aldrich) or saline (0.9%, 20ul/g, s.c.) were administered twice-daily at 0900h and 1700h from P1 – P14, which is functionally equivalent to the third trimester of human pregnancy (Barr et al., 2011; Byrnes & Vassoler, 2018; Ferrante & Blendy, 2024; Richardson et al., 2006). Additional details on the rationale and procedure are provided in **Supplementary Information**. On P7 and P14, morning injections were administered following phenotyping during spontaneous morphine withdrawal at approximately 1100h.

### Morphine administration in CFW pups from P1 to P14 (Borrelli et al., 2021)

Data analyzed in CFW mice was from a previous study that used a similar morphine treatment regimen, except a higher dose (15 mg/kg, s.c.). In the current study, we used a lower dose in FVB/NJ mice to minimize lethality. See **Supplementary Information** for additional details.

### Recording of ultrasonic vocalizations (USVs)

Individual pups were each placed into a Plexiglass box (43 cm length x 20 cm width x 45 cm height; Lafayette Instruments) within a sound-attenuating chamber (Med Associates). USVs were recorded using the Ultrasound Recording Interface (Avisoft Bioacoustics UltrasoundGate 816H) for 10min (P7) or 15min (P14). Locomotor activity during all USV testing sessions was recorded using infrared cameras and tracked with ANY-maze software (Stoelting).

### Thermal nociception testing on P7 and P14 in FVB/NJ pups

After USV recordings, each pup was removed from the sound-attenuating chamber and placed in a Plexiglass cylinder (diameter, 15 cm; height, 33 cm) on a 52.5°C hot plate (IITC Life Science). On P7, the nociceptive response was defined as the latency for the pup to roll onto its back (Borrelli et al., 2021). On P14, the nociceptive response was defined as the latency to jump, attempt to jump, hind paw lick or attempt to hind paw lick. Pups were removed from the hot plate immediately after observing a nociceptive response or after the 30s cut-off (P7) or the 60s cut-off (P14) if no response was observed. Upon completion of nociceptive testing, each pup was weighed and administered their morning injection and returned to their home cage.

### Supervised ultrasonic vocalization classification

Labeled syllables and accompanying acoustic features were used to train a custom random forest classifier in Python. Additional classifier information is provided in the **Supplementary Information.**

### Bulk RNA-seq of brainstem from morphine-withdrawn pups

The mice used for RNA-seq also underwent phenotyping on P8 and P15 while under the influence of morphine in the a.m. to assess the effect of alleviation of opioid withdrawal on USVs. Additional details are provided in **Supplementary Information**. Maintenance doses resumed for the p.m. injections on P8 and P15. During spontaneous withdrawal (16 h post-morphine) on P16, mice were sacrificed by live, rapid decapitation, and brains were removed from the skull. Brainstem tissue, including the pons, medulla and part of the spinal cord, was immediately collected and stored in RNAlater at 4°C for 5 days, then blotted dry and stored at -80°C for later RNA extractions. Further details on regional brain dissections, RNA extractions, RNA-seq, and analysis are provided in the **Supplementary Information.**

### Real-time quantitative PCR (RT-qPCR) of *Oprk1, Pdyn*, and *Slc6a3* in the brainstem and midbrain on P16

Brainstem and midbrain tissue for RT-qPCR was collected as described in the **Supplementary Information**. cDNA was synthesized using oligo-DT primers from total RNA using a cDNA reverse transcription kit (ThermoFisher Scientific, Cat#4368814). RT-qPCR was performed using PowerUP SYBR Green (ThermoFisher Scientific, Cat#A25741). The primer sequences are listed in the **Supplemental Information**.

### Brainstem gene pathway enrichment analysis

Pathway enrichment analysis was performed on the lists of detectable genes for FVB/NJ and CFW mice. Pathways were obtained from a curated list across multiple different sources (Merico et al., 2010). Background genes were set to only include genes within our data sets. Gene were ranked by the absolute log10 p-values representing differential expression significance multiplied by the sign of the log2FC expression of each gene to preserve directionality. Pathway enrichment analysis was performed on subsequent gene rankings using the fgsea R package with an adaptive multilevel splitting approach and a minimum pathway size set to 15 genes and maximum set to 500 genes (Cérou et al., 2019; Korotkevich et al., 2016). Pathways were separated by those containing upregulated vs. downregulated genes. The collapsePathways command in fgsea was used to remove redundant/similar pathways that were significant (p < 0.05). The top 10 enriched pathways are listed by gene ratio (leading-edge genes/total genes in pathway), with circle size indicating number of leading-edge genes within each pathway and color indicating the pathway *p* value. Pathway clustering was performed using the treeplot function in enrichplot, using a ward.D clustering algorithm with the number of clusters set at 5 (Ward, 1963). The pathway circle size indicates the number of leading-edge genes within each pathway, and colors indicate the pathway *p* value. Figures were created in R studio (https://www.r-project.org/).

### Effect of the kappa opioid receptor antagonist nor-BNI on ultrasonic vocalizations

A separate cohort of FVB/NJ mice was used to test the effects of the KOR antagonist nor-BNI (nor-Binaltorphimine Dihydrochloride; Sigma-Aldrich) on USVs. From P1–P14, injections of either morphine (10mg/kg, s.c) or saline (0.9%, 20 ul/g, s.c.) were administered twice-daily at 0900 h and 1700 h. Nor-BNI has an extremely slow onset of KOR antagonism and a prolonged duration of action at KOR in rodents (Kishioka et al., 2013; Marchette et al., 2021; Munro et al., 2012). Given the absence of pharmacokinetic studies of nor-BNI in neonatal mice, we chose the highest dose (30 mg/kg, s.c.) noted in adult rodent studies (CITE REF) to ensure a complete and prolonged KOR antagonism. Thus nor-BNI (30 mg/kg, s.c.) or saline 0.9%, (20 ul/g, s.c.) was administered at 1300 h on P14, approximately 4 h after the morning morphine or saline injections. Other studies waited between 5 h – 48 h before testing the effects of nor-BNI on rodent behavior, thus we allowed 20h before phenotyping on P15 to ensure we were capturing the KOR-specific effects of nor-BNI on USV emissions (Kelsey et al., 2015; Schlosburg et al., 2013; Walker et al., 2011). USVs were recorded at 0900h for 15 min.

### Effect of the kappa opioid receptor agonist U50,488h on ultrasonic vocalizations

A separate cohort of morphine-naïve FVB/NJ mice was used to determine whether administration of the KOR agonist U50,488h ((±)-U-50488h hydrochloride; Fisher Scientific) would be sufficient to induce USVs mirroring the opioid withdrawal profile. From P1– P14, pups were injected with saline (0.9%, 20 ul/g, s.c.) twice daily at 0900 h and 1700 h to replicate the NOWS injection regimen. On P10 and P14, pups were injected with either U50,488h (0.625 mg/kg, s.c.) or saline (s.c.) and then returned to their home cage. We allowed 10 min before recording to ensure the effects of U50,488h on USV emission within previously reported time frames (Carden et al., 1994; Huang et al., 2020). USVs were recorded at 0900 h for 15 min. Pups continued to receive saline injections following P10 testing until testing for the behavioral effects of U50,488h again on P14, which is when we normally record USVs. **See Supplemental Information for U50,488h dose-response pilot and P10 justification.**

### Statistical analysis

Analysis was performed in R (https://www.r-project.org/). All data are presented as the mean ± standard error of the mean (SEM), and *p*<0.05 was considered significant. Figures were created in GraphPad Prism (https://graphpad.com/). Body weight, temperature, USV locomotion, and USVs over time were analyzed using linear mixed models with Saline Treatment and Female Sex as the reference variables and Pup as a random effect (repeated measures). Sex was removed from the model if there were no interactions. Significant interactions of interest were followed up with least-square means (predicted marginal means) and Tukey’s Honestly Significant Difference (HSD) tests. All other data were analyzed using linear models with Saline Treatment and Female Sex as the reference variables. For nor-BNI experiments, we were mostly interested in the effect of nor-BNI on USV emissions in morphine-treated females, thus Morphine Treatment (P1 – P14 treatment), Saline Treatment (pre-treatment) and Female sex were used as reference variables. Based on the RNA-seq results, we had a specific hypothesis for the direction of fold-change in gene expression, therefore, for RT-qPCR analysis, one-tailed t-tests were performed. For *Slc6a3*, one sample was identified as an outlier (ROUT (Q = 1%)) and was removed from the analysis.

## RESULTS

### Morphine exposure from P1–P14 is sufficient to induce multiple signs of spontaneous opioid withdrawal in neonatal FVB/NJ pups

An experimental timeline is provided in **Fig.1a**. Reduced body weight and poor body temperature regulation are commonly observed in infants diagnosed with NOWS (Finnegan et al., 1975; Hudak et al., 2012; Jansson & Velez, 2012; Patrick et al., 2020). Twice-daily morphine injections (10 mg/kg, s.c.) from P1–P14 significantly reduced body weight (**Fig.1b**) and temperature during withdrawal (**Fig.1c**). Hyperalgesia is common in opioid withdrawal (Corder et al., 2018). Morphine-withdrawn pups displayed thermal hyperalgesia on P7, as indicated by a reduced latency to elicit a nociceptive response on the hot plate (**Fig.1d)**. In examining hot plate velocity on P7 during the time interval of exposure, there was no effect of morphine treatment, indicating that reduced locomotor activity did not prolong hot plate exposure and thus did not confound the group difference (**Fig.1e**). On P14, morphine-withdrawn pups displayed reduced hot plate latency (**Fig.1f**) and increased hot plate velocity (**Fig.1g)**. Because increased velocity decreases paw exposure to the hot plate, we can again conclude that increased locomotor activity did not confound hyperalgesia in morphine-withdrawn pups.

**Fig. 1.**
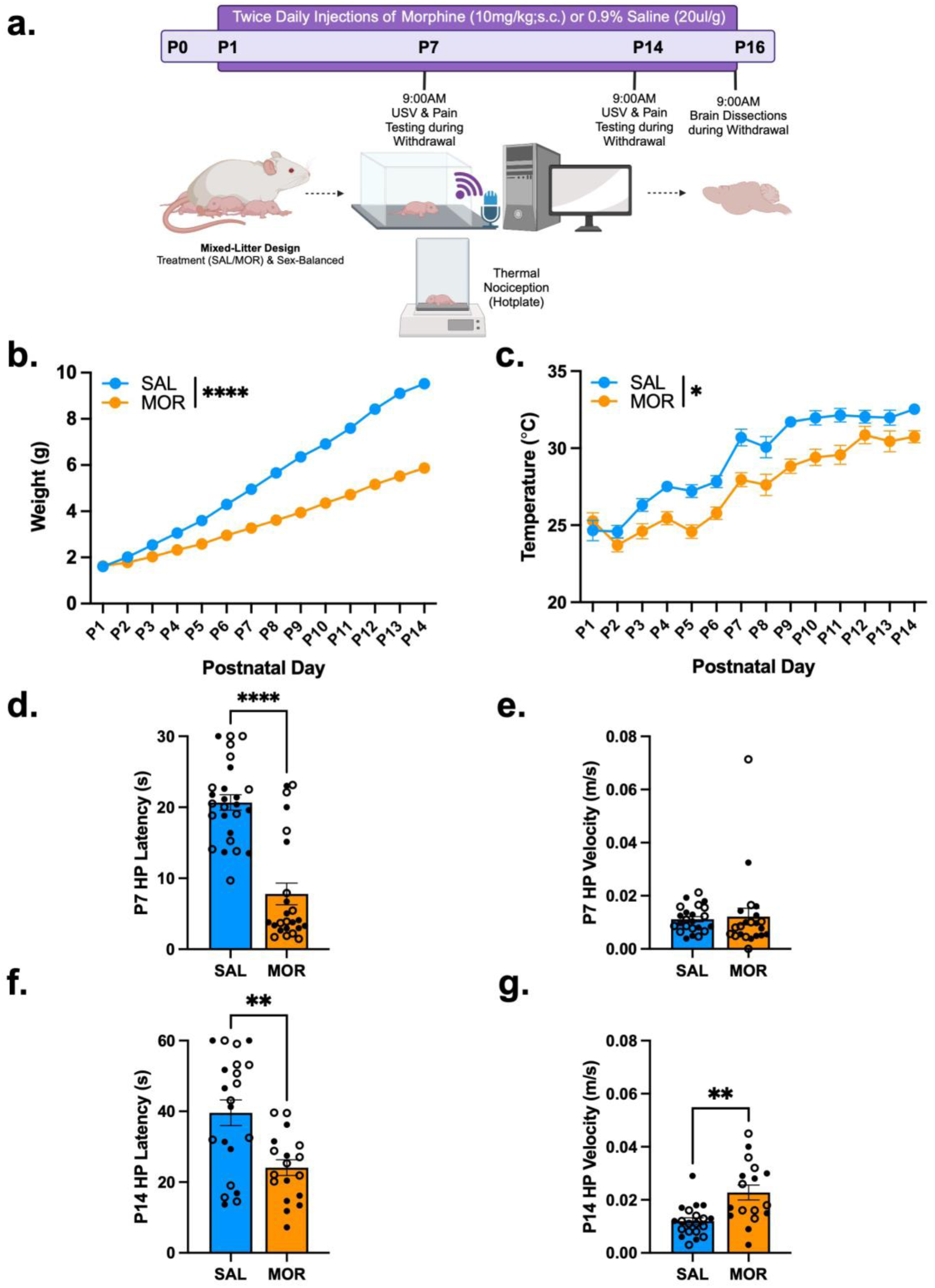
Morphine exposure from P1–P14 is sufficient to induce opioid withdrawal traits in neonatal FVB/NJ pups. Data are plotted as the mean ± SEM. Saline = blue lines/bars; Morphine = orange lines/bars. Closed circles = Females; Open circles = males. **(a) Experimental Timeline. (b) Body Weight**: The effect of Morphine Treatment on body weight was dependent on Postnatal day (β = - 0.30, SE = 0.0055, t(611) = -54.38, \*\*\*\**p* < 0.0001), where morphine-treated pups weighed significantly less than saline-treated pups from P3 (\*\**p* = 0.0013) – P14 (P4 – P14: \*\*\*\**p* < 0.0001). **(c) Temperature**: The effect of Morphine Treatment on body temperature was dependent on Postnatal day (β = -0.087, SE = 0.041, t(611) = -2.12, \**p* = 0.035), where morphine-treated pups displayed hypothermia compared to saline-treated pups from P3–P11 and P13 – P14 (all \**p* ≤ 0.026). SAL, n = 25 (11F, 14M); MOR, n = 22 (11F, 11M). **(d) P7 Hot Plate Latency**: There was no effect of Sex (β = 0.49, SE = 2.64, t(46) = 0.19, *p* = 0.85) or a Morphine Treatment x Sex interaction (β = 0.26, SE = 3.81, t(46) = 0.068, *p* = 0.95). The simplified model revealed that Morphine Treatment was associated with a decrease in hot plate latency compared to saline controls (β = -12.88, SE = 1.86, t(48) = -6.91, \*\*\*\**p* < 0.0001). **(e) P7 Hot Plate Velocity**: There was no effect of Sex (β = 0.00037, SE = 0.005, t(39) = 0.076, *p* = 0.94) or a Morphine Treatment x Sex interaction (β =0.0030, SE = 0.0069, t(39) = 0.43, *p* = 0.67). The simplified model revealed that Morphine Treatment had no effect on hot plate velocity (β = 0.001, SE = 0.0034, t(41) = 0.27, *p* = 0.79). SAL, n = 23–26 (11–12F, 12–14M); MOR, n = 22–24 (12–13F, 10–11M). **(f) P14 Hot Plate Latency**: There was no effect of Sex (β = 1.089, SE = 6.085, t(35) = 0.18, *p* = 0.86) or a Morphine Treatment x Sex interaction (β = 7.33, SE = 8.91, t(35) = 0.82, *p* = 0.67). The simplified model revealed that Morphine Treatment was associated with decreased hot plate latency compared to saline controls (β = -15.50, SE = 4.42, t(37) = -3.51, \*\**p* = 0.0012). **(g) P14 Hot Plate Velocity**: There was no effect of Sex (β = -0.0047, SE = 0.0041, t(33) = -1.15, *p* = 0.23) or a Morphine Treatment x Sex interaction (β = 0.0094, SE = 0.0059, t(33) = 1.58, *p* = 0.12). The simplified model revealed that Morphine Treatment was associated with increased hot plate velocity compared to saline controls (β = 0.011, SE = 0.0030, t(35) = 3.67, \*\*\**p* = 0.00081). SAL, n = 20–21 (8–9F, 12M); MOR, n = 17–18 (9F, 8–9M)

### USV locomotor activity of FVB/NJ pups during spontaneous morphine withdrawal on P7 and P14

On P7, morphine-withdrawn pups traveled a greater distance during the first 2–5min of the USV recording session compared to saline pups (**Fig.2a**). For the entire 10 min, morphine-withdrawn pups traveled a greater distance versus saline-treated pups (**Fig.2b**). There was no effect of morphine withdrawal on P7 average velocity (**Fig.2c**). On P14, there was no significant difference in total distance traveled over 15 min (**Fig.2d**) nor average velocity between treatment groups (**Fig.2e**).

**Fig. 2.**
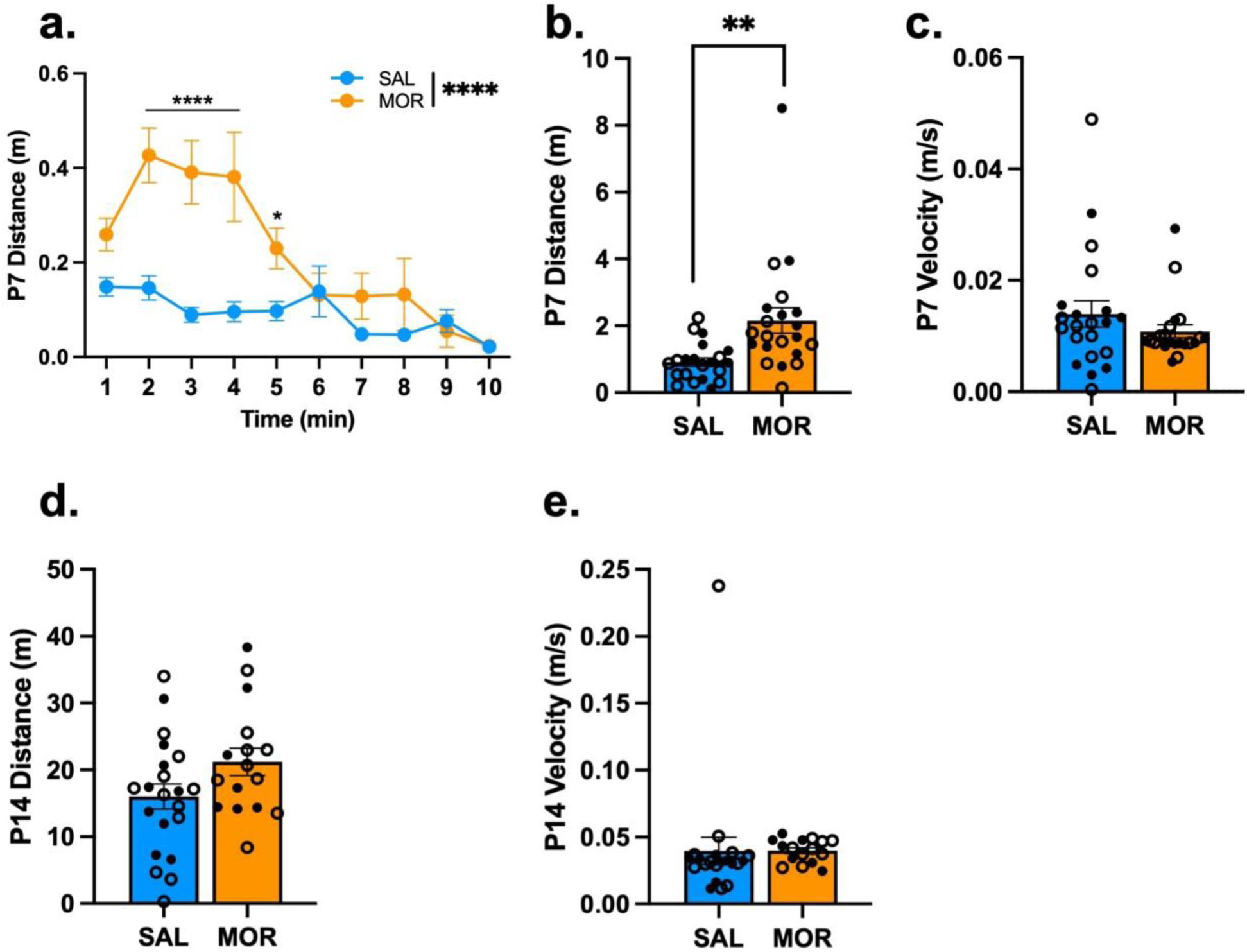
Locomotor activity during USV assessment of FVB/NJ pups undergoing spontaneous morphine withdrawal on P7 and P14. Data are plotted as the mean ± SEM. Saline = blue lines/bars; Morphine = orange lines/bars. Closed circles = Females; Open circles = males. **(a) P7 Distance**: The effect of Morphine Treatment on USV distance was dependent on Time (β = -0.030, SE = 0.0056, t(378) = -5.34, \*\*\*\**p* < 0.0001), where morphine-treated pups traveled a greater distance than saline controls from 2 – 5 min of the recording session (all \**p*^adj^ ≤ 0.031). **(b) P7 Total Distance:** There was no effect of Sex (β = -0.11, SE = 0.58, t(38) = -0.20, *p* = 0.85) or a Morphine Treatment x Sex interaction (β = -0.73, SE = 0.80, t(38) = -0.92, *p* = 0.36). The simplified model revealed that Morphine-treated pups traveled a greater distance during USV recordings compared to saline-treated pups (β = 1.24, SE = 0.40, t(40) = 3.12, \*\**p* = 0.0033). **(c) P7 Average Velocity:** There was no effect of Sex (β = 0.0022, SE = 0.0040, t(38) = 0.59, *p* = 0.56) or a Morphine Treatment x Sex interaction (β = -0.0020, SE = 0.0055, t(38) = -0.37, *p* = 0.72). The simplified model revealed that Morphine Treatment had no effect on USV velocity (β = -0.0031, SE = 0.0027, t(40) = -1.17, *p* = 0.25). SAL, n = 21 (9F, 12M); MOR, n = 21 (11F, 10M). **(d) P14 Distance:** There was no Morphine Treatment x Time interaction (β = 0.0029, SE = 0.012, t(518) = 0.24, *p* = 0.81), so data was collapsed across the 15 min recording session. There was no effect of Sex (β = -0.93, SE = 3.84, t(33) = -2.42, *p* = 0.81) or a Morphine Treatment x Sex interaction (β = -0.26, SE = 5.83, t(33) = -0.045, *p* = 0.96). The simplified model revealed that Morphine Treatment had no effect on the total distance traveled (β = 5.20, SE = 2.81, t(35) = 1.85, *p* = 0.073). **(e) P14 Average Velocity:** There was no effect of Sex (β = 0.019, SE = 0.016, t(33) = 1.20, *p* = 0.24) or a Morphine Treatment x Sex interaction (β =-0.020, SE = 0.024, t(33) = -0.82, *p* = 0.42). The simplified model revealed that Morphine Treatment had no effect on USV velocity (β = 0.00020, SE = 0.012, t(35) = 0.017, *p* = 0.99). SAL, n = 21 (9F, 12M); MOR, n = 13 (7F, 9M)

### USV syllable profile of FVB/NJ pups during spontaneous morphine withdrawal on P14

Neonatal USVs communicate negative internal states. Thus, we measured USVs as an indicator of enhanced distress during spontaneous morphine withdrawal (Branchi et al., 2001; D’Amato et al., 2005; Ehret, 2005). On P7, there was a significant Morphine Treatment x Time interaction (**Fig.S1a**); however, there was no overall effect of Morphine Treatment on the total number of USVs over 10 min (**Fig.S1b).** On P14, there was a time-dependent increase in vocalizations in morphine-withdrawn female pups (**Fig.3a**). Overall, Morphine-withdrawn females emitted significantly more USVs compared to saline-treated females, whereas there was no Treatment effect in males (**Fig.3b**).

**Fig. 3.**
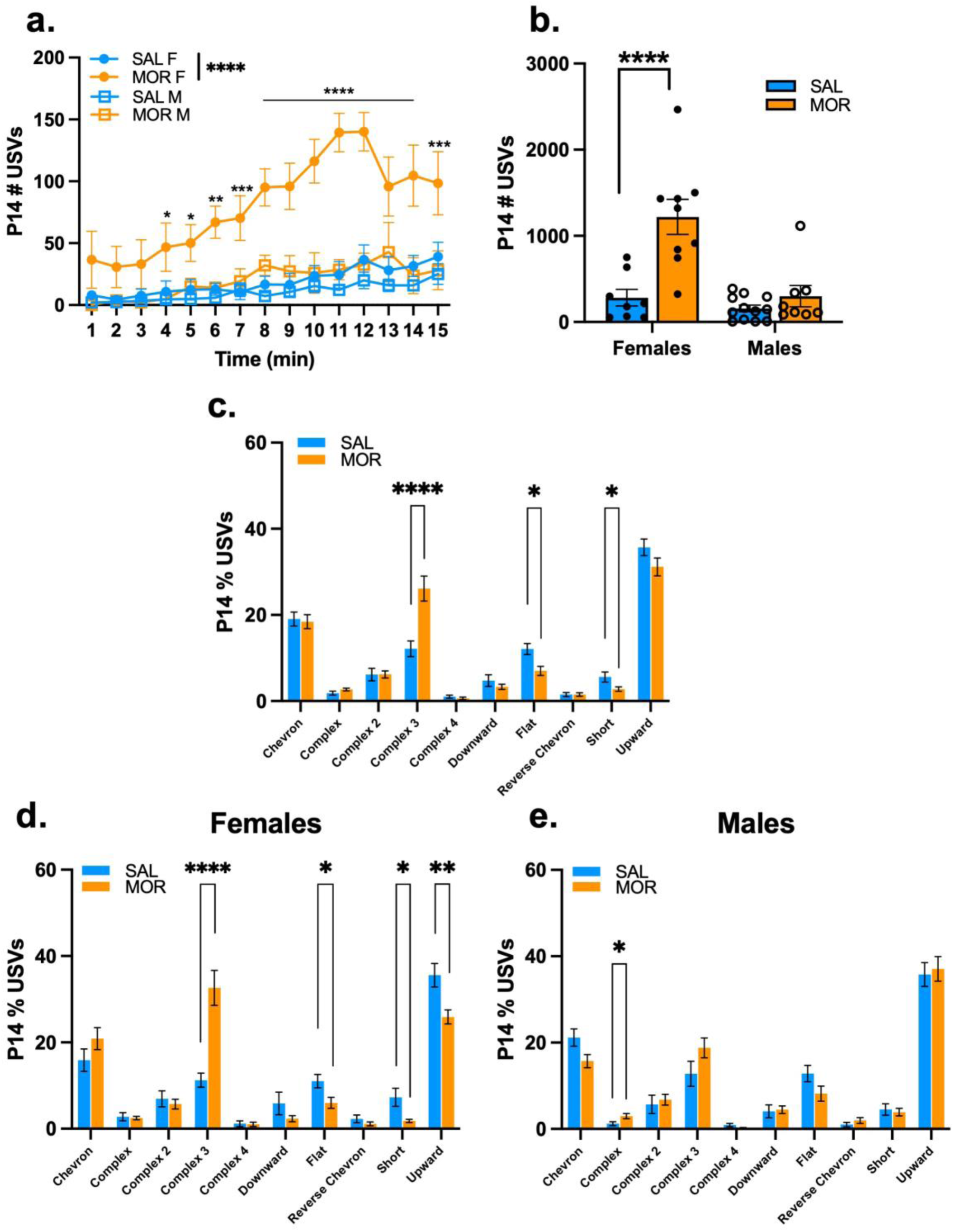
USV profiles of FVB/NJ pups during spontaneous morphine withdrawal on P14. Data are plotted as the mean ± SEM. Saline = blue lines/bars; Morphine = orange lines/bars. Closed circles = Females; Open circles = males. **(a) P14 USV Emissions:** There was a Morphine Treatment x Sex x Time interaction (β = -3.60, SE = 1.03, t(518) = -3.49, \*\*\**p* = 0.0052), where morphine females vocalized more than saline females over time, specifically during the 4 – 15 min intervals (all \**p*^adj^ ≤ 0.035) . **(b) P14 Total USV Emissions**: The effect of Morphine Treatment on USV emission was dependent on Sex (β = 919.6, SE = 180.7, t(33) = 5.088, \*\*\*\**p* < 0.0001). Specifically, morphine-treated females vocalized more than saline females (β = 937.0, SE = 181, t(33) = 5.19, \*\*\*\**p* < 0.0001). **(c) P14 Syllable Profile:** Morphine Treatment was associated with an increase in the percentage of Complex 3 syllables emitted (β = 0.14, SE = 0.033, t(35) = 4.19, \*\*\**p* = 0.00018), and a decrease in the percentage of Flat (β = -0.051, SE = 0.017, t(35) = -3.0, \*\**p* = 0.0051), and Short (β = -0.030, SE = 0.014, t(35) = -2.08, \**p* = 0.045) syllables emitted compared to saline controls. There were Morphine Treatment x Sex interactions for Complex 3 (β = 0.15, SE = 0.060, t(33) = 2.54, \**p* = 0.016) and Upward (β = 0.14, SE = 0.044, t(33) = 3.13, \*\**p* = 0.0036) syllables, so we investigated the syllable profile for each sex separately. **(d) Female P14 Syllable Profile:** Morphine Treatment was associated with an increase in the percentage of Complex 3 syllables (β = 0.21, SE = 0.046, t(15) = 4.8, \*\*\**p* = 0.00030) and a decrease in Flat (β = -0.051, SE = 0.020, t(15) = -2.55, \**p* = 0.022), Short (β = -0.055, SE = 0.020, t(15) = -2.69, \**p* = 0.017), and Upward syllables (β = -0.097, SE = 0.031, t(15) = -3.13, \*\**p* = 0.0069). **(e) Male P14 Syllable Profile:** Morphine Treatment was associated with an increase in the percentage of Complex syllables (β = 0.017, SE = 0.0072, t(18) = 2.41, \**p* = 0.027). SAL, n = 20 (8F, 12M); MOR, n = 17 (9F, 8M)

USVs are classified into syllable types based on their acoustic features (frequency, duration) (Grimsley et al., 2011; Heckman et al., 2016; Portfors, 2007). We used a random forest model to classify USV syllables based on these known classifications (**Table.S1)** during neonatal morphine withdrawal. On P7, morphine-withdrawn pups emitted fewer Complex 3 syllables than saline pups (**Fig.S1c**). On P14, morphine-withdrawn pups showed the opposite, namely a robust *increase* in the percentage of Complex 3 syllables (**Fig.3c**). The increase in Complex 3 syllables was completely driven by morphine-withdrawn females, as there was no treatment difference in males (**Fig.3d,e**). Morphine-withdrawn females also showed a decrease in the percentage of Flat, Short, and Upward syllables, whereas males only showed an increase in the percentage of Complex syllables (**Fig.3d,e**).

The identification of increased Complex 3 syllables during spontaneous opioid withdrawal was a robust and novel, female-selective observation. To determine whether increased Complex 3 emissions in females generalize to another genetic background, we also examined USV syllable subtypes in morphine-withdrawn outbred CFW stock mice from a historical dataset (Borrelli et al., 2021) comprising a nearly identical morphine regimen, except with a higher twice daily dose (15 mg/kg, s.c.; see **Supplementary Information**). On P7, treatment had no effect on the USV syllable profile (**Fig.S2a**). On P14, the USV profile in morphine-withdrawn CFW pups showed a similar increase in the percentage of Complex 3 syllables (**Fig.S2b**), except in this case, both sexes showed this effect (perhaps due to the higher historical morphine dose that was used in the CFW study (Borrelli et al., 2021)). Additional similarities between morphine-withdrawn CFW and FVB pups included a decrease in the percentage of Flat and Upward syllables on P14 (**Fig.S2b**).

### Brainstem transcriptomic analysis identified an upregulation of the kappa opioid receptor

In a subset of FVB/NJ mice, we administered an a.m. maintenance dose of morphine (10 mg/kg) prior to USV recording on P8 and P15 to evaluate the semi-acute effects of morphine exposure on the USV syllable profile (**Fig.S3**). We collected brainstem tissue during spontaneous withdrawal on P16 (16 h post-final morphine injection) to evaluate the effects of chronic morphine exposure and withdrawal on gene expression (**Fig.4a**). A schematic of regional brain dissections is provided in **Fig.S4**. Transcriptomic analysis via RNA-seq revealed a significant upregulation of *Oprk1* (log_2_FC=0.36, \*\**p*=0.0049), which codes for the kappa opioid receptor (**KOR**) and *Slc6a3* (dopamine transporter (DAT); log_2_FC=3.69, \*\**p*=0.0067) during spontaneous morphine withdrawal (**Table.S2**). We also observed a downregulation of genes associated with myelination, such as *Mbp* (myelin basic protein; log_2_FC= 0.41, \*\*\*\**p*<0.0001) and *Plp1* (proteolipid protein 1; log_2_FC=-0.38, \*\*\*\**p*<0.00032) (**Table.S3**).

**Fig. 4.**
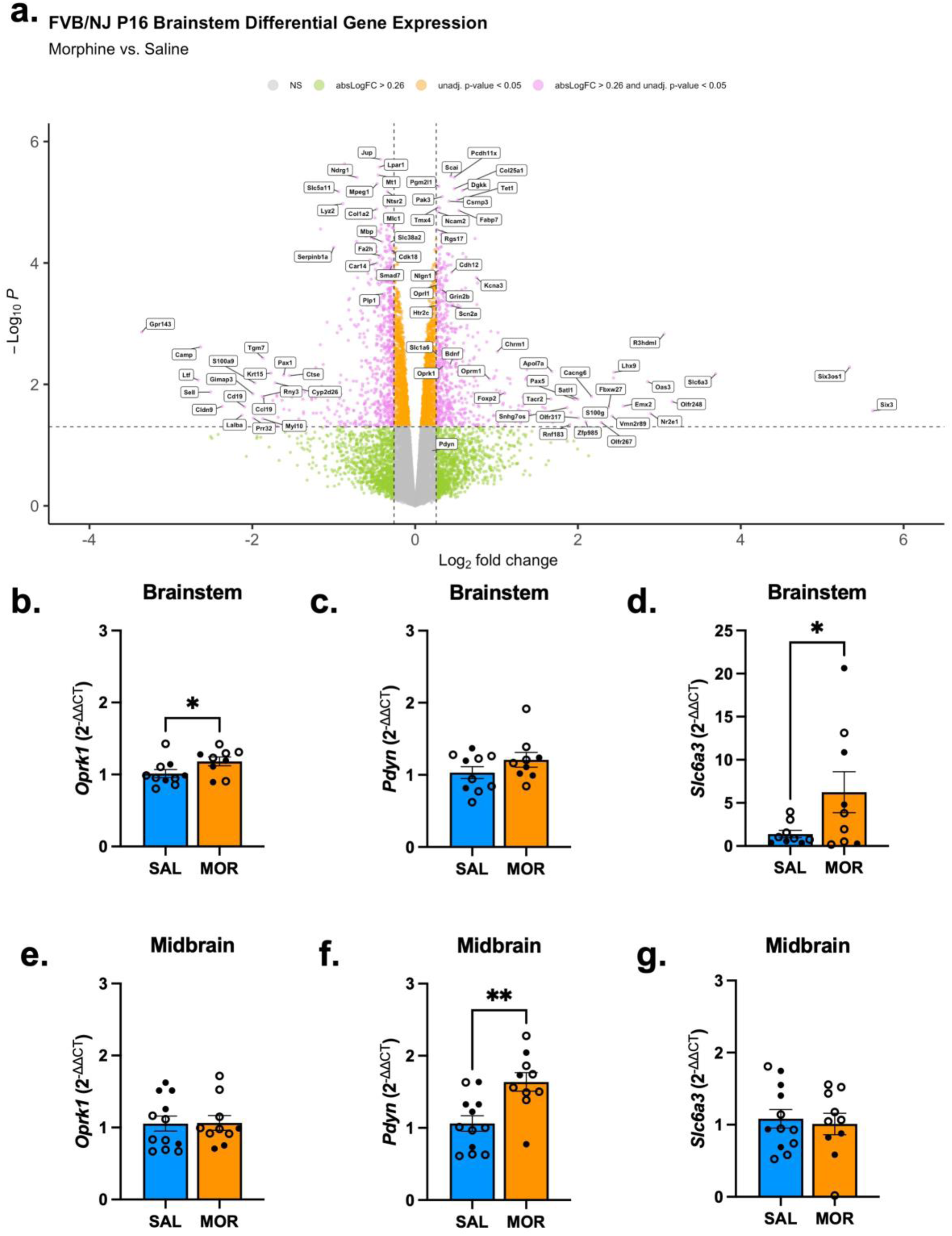
Differentially expressed genes in the brainstem and midbrain during spontaneous morphine withdrawal on P16. **(a) Brainstem RNAseq:** Data reflects the effect of morphine exposure on gene expression relative to saline pups (sex-collapsed; Saline, n = 4 (2F, 2M); Morphine, n = 4 (2F, 2M). Log_2_ Fold Change (FC) represents gene expression, and Log_10_ *p* values (unadjusted) reflect significance. To avoid crowding, we labeled genes with an absolute (abs) log_2_FC ≥ 1.5 and/or Log_10_P ≥ 4.0. Dots are individual genes. Grey = absolute (abs) log_2_FC ≥ 0.26; non-significant (unadjusted *p* < 0.05; NS). Green = abslog_2_FC ≥ 0.26, non-significant; Orange = abslog_2_FC ≤ 0.26, significant (unadjusted *p* < 0.05); Pink = abslog_2_FC ≥ 0.26, significant (unadjusted *p* < 0.05). RT-qPCR: Data are plotted as the mean ± SEM. Saline = blue bars; Morphine = orange bars. Closed circles = Females (SAL, n = 3–4; MOR, n = 3–4); Open circles = males (SAL, n = 6–8; MOR, n = 5–7). **(b) Brainstem *Oprk1***: Morphine Treatment was associated with upregulation of *Oprk1* (t(17) = 2.06, \**p* = 0.0028). **(c) Brainstem *Pdyn***: There was no effect of Morphine Treatment on *Pdyn* expression (t(17) = 1.35, *p* = 0.097). **(d) Brainstem *Slc6a3***: Morphine Treatment was associated with upregulation of *Sl6a3* (t(16) = 2.00, \**p* = 0.031). SAL, n = 10 (3F, 6 – 7M); MOR, n = 9 (4F, 5M). **(e) Midbrain *Oprk1***: Morphine Treatment had no effect on *Oprk1* expression (t(20) = 0.066, *p* = 0.47). **(f) Midbrain *Pdyn***: Morphine Treatment was associated with upregulation of *Pdyn* (t(20) = 3.47, \**p* = 0.0012). **(g) Midbrain *Slc6a3***: There was no effect of Morphine Treatment on *Slc6a3* expression (t(20) = 0.37, *p* = 0.64)

We performed RT-qPCR on brainstem and midbrain tissue to validate these mRNA transcripts with a larger sample size. Consistent with our brainstem RNA-seq results, we observed withdrawal-induced upregulation of *Oprk1* (**Fig.4b**), no effect on *Pdyn* expression (**Fig.4c**), and upregulation of *Slc6a3* (**Fig.4d**). Interestingly, for the midbrain, we did not observe a withdrawal effect on *Oprk1* expression (**Fig.4e**). However, we found upregulation of *Pdyn* (**Fig.4f**). There was no effect of withdrawal on *Slc6a3* (**Fig.4g**) expression.

Enrichment analysis of upregulated genes in the brainstem revealed processes associated with synaptic organization, structure, and activity (**Fig.5a**), and enriched pathways of downregulated genes associated with extracellular matrix organization and glial development (**Fig.5b**).

**Fig. 5.**
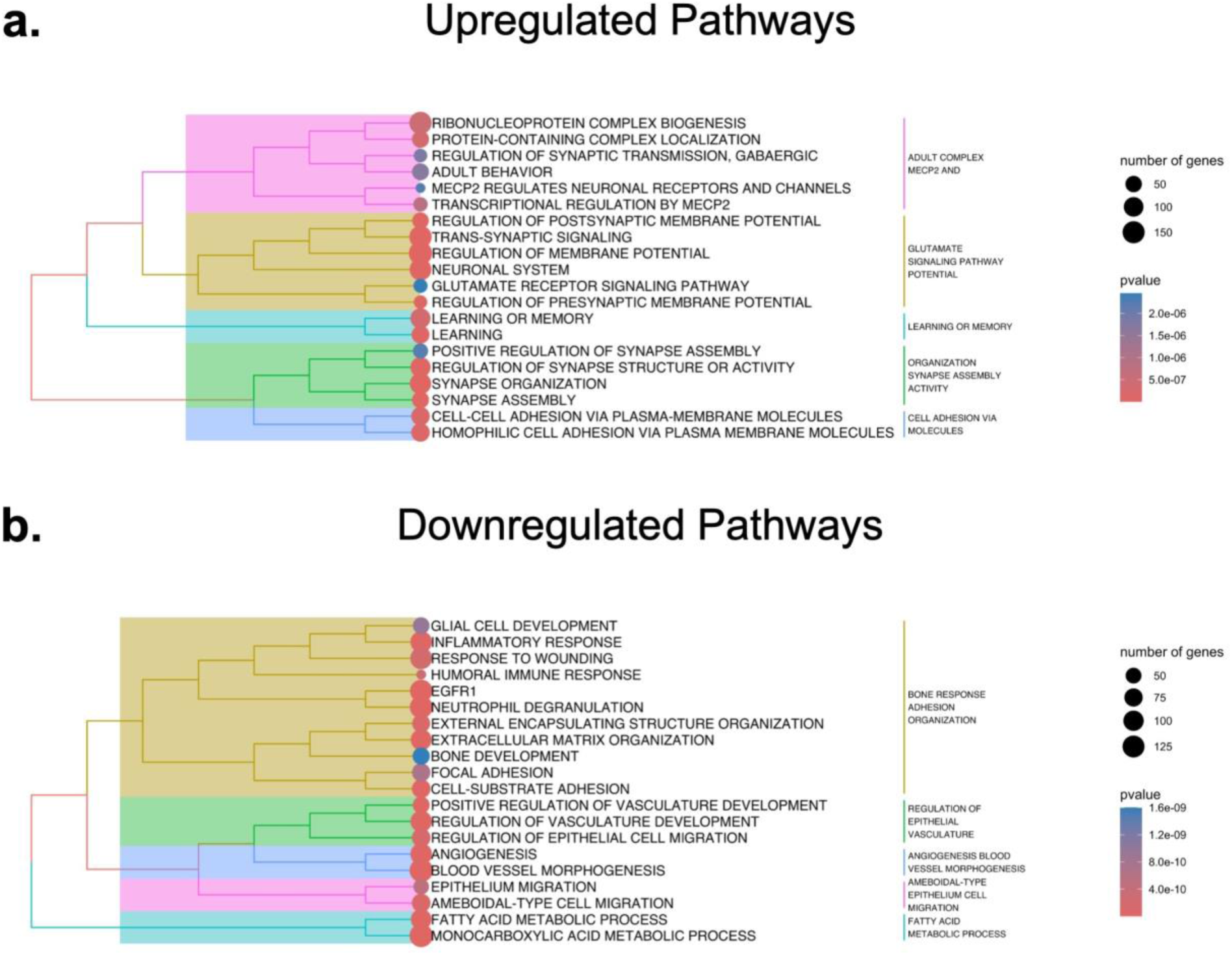
Pathway enrichment analysis of differentially expressed genes in the brainstem during spontaneous morphine withdrawal on P16. Colors (pink, yellow, teal, green, blue) correspond to pathway names. The size of the circles reflects the number of genes in the pathway (the larger the circle, the more genes are associated with the pathway). The color of the circles reflects the significance (decreasing *p* value (more significant) from blue to red). **(a) Upregulated Pathways**: clustering of the top 20 enriched pathways consisting of upregulated genes in FVB/NJ mice. **(b) Downregulated Pathways**: Clustering of the top 20 enriched pathways consisting of downregulated genes in FVB/NJ mice. Clustering was determined by calculating Jaccard similarity coefficients between pathways, with the number of clusters set to 5

### The kappa opioid receptor antagonist nor-BNI reduced the number of USVs in females during spontaneous morphine withdrawal

KOR activation contributes to the dysphoric aspects of withdrawal, such as hyperirritability, anhedonia, and anxiety (Land et al., 2008; Margolis & Karkhanis, 2019; Shippenberg et al., 2007). Furthermore, sex differences in the dynorphin/KOR system on motivated behaviors in the context of drug addiction are well-documented in rodents and humans (Chartoff & Mavrikaki, 2015). Therefore, we aimed to determine whether the dynorphin/KOR system contributes to increased USVs and Complex 3 emissions during opioid withdrawal. On P15, 20 h post-administration of the KOR antagonist, nor-BNI, there was a significant decrease in USVs in morphine-withdrawn females but no effect of nor-BNI on USVs in males (**Fig.6a**). Furthermore, there was a reduction in USVs over time in morphine-treated females, but not males, pre-treated with nor-BNI compared to morphine controls (**Fig.6b**). These results identify a KOR contribution to the female-specific enhancement of USVs in morphine-withdrawn pups. However, KOR activation was not specific for a particular syllable type, as nor-BNI had no effect on the overall USV syllable profile (**Fig.6c**).

**Fig. 6.**
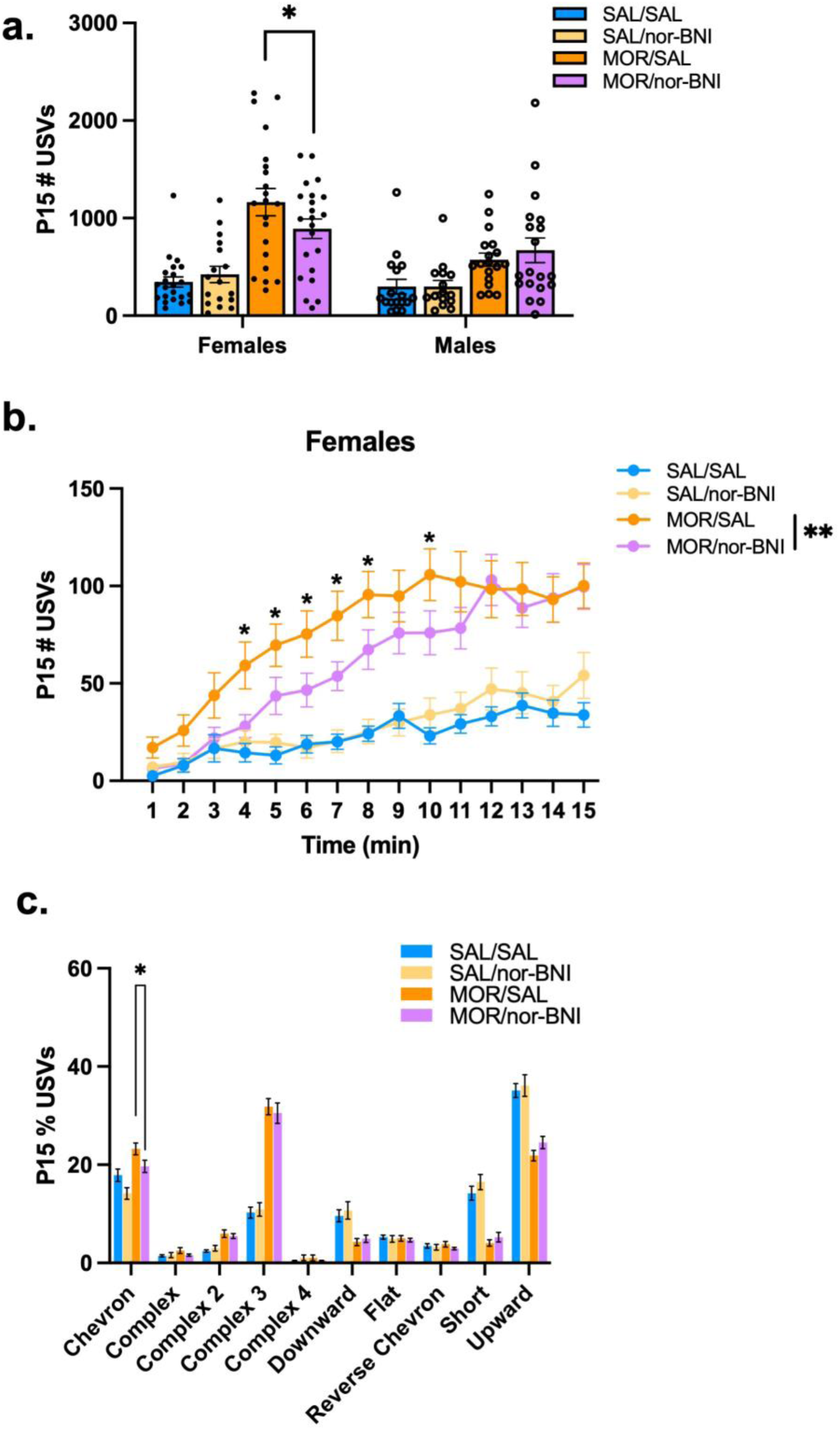
The kappa opioid receptor antagonist nor-BNI reduces USV emissions in FVB/NJ female pups during spontaneous withdrawal on P15. Data are plotted as the mean ± SEM. Saline animals pre-treated with saline = blue lines/bars; Saline animals pre-treated with nor-BNI = yellow lines/bars; Morphine animals pre-treated with saline = orange lines/bars; Morphine animals pre-treated with nor-BNI = purple lines/bars. Closed circles = Females; Open circles = males. **(a) P15 Total USV Emissions**: There was a Morphine Treatment x nor-BNI Treatment x Sex interaction (β= -874.5, SE = 421.2, t(143) = -2.076, \**p* = 0.040), where Morphine-treated females pre-treated with nor-BNI (MOR/norBNI) emitted fewer USVs compared to morphine-treated females pre-treated with saline (MOR/SAL) (β = -272.35, SE = 127.11, t(143) = -2,143 \**p* = 0.034). **(b) Female P15 USV Emissions**: Given the three-way interaction, we analyzed females and males separately across time to simplify the linear mixed model. There was a time-dependent decrease in USVs in morphine-treated females pre-treated with nor-BNI (MOR/norBNI) compared to morphine-treated females pre-treated with saline (MOR/SAL) (β = -1.42, SE = 0.52, t(1148) = -2.72, \*\**p* = 0.0068), specifically during the 4–8 and 10 min intervals (all *\*p*^adj^ ≤ 0.037). **(c) P15 Syllable Profile:** There was no effect of Sex on any Syllable emitted (all *p* ≥ 0.20), so Sex was removed from the models. Morphine pups pre-treated with nor-BNI emitted fewer Chevron syllables compared to morphine-treated pups pre-treated with saline (β = -0.036, SE = 0.017, t(147) = -2.065 \**p* = 0.041). SAL/SAL, n = 40 (22F, 18M); SAL/norBNI, n = 33 (17F, 16M); MOR/SAL, n = 39 (21F, 18M); MOR/norBNI, n = 39 (22F, 17M)

### The kappa opioid receptor agonist U50,488h is sufficient to induce an increase in USVs

Previous studies reported that the administration of KOR agonists, such as U50,488h, is aversive and can induce anhedonia and decreased reward sensitivity (Conway et al., 2019; Huang et al., 2020; Vonvoigtlander et al., 1983). Here, we wished to determine whether KOR activation with the KOR agonist U50,488h was sufficient to increase USVs and induce a syllable profile similar to morphine-withdrawn pups, particularly in females. To conduct this experiment, we considered a postnatal day where saline-treated controls would show sufficient baseline USVs to detect either an increase or decrease in USVs. In neonates, the rate of USVs peaks around P7–P10 and declines during the second postnatal week (Elwood & Keeling, 1982). Thus, we tested the KOR agonist U50,488h on P10 and then again on P14, when we normally assess USVs (**Fig.3**). Based on a pilot study, the U50,488h dose of 0.625mg/kg was chosen because it induced an increase in USVs without affecting locomotor activity (**Fig.S5a,b**). On P10, U50,488h increased USVs relative to saline controls during the first 10 min of the recording session (**Fig.7a**). Additionally, in considering the entire 15 min, U50,488h increased overall USVs in both sexes (**Fig.7b**). In contrast to our original hypothesis (although in line with the null effect of the KOR antagonist in morphine-withdrawn mice), there was no effect of KOR activation on the percentage of Complex 3 emissions on P10 in either sex (**Fig.7c**). In considering other syllable types, there was a significant decrease in the percentage of Downward and Short emissions (**Fig.7c**). On P14, there was a time-dependent effect of U50,488h, whereby KOR activation increased USVs during the 5–9 min of the recording session (**Fig.7d**). Furthermore, on P14, U50,488h increased USVs in females, but not in males (**Fig.7e**). In contrast to our original hypothesis, U50,488h *decreased* the percentage of Complex 3 syllables in both sexes (**Fig.7f**). U50,488h also decreased the emission of other multi-component syllables, including Complex 2 and Complex 4 in both sexes (**Fig.7f**).

**Fig. 7.**
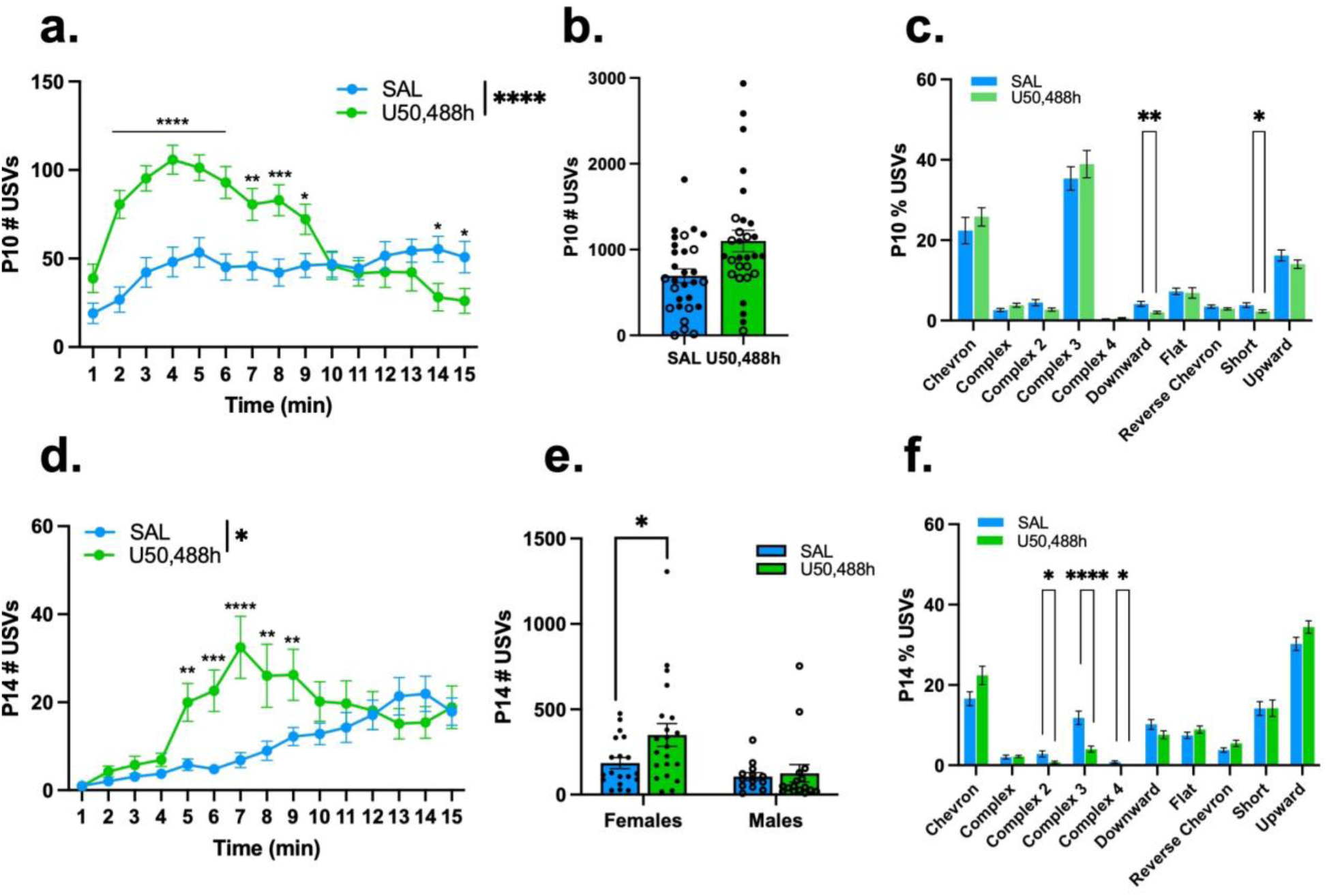
The kappa-opioid receptor agonist U50,488h, increases USV emission in morphine-naïve FVB/NJ pups on P10 and P14. Data are plotted as the mean ± SEM. Saline = blue lines/bars; U50,488h = green lines/bars. Closed circles = Females; Open circles = males. **(a) P10 USV Emission**: The effect of U50,488h Treatment depended on Time (β = -5.89, SE = 0.52, t(812) = -11.42, \*\*\*\**p* < 0.0001). U50,488h-treated pups vocalized more than saline pups during the 2–9min intervals (all \**p*^adj^ ≤ 0.020), and less than saline pups during the 14–15min intervals (both \**p*^adj^ ≤ 0.028). **(b) P10 Total USV Emission**: There was no effect of Sex (β = -167.71, SE = 205.43, t(54) = -1.34, *p* = 0.19) or a U50,488h Treatment x Sex interaction (β =-34.18, SE = 288.38, t(54) = -0.12, *p* = 0.91). The simplified model revealed that U50,488h Treatment was associated with increased USV emission compared to saline controls (β= 406.2, SE= 148.9, t(56) = 2.73, \*\**p* = 0.0085). **(c) P10 Syllable Profile:** There was no effect of Sex on the percentage of syllables emitted (all *p* ≥ 0.42) or a U50,488h Treatment x Sex interaction (all *p* ≥ 0.42). The simplified model revealed that U50,488h Treatment was associated with a decrease in the percentage of Downward syllables (β = -0.021, SE = 0.0070, t(56) = -3.04, \*\**p* = 0.0036) and Short syllables (β = -0.015, SE = 0.0072, t(56) = -2.11, \**p* = 0.039) compared to saline controls. **(d) P14 USV Emission:** The effect of U50,488h on USV emission depended on Time (β = -0.56, SE = 0.025, t(966) = -2.25, \**p* = 0.025), where U50,488h-treated pups vocalized more than saline pups during the 5–9min intervals (all \*\**p*^adj^ ≤ 0.0087). **(e) P14 Total USV Emission:** There was a U50,488h Treatment x Sex interaction (β = 202.10, SE = 97.93, t(65) = 2.06, \**p* = 0.043), where U50,488h Treatment was associated with increased USV emission in females compared to saline-treated females (β = 164.85, SE = 67.34, t(65) = 2.45, \**p* = 0.017). **(f) P14 Syllable Profile**: There was no effect of Sex on the percentage of syllables emitted (all *p* ≥ 0.12) or a U50,488h Treatment x Sex interaction (all *p* ≥ 0.15). The simplified model revealed that U50,488h Treatment was associated with a decrease in Complex 2 (β = -0.021, SE = 0.0082, t(67) = -2.57, \**p* = 0.012), Complex 3 (β = -0.078, SE = 0.018, t(67) = -4.43, \*\*\*\**p* < 0.0001), and Complex 4 (β = -0.0080, SE = 0.0028, t(67) = -2.91, \*\**p* = 0.0049) syllables. SAL, n = 29–32 (18–19F, 11–13M); U50,488h, n = 31–37 (17–21F, 12–16M)

## DISCUSSION

Consistent with previous studies, twice-daily injections of morphine from P1 to P14 was sufficient to induce withdrawal-associated model traits in FVB/NJ pups on P14, including low body weight, hypothermia, hyperalgesia, and increased USVs on P14 (Borrelli et al., 2021; Carden et al., 1991; S. A. Robinson et al., 2020). Neonatal mice emit USVs exclusively in isolation as a distress signal to communicate negative internal states and promote maternal attention (Branchi et al., 2001; D’Amato et al., 2005; Dirks et al., 2002; Ehret, 2005). Rodent studies observed increased USVs during opioid withdrawal in neonatal pups (mice and rats) and adult rats (Barr & Wang, 1992; Borrelli et al., 2021; S. A. Robinson et al., 2020; Vivian & Miczek, 1991). On P14, we observed increased USVs, specifically in FVB/NJ female pups during spontaneous morphine withdrawal.

The severity of NOWS and the need for treatment are determined by observer-rated scales using the Finnegan Neonatal Abstinence Scoring System (FNASS) (Finnegan et al., 1975). Thus, NOWS assessment is subjective and may contribute to unnecessary treatment interventions and lengthy hospital stays (Jansson et al., 2009; Milliren et al., 2018; Timpson et al., 2018). Therefore, unbiased methods for assessing withdrawal symptoms may lead to better prediction of optimal treatment plans, improved infant outcomes, and reduced hospital costs. NOWS infants present with excessive, high-pitched cries that can be differentiated from non-NOWS cries (Blinick et al., 1971). Again, the severity of NOWS cries is subjective and can be interpreted differently between listeners. A recent study revealed that infants with NOWS emit cries with distinct features unrecognizable by humans, such as altered frequency formants, utterances, and amplitude (Manigault et al., 2022). Thus, alterations in acoustic features of cries may be associated with the severity of NOWS, which could be useful for predicting the need for pharmacological treatment compared to subjective evaluation.

A major goal of this study was to identify the USV spectrotemporal syllable profile associated with the negative affective state of neonatal morphine withdrawal. For P14, we identified a robust increase in Complex 3 syllables in mice from two genetic backgrounds, implicating Complex 3 syllables as a potential preclinical biobehavioral marker for symptomatic severity of the negative affective state associated with morphine withdrawal. Furthermore, morphine-withdrawn FVB/NJ females emitted a greater proportion of Complex 3 syllables compared to morphine-withdrawn males, suggesting female pups experience a more severe affective withdrawal state. However, sex differences in Complex 3 emissions during neonatal opioid withdrawal may depend on genetic background as both female and male outbred CFW pups emitted more Complex 3 syllables compared to saline-treated pups. This female-specific effect potentially depends on genetic background, as morphine-withdrawn CFW mice of both sexes showed increased USVs. Alternatively, it should be noted that the lack of sex difference in CFW mice could potentially be explained by the FVB/NJ pups receiving a lower dose of morphine (10mg/kg) rather than genetic background. For example, there could be impaired morphine metabolism and clearance at 10 mg/kg that could underlie the female-specific increase in Complex 3 syllables, whereas the higher dose in CFWs (15 mg/kg) could override this pharmacokinetic effect on behavior.

We analyzed the transcriptome of the brainstem and midbrain tissue due to its role in morphine withdrawal and the production of vocalizations in rodents. On P16 during morphine withdrawal, there was an upregulation of *Oprk* in the brainstem, which codes for kappa opioid receptors (KOR), and increased expression of *Pdyn*, which codes for dynorphin, in the midbrain. KORs and their endogenous ligand, dynorphin (Mansour et al., 1995), mediate withdrawal-associated dysphoria in rodents and humans (Chartoff & Mavrikaki, 2015; Land et al., 2008; Leconte et al., 2022; Shippenberg et al., 2007). Studies have shown that projections from the periaqueductal grey (PAG) in the midbrain to the nucleus retroambiguus (RAm) of the brainstem are necessary for USV production in adult mice (Jürgens, 2002, 2002; Tschida et al., 2019). Therefore, increased dynorphin release in the midbrain may activate KOR in the brainstem and contribute to enhanced withdrawal-induced dysphoria and USVs. We assessed KOR antagonism as a potential target for reducing negative affect in the morphine-withdrawn state. Pre-treatment with the KOR antagonist, nor-BNI on P14 (a known postnatal day for morphine withdrawal-induced increase in USVs), decreased USVs exclusively in morphine-treated females during withdrawal on P15 (20 h pre-treatment) but had no effect on the composition of syllable emissions. This result suggests a female-specific component of KOR in mediating USV emissions and by extension, the severity of the aversive state, but it does not account for the increase in Complex 3 syllables observed during withdrawal. Studies in adult rats have shown that the KOR antagonist nor-BNI reduces withdrawal symptoms (wet dog shakes, jumps, weight loss, and conditioned place aversion) in morphine-dependent rats (Kelsey et al., 2015), attenuates ethanol withdrawal-induced decreases in immobility during the Forced Swim Test (Jarman et al., 2018), and blocks the decrease in open arm explorations in the elevated plus maze during ethanol withdrawal (Valdez & Harshberger, 2012). Most importantly, nor-BNI can block the increase in aversive 22-kHz USVs in rats during ethanol withdrawal and following administration of U50,488h (Berger et al., 2013). Administration of KOR agonist U50,488h was sufficient to increase USVs on P10 in both sexes and on P14, specifically in females. Rat studies have observed enhanced USVs in neonates following U50,488h administration (Carden et al., 1991, 1994; Nazarian et al., 1999). However, few, if any, of these studies have described the spectrotemporal syllable profile. In contrast to our prediction, U50,488h induced a decrease (rather than an increase) in Complex 2, -3, and -4 syllables on P14. It is likely that exogenously administered U50,488h is activating additional circuitry beyond that which is driving increased USVs during morphine withdrawal (e.g., sensory, affective, and/or motor circuitry) that underlies the reduction in Complex 3 emissions. Together, these data are consistent with a female-specific role of KOR activation on P14–P15 in mediating overall USVs but not the type of syllables emitted, indicating that changes in syllable profiles during morphine withdrawal involve additional mechanisms.

Interestingly, there was a significant upregulation of *Slc6a3* (dopamine (DA) transporter, DAT) in the brainstem. Activation of KOR at DAergic terminals inhibits DA release and can promote negative affective states that drive drug-seeking behavior (Shippenberg et al., 1993; Zachry et al., 2021). Accordingly, increased DAT expression would be expected to increase synaptic DA uptake and further support a negative affective state. In neonatal rats, KOR activation with U50,488h increased USV emissions (Carden et al., 1991, 1994; Nazarian et al., 1999), which can be attenuated by DA receptor agonists (Nazarian et al., 1999). Although DAergic neurons in the midbrain do not project to lower parts of the the brainstem (Reyes et al., 2009), activation of KOR on DAergic projections in other regions, such as the ventral tegmental area (VTA), to forebrain structures, including nucleus accumbens (Margolis et al., 2003, 2006; Speciale et al., 1993), may contribute to reduced brainstem DA levels. Thus, increased *Oprk1* and *Slc6a3* expression in the brainstem together could reflect changes in expression in, e.g., DAergic forebrain terminals and together contribute to the aversive state during withdrawal and mediate enhanced USV emission.

Sex differences in reward and stress-related pathways mediated by the dynorphin/KOR system have been observed in rodents and humans (Becker & Chartoff, 2019; Conway et al., 2019; Russell et al., 2014; Vijay et al., 2016). In adult rats, U50,488h-mediated KOR activation decreases reward motivation and induces greater thermal analgesic effects in females compared to males (Bartok & Craft, 1997; Conway et al., 2019; Russell et al., 2014). Human studies have also observed increased analgesic efficacy of KOR agonists in females versus males (Gear et al., 1996; Mogil et al., 2003). Female rats are less sensitive to the DA-reducing and depressive-like effects of KOR agonists (Conway et al., 2019; Laman-Maharg et al., 2018; Russell et al., 2014). These discrepant results in adult rodents versus neonatal females in our study could be due to differences in the immature and rapidly evolving circuitry in neonates versus adults. Rodent studies have shown spatiotemporal variations in KOR expression during development in rats and mice (Georges et al., 1998; Tan et al., 2018; Zhu et al., 1998). Additionally, adult human studies have shown sex differences in KOR availability and expression (Vijay et al., 2016). Time-dependent sex differences in the dynorphin/KOR circuitry might also explain why U50,488h increased USVs in both sexes on P10 but only in females on P14.

There are several limitations to this study. First, the third trimester-approximate model involves neonatal injections, potentially affecting maternal care quality between morphine- and saline-treated pups. There is conflicting evidence regarding the effects of opioid exposure (both to the dam and pups) on maternal care behavior, although evidence indicates that it is dependent on maternal opioid exposure rather than pup exposure (Alipio et al., 2021; Grecco et al., 2021; Smith et al., 2022). In our model, dams were not exposed to morphine – thus, behavioral differences between morphine- and saline-treated pups are most likely due to morphine withdrawal and not maternal care deficits. An additional limitation is that we used the KOR antagonist, nor-BNI, which has a delayed onset of effect (Kishioka et al., 2013; Munro et al., 2012), requiring modifications to our NOWS paradigm. Nor-BNI also has early mu-opioid receptor antagonistic effects, which could induce withdrawal shortly after administration. Nonetheless, future studies could implement faster-acting KOR antagonists, such as CERC-501(LY2456302) (Rorick-Kehn et al., 2014) to assess USVs on P14 rather than P15. However, CERC-501 also binds to mu-opioid receptors, has a lower binding affinity for KOR than nor-BNI and has long-acting effects at KOR (Rorick-Kehn et al., 2014). Additionally, using constitutive or Cre-dependent KOR-knockout mouse models could further elucidate the relationship between the dynorphin/KOR system and USVs during neonatal morphine withdrawal.

We observed robust sex differences in USV profiles in a third trimester-approximate treatment regimen for NOWS model traits. Notably, we identified a unique USV profile, as characterized by an increase in the proportion of Complex 3 syllables, associated with morphine withdrawal. We propose enhanced Complex 3 emissions as a novel marker for withdrawal-induced dysphoria, which can serve as a new phenotype for testing the efficacy of novel therapeutics in mice. We identified a KOR component of mediating USV emissions in morphine-withdrawn FVB/NJ, suggesting that the KOR system in females may contribute to greater withdrawal distress and dysphoria compared to males. These results contribute to our understanding of the biological mechanisms underlying sex differences in NOWS symptom severity.

More broadly, we provide a quantitative, machine learning-based approach to studying negative internal states associated with neonatal opioid withdrawal in mice, identifying specific USV profiles as indicators of opioid withdrawal. Examining adaptations in spectrotemporal features of neonatal mouse vocalizations unique to neonatal opioid withdrawal provides a translational approach for modeling NOWS severity beyond excessive crying in the clinic and assessing potential experimental therapeutics in reducing emotional symptom severity.

## ACKNOWLEDGMENTS

We thank the Boston University Genome Science Institute and Microarray and Sequencing Resource Core for the Illumina mRNA-seq library prep kit and flow cell awarded to Jacob A. Beierle.

## DECLARATIONS

### Funding

This study was funded by the National Institute on Drug Abuse (NIDA) grants U01DA050243 (C.D.B.), U01DA55299 (C.D.B.), T32DA055553 (B.M.B.)

### Conflicts of interest

The authors have nothing to disclose.

### Author Contributions

K.K.W.: Data collection, analysis, and writing of manuscript; T.M.: Data collection; K.J.: Data collection; C.S.M.: Data collection; N.M.A.; Data collection; K.T.R.: Data collection; M.B.R.; Data collection; J.A.B.: Data collection and RNAseq analysis; S.A.M.: Data collection; E.J.S.: Data collection; B.M.B.: Data collection; W.B.L: Gene pathway enrichment and analysis; K.N.B.; CFW Data collection; E.J.Y.; CFW Data collection; E.M.W.: Writing of manuscript; C.D.B.: Experimental design, analysis input, writing of manuscript.

## Notes

**CONFLICTS OF INTERESTS** The authors have nothing to disclose.

### Competing Interest Statement

The authors have declared no competing interest.

### Summary of Updates

In the revised manuscript, we have added new qPCR (Fig.4B - G) and enrichment analysis (Fig.5) data. The discussion is updated to include these new results. We revised Fig.7, which contained a figure ordering mistake (switched 7A & 7D). We provide an additional supplemental figure (Fig. S5) illustrating our neonatal brain dissection method. Data analysis and figure generation for Fig. 5 were performed by William B. Lynch, who we added to our new author list. We updated Fig.S6 for visual clarity. We updated all figures to display sex (open circles = females, closed circles = males). All time-series data contains individual adjusted p values. We corrected a mistake in the results section for Fig 2 - there were no time-dependent effects. We corrected a mistake in the nor-BNI methods, where pups received 10mg/kg morphine (NOT 15mg/kg). We provide additional justification on nor-BNI dose selection and maternal care behavior.

