## Supplementary material for "The ultrasonic vocalization (USV) syllable profile during neonatal opioid withdrawal and a kappa opioid receptor component to increased USV emissions in female mice": Supplementary_Material.pdf

**SUPPLEMENTARY INFORMATION**

### Supplementary Methods

#### Morphine administration in FVB/NJ pups from P1 to P15

Preclinical and clinical studies indicate that the third trimester is both necessary and sufficient to induce a neonatal withdrawal state(Craig et al., 2003; Rice & Barone, 2000; Semple et al., 2013). This model, comprising P1-P14/P15 morphine exposure, allows us to precisely control individual dosing and avoid maternal exposure and downstream consequences on offspring behavior – two critical requirements for future quantitative genetic studies that we plan to conduct. Pups were sexed on P1, and each litter was approximately treatment- and sex-balanced to control for cage environment across treatments. Behavioral phenotyping occurred on P7 and P14, 16 h following morphine administration during spontaneous withdrawal.

#### P8 and P15 phenotyping of mice used for bulk RNA-seq

The subset of FVB/NJ mice used for RNA-seq were also phenotyped on P8 and P15 following their morning morphine injections (during a state of withdrawal alleviation as opposed to under a state of withdrawal). In this case, morning (0900h) and evening (1700h) injections on P7 and P14 resumed after phenotyping. On P8 and P15 we wished to assess USVs while the neonates were under the influence of morphine (versus saline). Neonates received their maintenance dose of morphine (10mg/kg, s.c., Sigma-Aldrich) or saline (0.9%, 20 ul/g, s.c.) at 0830h and placed in their home cage. After 30 min, USVs were recorded for 10 min (P8) or 15 min (P15). On P16, 16h post-injections (i.e., during the withdrawal state), mice were euthanized by live, rapid decapitation, and brains were removed from the skull and dura and immediately collected for bulk mRNA sequencing (RNA-seq). All mouse handling and behavioral testing were performed by female experimenters.

#### Morphine administration in CFW pups from P1 to P14

From P1-P14, injections of either morphine sulfate pentahydrate (15 mg/kg, s.c.; Sigma-Aldrich) or saline (0.9%, 20 ul/g, s.c.) were administered twice daily at 0900h and 1700h. Each litter was approximately treatment- and sex-balanced to control for cage environment across treatments. Behavioral phenotyping occurred on P7 and P14, 16h following morphine administration during spontaneous withdrawal.

#### Supervised USV classification

DeepSqueak (Coffey et al., 2019) and MATLAB (version 2022a) were used to detect individual USVs from mouse pup audio (.wav files) obtained from Avisoft. Approximately 16,600 USVs belonging to each sex and treatment were manually classified in DeepSqueak based on previously characterized syllables and spectrotemporal features (Caruso et al., 2022; Portfors, 2007)(**Table.S1**). Excel files (.xlsx) containing labeled USVs and accompanying spectrotemporal features (from both morphine- and saline-treated females and males) were exported. Call length (s), principle frequency (kHz), low frequency (kHz), peak frequency (kHz), delta (change) frequency (kHz), and mean power (dB/Hz) were used to train a random forest (sk.learn) classifier in Python. Minority syllable classes were oversampled so that each syllable was weighed equally during training. The labeled dataset was split into training (70%) and testing (30%) sets. Overall, the model had an AUC of 99.6% (accuracy = 94.2%; precision = 94.1%; recall = 94.3%). Thus, we were confident in its performance on unlabeled datasets. Python script for training a random forest model and USV classification are available on GitHub (https://github.com/camronbryant/NOWS_USV_classifier).

**Bulk RNA-seq of brainstem from morphine-withdrawn pups**

Brainstem RNA was extracted from eight FVB/NJ pups (2 mice per sex per treatment) using Trizol (Qiagen), ethanol precipitation, filtering columns (Qiagen), DNAse digestion (Qiagen), and elution with RNAse and nucleotide-free water. RNA library preparation (poly-A selection) and RNA-seq (100x100 bp paired-end reads) were conducted at the Genome Sciences Institute at Boston University Chobanian and Avedisian School of Medicine on an Illumina NextSeq2000 using a 200-cycle P2 flowcell. We used the R/Bioconductor package "scruff"(Wang et al., 2019) to conduct demultiplexing, read alignment, read counting, quality checking and data visualization. Reads were trimmed for quality using Trimmomatic and were then aligned to the mm10 mouse reference genome to generate BAM files, which were in turn aligned to the reference genome using STAR. For differential gene expression analysis, featureCounts was used to count reads that mapped to the "exon" feature in a GTF file obtained from Ensembl (GRCm38). Genes without a minimum of 10 reads per million in at least five of eight samples were excluded from analysis using EdgeR (Robinson et al., 2010) and differential gene expression analysis of normalized counts was conducted considering the effect of Treatment while controlling for Sex and Litter. Results are reported using the topTable (limma) function.

**Primer sets for RT-qPCR**

| Gene | Forward (5' – 3') | Reverse (5' – 3') |
|---|---|---|
| ***Oprk1*** | AGTGGGCAATTCTCTGGTCA | GCACATCTCCAAAAGGCCAA |
| ***Pdyn*** | AAAACCCAGCTCCTAGACCC | ACTCCGATGCAGTTCCTCAT |
| ***Slc6a3*** | GCATCCTGTTCACATATTACAC | TTGTCTCCCAACCTGAATTC |

**Effect of the kappa opioid receptor agonist U50,488h on ultrasonic vocalizations**

We piloted the effects of five doses of U50,488h (0.625, 1.25, 2.5, 5.0 mg/kg) on vocalizations and locomotor activity. On P10 and P14, pups were injected with either U50,488h (0.625mg/kg, s.c.) or saline and returned to their home cage 10 min prior to testing. P10 was chosen as the first assessment day, given that vocalizations significantly decline after this developmental time point (Elwood & Keeling, 1982) and to ensure that a sufficient level of USVs would be emitted to detect either decreases or increases in USVs in response to U50,488h. USVs were recorded at 0900 h for 15 min. Pups continued to receive saline injections following P10 testing until testing for the behavioral effects of U50,488h again on P14, which is when we normally record USVs.

**Supplementary Tables**

**Table S1. USV syllable characteristics.** USV syllables and their spectrotemporal features (Caruso et al., 2022; Portfors, 2007). Representative images were collected from DeepSqueak (Coffey et al., 2019).

| Syllable | Definition | Frequency Change | Spectrotemporal Shape (Frequency/Time) |
|---|---|---|---|
| Chevron | Inverted U-shaped | Increase then decrease > 5 kHz | 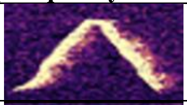 |
| Complex | > 3 directional changes in frequency | Changes > 5 kHz | 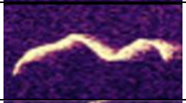 |
| Complex 2 | 2 components with no time separation | 1 directional change in frequency > 5 kHz | 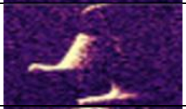 |
| Complex 3 | 3 components with no time separation | 2 directional changes in frequency > 5 kHz | 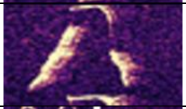 |
| Complex 4 | 4 components with no time separation | 3 directional changes in frequency > 5 kHz | 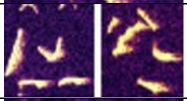 |
| Downward | Duration > 10ms | Decrease > 5 kHz | 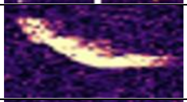 |
| Flat | Duration > 10ms | Changes < 5 kHz | 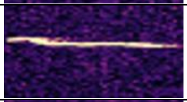 |
| Reverse Chevron | U-shaped | Decrease then increase > 5 kHz | 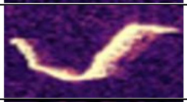 |
| Short | Duration < 10ms | Any | 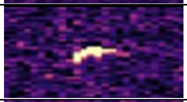 |
| Upward | Duration > 10ms | Increase > 5 kHz | 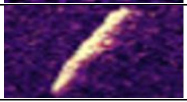 |

**Table S2.** Upregulated genes ($log_2FC \geq 0.26$; $p < 0.01$) between morphine (2F, 2M) vs. saline (2F, 2M) mice

*Provided in a separate document

**Table S3.** Downregulated genes ($log_2FC \leq -0.26$; $p < 0.01$) between morphine (2F, 2M) vs. saline (2F, 2M) mice

*Provided in a separate document.

**Supplementary Figures and Legends**

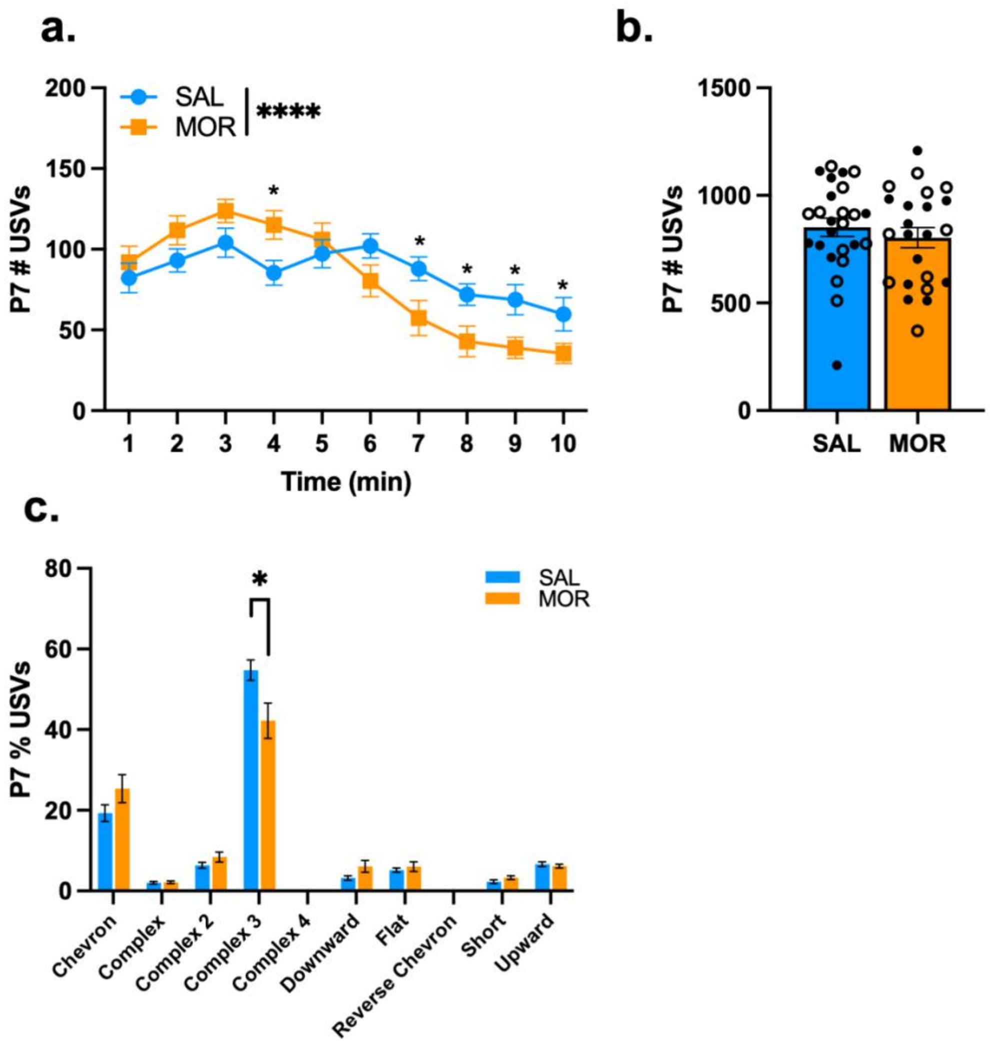


**Fig. S1. USV profiles of FVB/NJ pups during spontaneous morphine withdrawal on P7.** Data are plotted as the mean ± SEM. Saline = blue lines/bars; Morphine = orange lines/bars. Closed circles = Females; Open circles = males. **(a) P7 USV Emission**: The effect of Morphine Treatment was dependent on Time (β = -6.67, SE = 1.282, t(432) = -5.20 ****$p < 0.0001$), where morphine-treated pups vocalized more than saline-treated pups during the 4 min interval (β = 29.69, SE = 12.3, t(335) =2.47, *$p = 0.016$), and less than saline-treated pups from 7 – 10 min (all *$p \leq 0.049$). **(b) P7 Total USV Emission**: There was no effect of Sex (β = 10.46, SE = 89.36, t(44) = 0.12, $p = 0.91$) or a Morphine Treatment x Sex interaction (β = -16.37, SE = 129.61, t(44) = -0.13, $p = 0.90$). There was no effect of Morphine Treatment (β = -48.70, SE = 63.09, t(46) = -0.77, $p = 0.44$). **(c) P7 Syllable Profile**: There was no effect of Sex ($p \geq$ 0.13) or a Morphine Treatment x Sex interaction ($p \geq$ 0.084)for any syllable type. Morphine Treatment was associated with a decrease in Complex 3 emissions (β = -0.13, SE = 0.049, t(46) = -2.53, *$p = 0.015$). SAL, n = 25 (12F, 13M); MOR, n = 23 (13F, 10M)

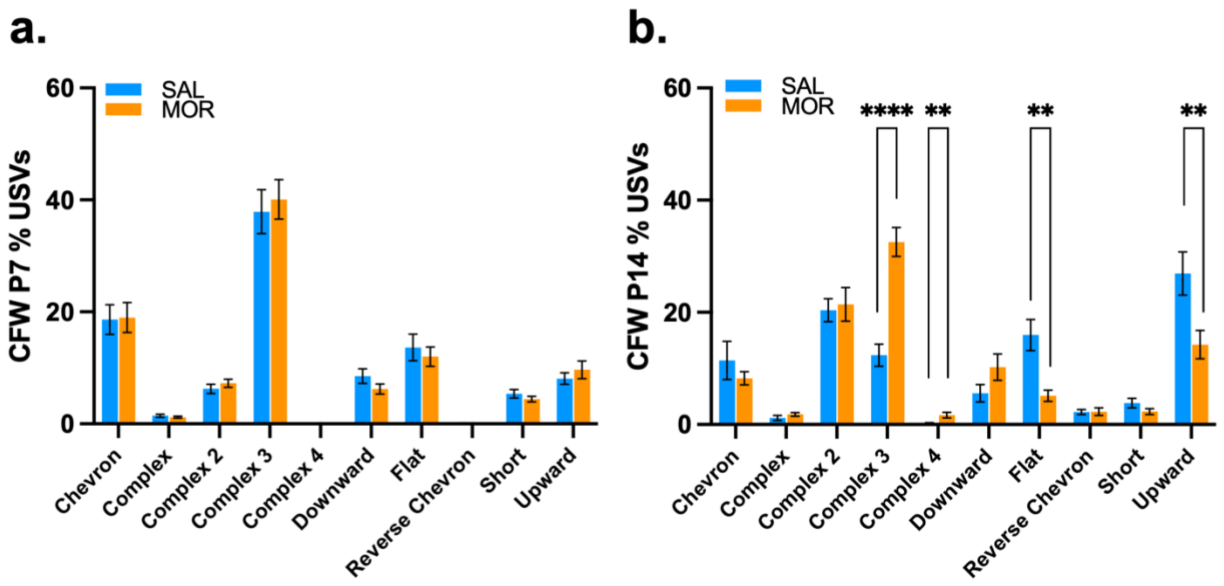


**Fig. S2. USV profiles of CFW pups during spontaneous morphine withdrawal on P7 and P14.** Data are plotted as the mean ± SEM. Saline = blue bars; Morphine = orange bars. **(a) P7 CFW Syllable Profile:** There was no effect of Sex (all $p \geq 0.18$), so Sex was removed from the model. Morphine Treatment had no effect on the proportion of syllables emitted (all $p \geq 0.14$). **(b) P14 CFW Syllable Profile**: There was no effect of Sex (all $p \geq 0.062$) for any syllable type, so Sex was removed from the model. Morphine Treatment was associated with an increase in Complex 3 (β = 0.20, SE = 0.032, t(56) = 6.29, ****$p < 0.0001$) and Complex 4 (β = 0.015, SE = 0.0051, t(56) = 2.86, **$p$ = 0.0059), and a decrease in Flat (β = -0.11, SE = 0.030, t(56) = -3.67, ***$p$ = 0.00055) and Upward (β = -0.13, SE = 0.047, t(56) = -2.73, **$p$ = 0.0086) syllables

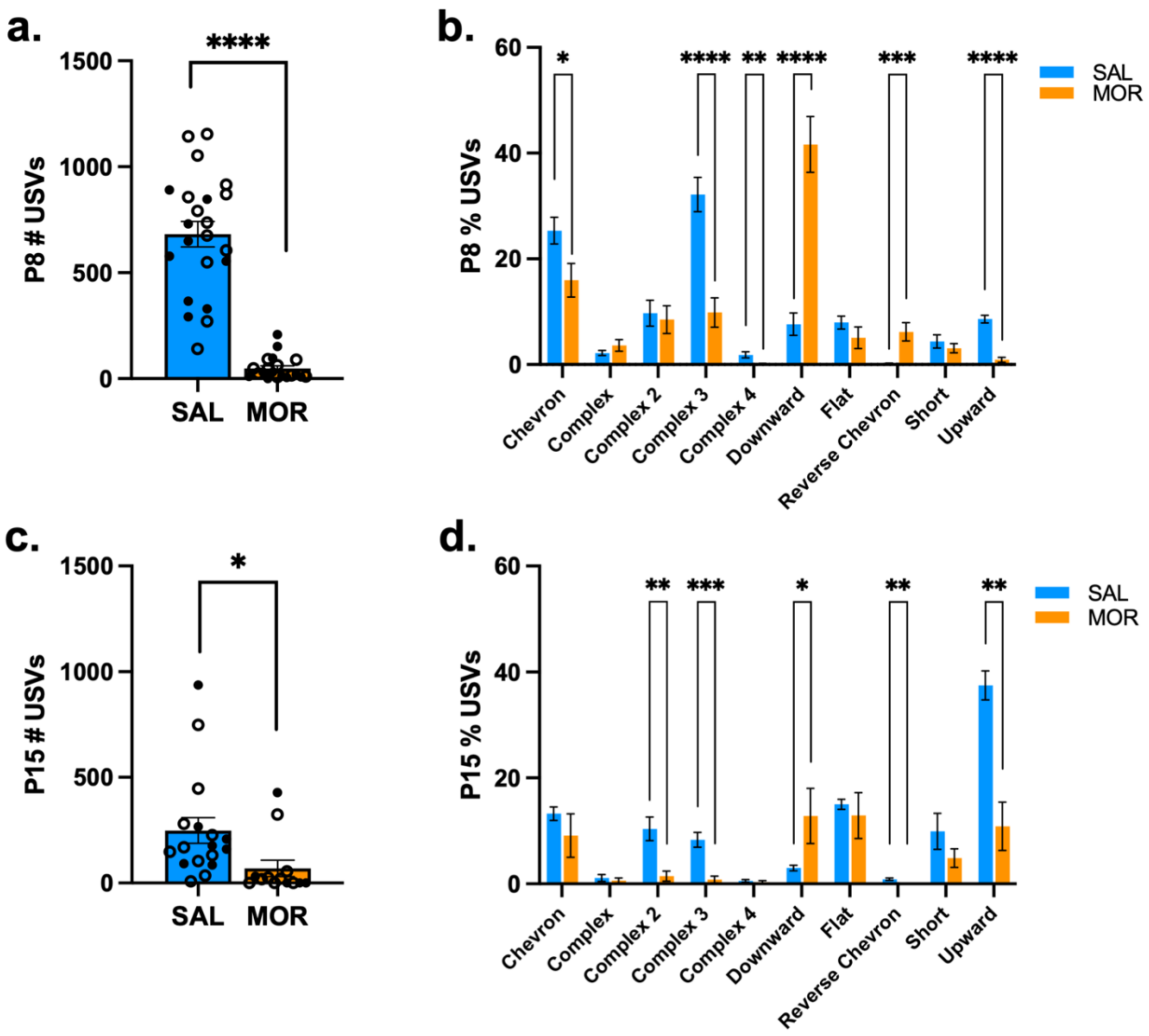


**Fig. S3. Effect of morphine administration on USV characteristics on P8 and P15.** Mice continued to receive their respective treatment at 0900 h (saline or 10 mg/kg morphine, i.p.) and 30 min later, mice were individually placed into the sound attenuating chambers for USV recordings. Data are plotted as the mean ± SEM. Saline = blue bars; Morphine = orange bars. Closed circles = Females; Open circles = males. Saline Treatment and Female Sex were set as the reference variables for linear regression. **(a) P8 Total USV Emission**: There was no effect of Sex (β = 169.49, SE = 88.37, t(38) = 1.92, *p* = 0.063), so Sex was removed from the model. Morphine Treatment was associated with a decrease in USVs (β = -633.32, SE = 64.27, t(40) = -9.85, ****$p < 0.0001$). **(b) P8 Syllable Profile**: There was no effect of Sex (all $p \geq 0.056$) on for any syllable type, so Sex was removed from the model. Morphine Treatment was associated with a decrease in Chevron (β = -0.094, SE = 0.040, t(40) = -2.33, *$p = 0.025$), Complex 3 (β = -0.22, SE = 0.043, t(40) = -5.18, ****$p < 0.0001$), Complex 4 (β = -0.017, SE = 0.0060, t(40) = -2.88, **$p = 0.0064$), and Upward (β = -0.077, SE = 0.0088, t(40) = -8.74, ****$p < 0.0001$) syllables, and an increase in Downward (β = 0.34, SE = 0.055, t(40) = 6.17, ****$p < 0.0001$) and Reverse Chevron (β = 0.060, SE = 0.016, t(40) = 3.67, ***$p = 0.00071$) syllables. **(c) P15 Total USV Emissions**: There was no effect of Sex (β = -90.18, SE = 108.08, t(26) = -0.83, $p = 0.41$), so Sex was removed from the model. Morphine Treatment was associated with a decrease in USVs (β = -179.19, SE = 76.74, t(28) = -2.34, *$p$ = 0.027). **(d) P15 Syllable Profile**: There was no effect of Sex (all $p \geq 0.23$) for any syllable type, so Sex was removed from the model. Morphine Treatment was associated with a decrease in Complex 2 (β = -0.089, SE = 0.027, t(28) = -3.31, **$p$ = 0.0026), Complex 3 (β = -0.075, SE = 0.017, t(28) = -4.40, ***$p$ = 0.00014), Reverse Chevron (β = -0.0089, SE = 0.0026, t(28) = -3.36, **$p$ = 0.0023), and Upward (β = -0.27, SE = 0.051, t(28) = -5.25, ****$p$ < 0.0001) syllables, and an increase in Downward (β = 0.098, SE = 0.046, t(28) = 2.15, *$p$ = 0.041) syllables. SAL, n = 17–22 (6–9F, 11–13M); MOR, n = 13–20 (5–11F, 8–9M)

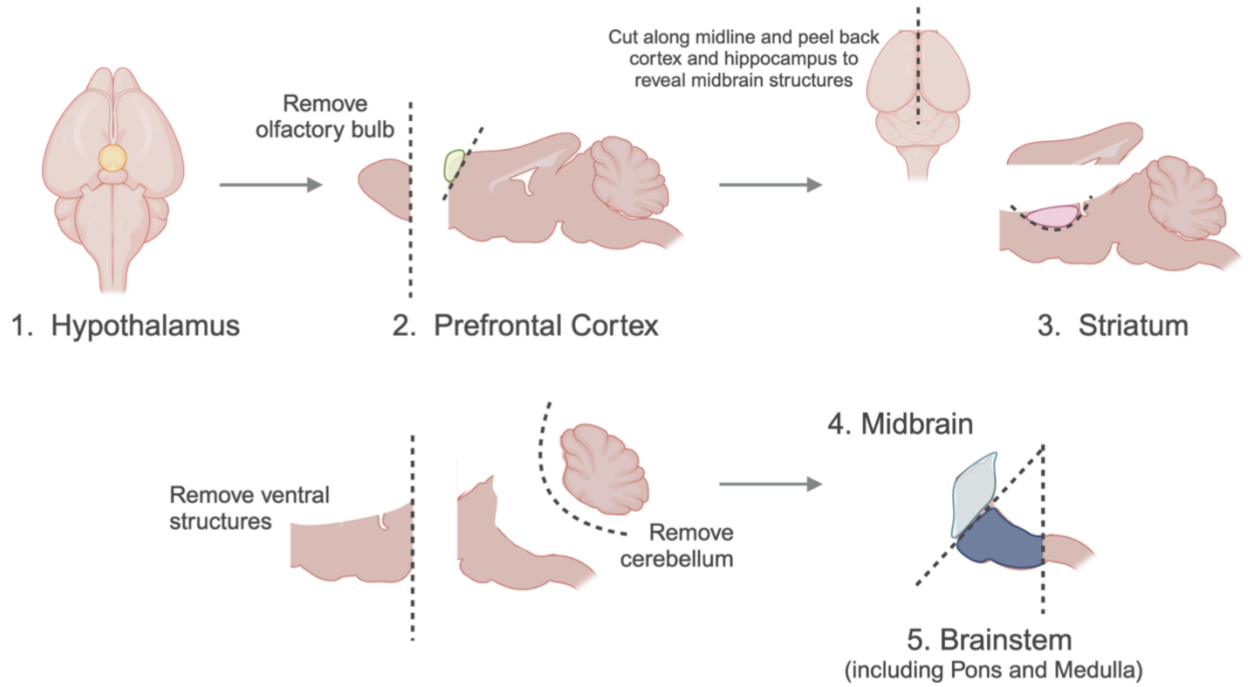


**Fig. S4. Freeform neonatal brainstem dissection schematic.** *Created with Biorender*

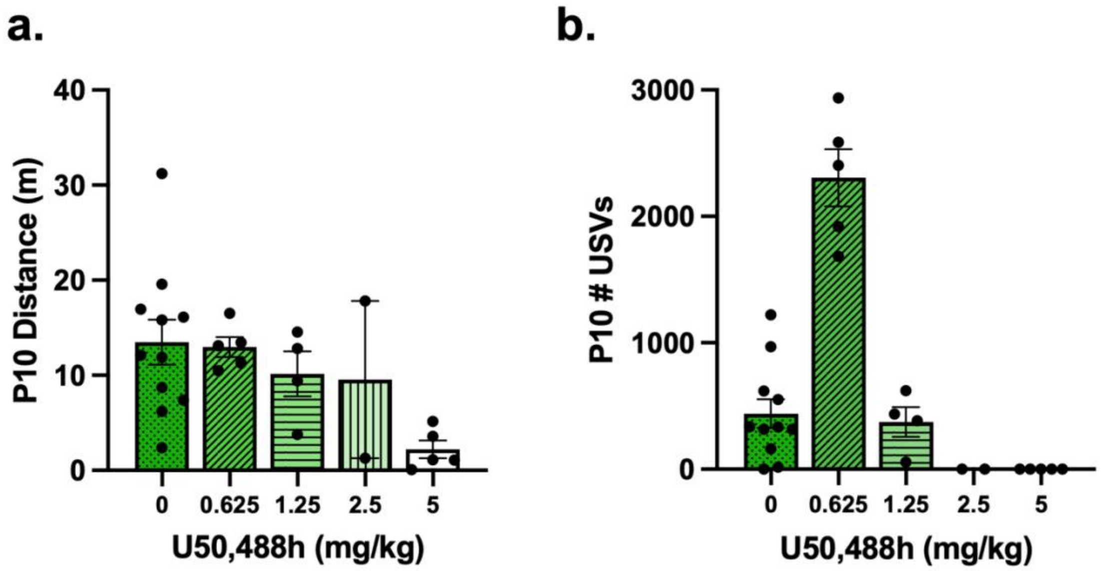


**Fig. S5. Pilot experiment examining the effect of U50,488 doses on dose-response activity on P10 in morphine-naïve FVB/NJ pups.** Data are plotted as the mean ± SEM. The U50,488h dose increases from left to right bars on the graphs. 0mg/kg (saline) = darkest green/dot pattern bars; 0.625mg/kg = medium-dark green/diagonal line pattern bars; 1.25mg/kg = medium green/horizontal line pattern bars; 2.5mg/kg = light green/vertical line pattern bars; 5mg/kg = white/no pattern bars. Mice were treated from P1 up to P10 with twice daily saline injections (0.9%, 20 ul/g, s.c.) **(a) P10 Total Distance**. High doses of U50,488h suppressed locomotor activity. **(b) P10 Total USV Emission.** High doses of U50,488h reduced vocalizations. SAL, n = 11 (5F, 6M); 0.625, n = 8 (8F, 0M); 1.25, n = 4 (2F, 2M); 2.5, n = 2 (0F, 2M); 5.0, n = 5 (3F, 2M). Because this was a pilot study with limited sample sizes and unbalanced sexes, we did not perform statistical analyses on this dataset
