## Supplementary material for "The ultrasonic vocalization (USV) syllable profile during neonatal opioid withdrawal and a kappa opioid receptor component to increased USV emissions in female mice": Table.S2.pdf

**Table S2.** Upregulated genes ($log_2FC \geq 0.26$; $p < 0.01$) between morphine (2F, 2M) vs. saline (2F, 2M) mice.

| Gene | logFC | Fold Change | t value | p value | Adjusted p value | B |
|---|---|---|---|---|---|---|
| **Six3os1** | 5.33598317 | 40.3915938 | 3.64298355 | 0.00523234 | 0.06081197 | -3.5383729 |
| **Slc6a3** | 3.69248289 | 12.9284991 | 3.48624404 | 0.00670303 | 0.06866063 | -2.1622714 |
| **R3hdml** | 3.05887635 | 8.33323318 | 4.47698817 | 0.0014783 | 0.03163717 | -3.9759112 |
| **Oas3** | 2.85657023 | 7.24291393 | 3.29926258 | 0.00904003 | 0.07994289 | -4.1106985 |
| **Lhx9** | 2.44091937 | 5.42987643 | 3.52193843 | 0.00633378 | 0.0665086 | -2.0343856 |
| **AC091309.7** | 2.43745597 | 5.41685686 | 3.38851805 | 0.00783359 | 0.07476748 | -4.0766463 |
| **Apol7a** | 1.68046417 | 3.20531062 | 3.54044129 | 0.0061508 | 0.06566202 | -3.825209 |
| **Krt90** | 1.46209744 | 2.75508617 | 3.31036135 | 0.00888004 | 0.07947189 | -3.7579646 |
| **Uox** | 1.37394374 | 2.59178085 | 3.58623284 | 0.00572136 | 0.06342905 | -3.6146818 |
| **Cwh43** | 1.36718732 | 2.57967142 | 3.39150623 | 0.00779622 | 0.07467945 | -3.5849209 |
| **Myh8** | 1.35761388 | 2.5626099 | 3.35230371 | 0.00830153 | 0.07679488 | -3.0954347 |
| **Kcnj4** | 1.31442401 | 2.48703017 | 3.64499367 | 0.00521584 | 0.06076284 | -2.2415039 |
| **C1ql4** | 1.16852936 | 2.24782443 | 3.25594276 | 0.00969368 | 0.08264749 | -2.3974283 |
| **Dnajc22** | 1.1650731 | 2.24244577 | 3.36563813 | 0.00812594 | 0.0759622 | -3.3193326 |
| **Gda** | 1.1357803 | 2.19737379 | 3.38119665 | 0.00792593 | 0.07520578 | -2.3005164 |
| **Pth2r** | 1.01654322 | 2.02306577 | 4.51217269 | 0.00140455 | 0.0309625 | -2.0015258 |
| **Chrm1** | 1.00884038 | 2.01229299 | 4.02618797 | 0.00289292 | 0.04427057 | -1.5863565 |
| **Vip** | 0.99823404 | 1.99755336 | 3.56572368 | 0.00590968 | 0.06417315 | -2.0039175 |
| **Slc10a1** | 0.97977073 | 1.97215197 | 3.42588113 | 0.00737951 | 0.07239561 | -3.2288999 |
| **Ift88os** | 0.9646741 | 1.9516226 | 3.37586344 | 0.0079939 | 0.075291 | -3.2922018 |
| **Lrrc6** | 0.93000908 | 1.90528799 | 4.54761101 | 0.00133422 | 0.03032392 | -1.592237 |
| **Oprm1** | 0.91211409 | 1.88180102 | 3.36457509 | 0.00813979 | 0.07600746 | -2.3664928 |
| **Cdhr1** | 0.87935443 | 1.83955197 | 4.26910737 | 0.00200754 | 0.03671917 | -1.0727121 |
| **Deup1** | 0.83931043 | 1.78919474 | 3.58301629 | 0.00575048 | 0.06342905 | -2.6543241 |
| **Nphs2** | 0.80389443 | 1.74580742 | 3.51629577 | 0.0063907 | 0.06689809 | -2.8688654 |
| **Cdh9** | 0.78785298 | 1.72650317 | 4.07058034 | 0.00270441 | 0.0426634 | -1.2827646 |
| **Gucy2f** | 0.77030239 | 1.70562725 | 4.51280546 | 0.00140326 | 0.0309625 | -1.5642778 |
| **Zpbp** | 0.76660858 | 1.70126582 | 5.89946812 | 0.00021512 | 0.01474675 | 0.23103637 |
| **Gabrb2** | 0.75824286 | 1.69142928 | 5.27355417 | 0.00048488 | 0.02001271 | 0.16229905 |
| **Irs4** | 0.75577998 | 1.68854424 | 3.56817778 | 0.00588681 | 0.06416633 | -2.0671469 |
| **Kcna3** | 0.75314853 | 1.68546718 | 6.06902549 | 0.00017419 | 0.01447525 | 0.99718396 |
| **Arntl2** | 0.74808002 | 1.67955614 | 4.16521082 | 0.0023447 | 0.04010253 | -1.4354375 |
| **Dnah12** | 0.7448234 | 1.67576912 | 3.31888825 | 0.00875913 | 0.07899051 | -2.5012089 |
| **Cntnap5b** | 0.73669799 | 1.66635755 | 5.12748319 | 0.0005908 | 0.02131828 | 0.11039875 |
| **Cntnap5c** | 0.7360122 | 1.66556563 | 7.34970817 | 3.97E-05 | 0.01161523 | 1.92613644 |
| **Cntnap3** | 0.72806228 | 1.65641282 | 3.88853721 | 0.00357123 | 0.05014584 | -1.7891183 |

| | | | | | | |
|---|---|---|---|---|---|---|
| **AC133601.1** | 0.72457991 | 1.65241941 | 3.40695546 | 0.00760598 | 0.0735327 | -2.5807908 |
| **Crb1** | 0.71738254 | 1.64419629 | 3.84343973 | 0.0038285 | 0.05177529 | -1.7608658 |
| **Mcm6** | 0.71073643 | 1.63663934 | 4.40123306 | 0.00165151 | 0.03374151 | -0.9818587 |
| **Serpina3h** | 0.69520193 | 1.61911104 | 3.45670007 | 0.00702565 | 0.07012685 | -2.6209949 |
| **Unc5d** | 0.67801301 | 1.59993469 | 6.35887552 | 0.0001225 | 0.01356866 | 1.588477 |
| **AW551984** | 0.6709767 | 1.59215049 | 3.80236035 | 0.00407989 | 0.05366872 | -2.0023846 |
| **Cdca2** | 0.67035215 | 1.59146138 | 3.24562639 | 0.00985644 | 0.08348522 | -2.8721487 |
| **Pappa** | 0.65978656 | 1.57984887 | 3.70611948 | 0.00473945 | 0.05787546 | -1.7807703 |
| **Selenov** | 0.65903777 | 1.57902911 | 3.5063972 | 0.00649186 | 0.06749615 | -2.3602059 |
| **Il1rapl2** | 0.64999427 | 1.56916197 | 4.44091673 | 0.00155823 | 0.03276551 | -1.3339138 |
| **Glra2** | 0.63813631 | 1.55631739 | 4.70631145 | 0.00106242 | 0.02803871 | -0.472322 |
| **AC127249.1** | 0.63461665 | 1.55252516 | 4.62777001 | 0.00118867 | 0.02884275 | -0.5528665 |
| **Zkscan16** | 0.62542325 | 1.54266333 | 6.72161275 | 8.00E-05 | 0.01316315 | 2.01203229 |
| **Cntn5** | 0.62426849 | 1.54142905 | 3.59681913 | 0.00562663 | 0.06288104 | -1.936507 |
| **Tenm1** | 0.6194825 | 1.536324 | 5.75131792 | 0.00025948 | 0.01626727 | 0.86777157 |
| **Evx2** | 0.60711355 | 1.52320862 | 3.30041998 | 0.00902321 | 0.07991985 | -2.3425495 |
| **Chrna7** | 0.60425042 | 1.5201887 | 5.45133838 | 0.00038281 | 0.01842957 | 0.51495138 |
| **Htr4** | 0.59915029 | 1.51482411 | 4.92957694 | 0.0007759 | 0.02445764 | -0.6975707 |
| **Adamts18** | 0.59787005 | 1.51348046 | 4.05288373 | 0.00277795 | 0.04328868 | -1.541955 |
| **Uhrf1** | 0.59501201 | 1.51048516 | 4.23970856 | 0.00209737 | 0.0377079 | -1.2470355 |
| **Wdr31** | 0.59289781 | 1.50827324 | 3.28585065 | 0.00923737 | 0.08092412 | -2.403001 |
| **Plcxd3** | 0.58639457 | 1.5014897 | 6.83939616 | 6.99E-05 | 0.01235543 | 1.85305157 |
| **Kcnv1** | 0.58556961 | 1.50063136 | 3.78710459 | 0.00417763 | 0.05425491 | -1.9869407 |
| **Dgkk** | 0.5816655 | 1.49657596 | 9.19330702 | 6.41E-06 | 0.00870145 | 4.4498269 |
| **Ttn** | 0.5757086 | 1.49040932 | 3.3281243 | 0.00863009 | 0.0782882 | -2.3226408 |
| **Zfhx4** | 0.57356889 | 1.48820048 | 5.70089607 | 0.00027677 | 0.01649839 | 0.77032713 |
| **Asb4** | 0.56569578 | 1.48010116 | 4.66088831 | 0.00113358 | 0.02836303 | -0.5497968 |
| **Zim1** | 0.55419472 | 1.46834881 | 3.62230818 | 0.00540522 | 0.06151649 | -2.0841313 |
| **Ndst4** | 0.55343241 | 1.46757314 | 4.65394548 | 0.0011449 | 0.02848064 | -0.6488386 |
| **Dusp9** | 0.55313648 | 1.46727214 | 3.30884473 | 0.00890173 | 0.07949593 | -2.5187128 |
| **Cdh4** | 0.55229144 | 1.46641296 | 5.20739768 | 0.00053007 | 0.02043832 | 0.18359766 |
| **Kcns2** | 0.55111797 | 1.46522069 | 5.60369206 | 0.00031372 | 0.01704154 | 0.6872884 |
| **Ano5** | 0.55031816 | 1.46440861 | 6.41163531 | 0.00011503 | 0.01356866 | 1.19548283 |
| **Lamc2** | 0.53915624 | 1.45312241 | 3.53542387 | 0.00619987 | 0.06586461 | -2.1412725 |
| **Fabp7** | 0.5371987 | 1.45115206 | 8.38702287 | 1.37E-05 | 0.00983403 | 3.72132161 |
| **Fndc9** | 0.53596706 | 1.44991373 | 3.45633458 | 0.00702974 | 0.07012685 | -2.1629367 |
| **Camk4** | 0.52626186 | 1.4401927 | 5.260769 | 0.00049328 | 0.02001271 | 0.28256139 |
| **Lncenc1** | 0.52037289 | 1.43432592 | 4.03183184 | 0.0028682 | 0.04393214 | -1.6140014 |
| **Tet1** | 0.52017725 | 1.43413143 | 8.80465361 | 9.18E-06 | 0.00983403 | 4.09483008 |

| | | | | | | |
|---|---|---|---|---|---|---|
| **Xkr4** | 0.51762716 | 1.43159872 | 3.5340001 | 0.00621386 | 0.06586461 | -2.1401781 |
| **Slc9a7** | 0.51740165 | 1.43137496 | 5.1753148 | 0.0005536 | 0.02060914 | 0.16101027 |
| **Dchs2** | 0.51459019 | 1.42858828 | 3.83530326 | 0.00387696 | 0.05196265 | -1.6732417 |
| **Slc16a14** | 0.51184705 | 1.42587454 | 5.65348609 | 0.00029416 | 0.01681625 | 0.57361153 |
| **Stxbp5l** | 0.51061028 | 1.42465272 | 4.60788974 | 0.00122311 | 0.02903664 | -0.6163308 |
| **Hrk** | 0.50889352 | 1.42295843 | 5.51255223 | 0.00035325 | 0.0181571 | 0.59726153 |
| **Npy** | 0.50497041 | 1.41909425 | 3.73674132 | 0.0045182 | 0.05652439 | -1.7682384 |
| **Cubn** | 0.50136864 | 1.41555582 | 3.6524663 | 0.005155 | 0.06045473 | -1.978283 |
| **Dcc** | 0.49931789 | 1.41354507 | 3.64917743 | 0.00518169 | 0.06059913 | -2.107832 |
| **Zbtb8b** | 0.4990298 | 1.41326284 | 3.72903043 | 0.00457287 | 0.05695481 | -1.7942746 |
| **Zfp853** | 0.49596193 | 1.41026075 | 3.69523331 | 0.00482082 | 0.05834729 | -1.7963323 |
| **Dll3** | 0.4948898 | 1.4092131 | 4.61275838 | 0.00121458 | 0.02901223 | -0.7736493 |
| **Lrrtm4** | 0.49404339 | 1.40838659 | 4.51327731 | 0.0014023 | 0.0309625 | -0.6870405 |
| **Amn** | 0.48845166 | 1.40293839 | 4.54051338 | 0.001348 | 0.03043199 | -0.6952503 |
| **Apold1** | 0.48695101 | 1.40147986 | 4.05524736 | 0.002768 | 0.04328868 | -1.3016492 |
| **Ccnf** | 0.48197487 | 1.3966542 | 4.04248819 | 0.00282213 | 0.04359044 | -1.5313 |
| **Col25a1** | 0.48042182 | 1.39515152 | 9.2668404 | 6.00E-06 | 0.00870145 | 4.50229441 |
| **Pcdh11x** | 0.47872133 | 1.39350805 | 9.75987561 | 3.89E-06 | 0.00870145 | 4.85795848 |
| **Nwd2** | 0.4765863 | 1.39144733 | 4.24575741 | 0.00207854 | 0.03744927 | -1.1032873 |
| **Kcnh5** | 0.47390703 | 1.38886563 | 3.82229659 | 0.00395578 | 0.05261065 | -1.7011396 |
| **AI504432** | 0.47182953 | 1.38686709 | 4.61545012 | 0.00120989 | 0.02896432 | -0.698757 |
| **Zdbf2** | 0.46684934 | 1.38208787 | 5.27023848 | 0.00048705 | 0.02001271 | 0.16831169 |
| **Gabrg3** | 0.46471998 | 1.38004947 | 5.46166885 | 0.00037764 | 0.01839444 | 0.52422811 |
| **Nell1** | 0.45845669 | 1.37407113 | 6.74011354 | 7.83E-05 | 0.01308018 | 2.02571182 |
| **Papln** | 0.45435829 | 1.37017322 | 4.5406041 | 0.00134782 | 0.03043199 | -0.8676921 |
| **Rgs9** | 0.44983926 | 1.36588807 | 5.23567335 | 0.00051023 | 0.02024575 | 0.24783626 |
| **Lrrc26** | 0.44976988 | 1.36582238 | 3.47954142 | 0.00677484 | 0.06886728 | -2.1797597 |
| **Tenm2** | 0.44954318 | 1.36560778 | 3.2548194 | 0.00971126 | 0.0827125 | -2.8803015 |
| **Lypd6** | 0.44791427 | 1.36406677 | 3.64354684 | 0.00522771 | 0.06081197 | -2.0376615 |
| **Mal2** | 0.4471829 | 1.36337544 | 4.62340552 | 0.00119614 | 0.028843 | -0.6696886 |
| **Cdh12** | 0.44677543 | 1.36299043 | 6.22716342 | 0.00014356 | 0.01397668 | 1.36770315 |
| **Pgr15l** | 0.44545044 | 1.36173921 | 3.88165765 | 0.00360926 | 0.05046981 | -1.5846808 |
| **Psg16** | 0.44307068 | 1.35949484 | 4.75913801 | 0.00098564 | 0.02729468 | -0.5220345 |
| **Glra3** | 0.44247742 | 1.35893591 | 7.39790137 | 3.77E-05 | 0.01161523 | 2.72074822 |
| **Ifit2** | 0.44087366 | 1.3574261 | 6.46806628 | 0.00010758 | 0.01356866 | 1.61593378 |
| **Scn2a** | 0.43965247 | 1.35627758 | 5.24662579 | 0.00050276 | 0.02009114 | 0.07713609 |
| **Scai** | 0.43771292 | 1.35445543 | 9.8103347 | 3.72E-06 | 0.00870145 | 4.95041462 |
| **Cbln2** | 0.43755077 | 1.3543032 | 4.62277082 | 0.00119723 | 0.028843 | -0.6564192 |
| **Csmd1** | 0.43471806 | 1.35164666 | 5.35133108 | 0.00043701 | 0.01931297 | 0.34435372 |

| | | | | | | |
|---|---|---|---|---|---|---|
| **Slitrk6** | 0.43299141 | 1.35002995 | 3.27461908 | 0.00940606 | 0.08159578 | -2.3851556 |
| **Runx1t1** | 0.43049582 | 1.34769667 | 3.63726486 | 0.00527957 | 0.06100089 | -2.0788555 |
| **Bdnf** | 0.42837973 | 1.34572137 | 3.7096638 | 0.00471326 | 0.05777243 | -1.8262039 |
| **Zscan18** | 0.42340398 | 1.34108807 | 4.35245089 | 0.00177443 | 0.03491705 | -0.930322 |
| **Ccdc106** | 0.4207286 | 1.33860342 | 3.66850207 | 0.00502693 | 0.05966321 | -1.8348018 |
| **Csrnp3** | 0.41504111 | 1.33333667 | 8.7524364 | 9.64E-06 | 0.00983403 | 4.06169237 |
| **Vwc2l** | 0.41495452 | 1.33325665 | 6.82249902 | 7.13E-05 | 0.01235543 | 2.11120428 |
| **Cntnap5a** | 0.41375235 | 1.33214613 | 6.80399707 | 7.28E-05 | 0.01240261 | 2.08597396 |
| **Il1rapl1** | 0.41067662 | 1.32930911 | 6.45825867 | 0.00010883 | 0.01356866 | 1.60269448 |
| **Fam19a2** | 0.41017485 | 1.32884686 | 7.90685245 | 2.21E-05 | 0.01130325 | 3.25250371 |
| **Dlgap2** | 0.40487262 | 1.32397201 | 5.26018099 | 0.00049367 | 0.02001271 | 0.27229174 |
| **Klhl1** | 0.40399604 | 1.32316781 | 3.74988964 | 0.00442656 | 0.05587535 | -1.8046055 |
| **Pcdhgc4** | 0.40198656 | 1.32132609 | 5.08374915 | 0.00062717 | 0.02192565 | 0.00449863 |
| **Sfrp2** | 0.40194011 | 1.32128355 | 4.50673645 | 0.00141568 | 0.03098571 | -0.7028758 |
| **Akap5** | 0.40124127 | 1.32064369 | 4.66594336 | 0.00112541 | 0.02836303 | -0.558623 |
| **Zfhx3** | 0.39678087 | 1.31656693 | 4.32708742 | 0.00184215 | 0.03548748 | -1.1024988 |
| **Sox11** | 0.39638961 | 1.31620993 | 4.68557604 | 0.00109429 | 0.02836303 | -0.48623 |
| **Palm2** | 0.39536718 | 1.31527747 | 5.22891801 | 0.00051489 | 0.02033526 | 0.19939664 |
| **Klhl11** | 0.39514122 | 1.31507149 | 6.83240264 | 7.05E-05 | 0.01235543 | 2.13389102 |
| **Ahi1** | 0.3948566 | 1.31481206 | 4.38647441 | 0.00168771 | 0.03396369 | -1.1821952 |
| **Hs3st4** | 0.39475642 | 1.31472076 | 5.97831905 | 0.00019492 | 0.01473847 | 1.13685663 |
| **Prrt4** | 0.39422453 | 1.31423615 | 4.96882867 | 0.00073474 | 0.02383497 | -0.1557373 |
| **Rph3al** | 0.39413705 | 1.31415646 | 3.42138604 | 0.00743266 | 0.07270552 | -2.1815967 |
| **Gabre** | 0.39381798 | 1.31386585 | 3.33646721 | 0.00851522 | 0.07779018 | -2.4121085 |
| **St6gal2** | 0.3920534 | 1.31225983 | 4.0844287 | 0.0026483 | 0.04229261 | -1.3963027 |
| **1-Mar** | 0.39048591 | 1.31083483 | 3.61805197 | 0.00544154 | 0.0615944 | -2.0611316 |
| **Scn3b** | 0.39020837 | 1.31058268 | 3.43447741 | 0.00727898 | 0.07182718 | -2.5976316 |
| **Scn3a** | 0.39019826 | 1.3105735 | 3.33015201 | 0.00860202 | 0.07811765 | -2.7330655 |
| **Ndst3** | 0.38965388 | 1.31007906 | 4.33307864 | 0.00182591 | 0.03543399 | -1.0632373 |
| **Sdk1** | 0.38935381 | 1.3098066 | 3.47720271 | 0.00680008 | 0.06895764 | -2.2298963 |
| **Cacna2d1** | 0.38768862 | 1.30829566 | 4.17131808 | 0.00232331 | 0.03985782 | -1.468207 |
| **Nexmif** | 0.38670865 | 1.30740729 | 4.03331434 | 0.00286174 | 0.04391302 | -1.4679377 |
| **Ankrd34c** | 0.38525168 | 1.30608762 | 3.40672514 | 0.00760879 | 0.0735327 | -2.2894041 |
| **Frmpd3** | 0.38419066 | 1.30512742 | 6.13757822 | 0.00016012 | 0.01435693 | 1.26710353 |
| **Ccne2** | 0.38417785 | 1.30511583 | 4.31630233 | 0.00187178 | 0.03548748 | -0.9536207 |
| **Chl1** | 0.38347714 | 1.30448209 | 6.42875362 | 0.00011271 | 0.01356866 | 1.60017319 |
| **Plxna1** | 0.38331071 | 1.30433162 | 4.18862499 | 0.00226379 | 0.03928824 | -1.4482459 |
| **Rev3l** | 0.38291047 | 1.30396981 | 5.96863674 | 0.00019728 | 0.01473847 | 1.07524071 |
| **Grm5** | 0.38221618 | 1.30334243 | 5.2776134 | 0.00048225 | 0.02001271 | 0.13389587 |

| | | | | | | |
|---|---|---|---|---|---|---|
| **Kcnmb2** | 0.38049804 | 1.30179117 | 3.76689664 | 0.00431092 | 0.05499191 | -1.7163306 |
| **Galntl6** | 0.38015175 | 1.30147875 | 4.74898174 | 0.00099993 | 0.02750867 | -0.4078813 |
| **Trhde** | 0.37873728 | 1.30020336 | 5.14256322 | 0.00057879 | 0.02108837 | 0.12854775 |
| **Aff2** | 0.37652813 | 1.29821392 | 5.03656946 | 0.00066913 | 0.02283089 | -0.0345343 |
| **Minar2** | 0.37608178 | 1.29781234 | 4.30232553 | 0.00191094 | 0.03580669 | -1.2485581 |
| **Igsf10** | 0.37491111 | 1.29675965 | 4.6560173 | 0.00114151 | 0.02848064 | -0.5074897 |
| **Slitrk4** | 0.37366325 | 1.2956385 | 6.04825488 | 0.00017872 | 0.01459793 | 1.23196717 |
| **Dnah1** | 0.37338101 | 1.29538506 | 3.3772711 | 0.0079759 | 0.075291 | -2.2379675 |
| **Rac3** | 0.3732608 | 1.29527713 | 3.65882584 | 0.0051038 | 0.060329 | -2.0742021 |
| **Slco5a1** | 0.373091 | 1.29512468 | 3.86173209 | 0.00372186 | 0.0510717 | -1.5757546 |
| **Gprin3** | 0.37202166 | 1.29416509 | 3.94835203 | 0.00325783 | 0.04773245 | -1.4661789 |
| **Syt16** | 0.36978741 | 1.29216241 | 4.89528469 | 0.00081389 | 0.02511028 | -0.2410421 |
| **St8sia6** | 0.36878157 | 1.29126183 | 3.84403694 | 0.00382497 | 0.05176904 | -1.6087178 |
| **Cers6** | 0.36840015 | 1.2909205 | 6.34708545 | 0.00012424 | 0.01356866 | 1.58080998 |
| **Arhgap36** | 0.36767186 | 1.29026899 | 4.56211862 | 0.00130653 | 0.02989594 | -0.8395013 |
| **Lrrc7** | 0.36502175 | 1.28790105 | 4.10724563 | 0.00255854 | 0.04164652 | -1.2748505 |
| **Cbln4** | 0.36430724 | 1.28726336 | 3.36187444 | 0.00817511 | 0.07612388 | -2.5508854 |
| **Erc2** | 0.36358478 | 1.28661889 | 5.37502722 | 0.00042346 | 0.01918467 | 0.291248 |
| **Tusc1** | 0.36347485 | 1.28652086 | 3.72678712 | 0.0045889 | 0.05701593 | -1.7745435 |
| **Slc13a5** | 0.36321579 | 1.28628986 | 4.21024682 | 0.0021917 | 0.03874292 | -1.091421 |
| **Rasgrf2** | 0.36302577 | 1.28612046 | 7.25901783 | 4.38E-05 | 0.01161523 | 2.56783885 |
| **Hmgb3** | 0.36092245 | 1.28424678 | 5.62481847 | 0.00030526 | 0.01691691 | 0.70632164 |
| **Tacr1** | 0.35938405 | 1.28287807 | 3.77254726 | 0.0042732 | 0.05469122 | -1.7566995 |
| **Prkar2b** | 0.35845519 | 1.28205237 | 5.97981042 | 0.00019455 | 0.01473847 | 1.09390269 |
| **Kalrn** | 0.35822627 | 1.28184895 | 6.55742117 | 9.68E-05 | 0.01356866 | 1.7891512 |
| **Nhs** | 0.35673988 | 1.28052896 | 4.91443221 | 0.00079244 | 0.02479356 | -0.1999439 |
| **Gsg1l** | 0.35640543 | 1.28023213 | 3.89519799 | 0.0035348 | 0.04988361 | -1.8235517 |
| **Cgref1** | 0.35561071 | 1.27952711 | 5.63867725 | 0.00029984 | 0.01691691 | 0.74853678 |
| **Hook1** | 0.35502554 | 1.27900823 | 5.47474665 | 0.0003712 | 0.01839444 | 0.48175393 |
| **Htr1b** | 0.35492602 | 1.27892 | 5.62051371 | 0.00030696 | 0.01691691 | 0.73038195 |
| **Ccdc112** | 0.35423079 | 1.27830384 | 3.97712106 | 0.00311758 | 0.04647897 | -1.4429678 |
| **Usp29** | 0.35364936 | 1.27778877 | 4.97749344 | 0.00072598 | 0.02377262 | -0.2559327 |
| **Lrfn5** | 0.35345342 | 1.27761523 | 6.48104994 | 0.00010594 | 0.01356866 | 1.70965575 |
| **Lhx4** | 0.35342943 | 1.27759399 | 3.98807891 | 0.00306585 | 0.04591701 | -1.4356991 |
| **Hs6st3** | 0.35119625 | 1.2756179 | 4.63113031 | 0.00118295 | 0.02882844 | -0.5332756 |
| **Alk** | 0.35115789 | 1.27558398 | 3.40241906 | 0.00766134 | 0.07382904 | -2.3209601 |
| **Hmgb2** | 0.35105414 | 1.27549225 | 4.54186173 | 0.00134537 | 0.03043199 | -0.667697 |
| **Pnma3** | 0.34952187 | 1.27413829 | 3.72153953 | 0.00462664 | 0.05728612 | -2.06582 |
| **Bend4** | 0.3488528 | 1.27354752 | 4.86392403 | 0.00085039 | 0.02565469 | -0.2390294 |

| | | | | | | |
|---|---|---|---|---|---|---|
| **Oprk1** | 0.34576269 | 1.27082264 | 3.681002 | 0.00492942 | 0.05899912 | -1.8710486 |
| **Frmpd4** | 0.3452084 | 1.27033447 | 5.61242828 | 0.00031019 | 0.01703916 | 0.66433741 |
| **Kcnb1** | 0.34517383 | 1.27030403 | 7.87328686 | 2.29E-05 | 0.01130325 | 3.21984669 |
| **Asxl3** | 0.34486071 | 1.27002836 | 3.82840567 | 0.00391855 | 0.05232182 | -1.7421916 |
| **Lama3** | 0.34445306 | 1.26966955 | 3.27468327 | 0.00940508 | 0.08159578 | -2.4140207 |
| **Fstl5** | 0.34284432 | 1.26825454 | 5.53994418 | 0.00034083 | 0.01786654 | 0.54544038 |
| **Cdh7** | 0.3426002 | 1.26803995 | 3.79424371 | 0.00413159 | 0.05391497 | -1.7977598 |
| **Smarca1** | 0.34246526 | 1.26792135 | 3.46594507 | 0.00692298 | 0.06949355 | -2.3296644 |
| **Lingo2** | 0.3421436 | 1.26763869 | 5.83803851 | 0.00023242 | 0.01549252 | 0.99238174 |
| **Klf12** | 0.3401922 | 1.26592523 | 4.2500822 | 0.00206519 | 0.03724853 | -1.2212713 |
| **Srsf12** | 0.33994842 | 1.26571134 | 5.36398467 | 0.00042972 | 0.01931297 | 0.38623961 |
| **Dbp** | 0.33986637 | 1.26563935 | 5.77858863 | 0.00025062 | 0.01600944 | 0.87472483 |
| **Kcnh7** | 0.33985975 | 1.26563355 | 7.26935107 | 4.33E-05 | 0.01161523 | 2.59650481 |
| **Uggt2** | 0.33862825 | 1.26455366 | 6.3792556 | 0.00011955 | 0.01356866 | 1.62172836 |
| **Kcnk9** | 0.33833339 | 1.26429523 | 6.3961297 | 0.00011717 | 0.01356866 | 1.55608164 |
| **Ildr2** | 0.33708516 | 1.26320182 | 4.58067604 | 0.001272 | 0.02958149 | -0.8137681 |
| **Peg10** | 0.33550717 | 1.26182092 | 4.72247913 | 0.00103827 | 0.02788671 | -0.6688751 |
| **Pak3** | 0.33414125 | 1.26062681 | 8.94136403 | 8.08E-06 | 0.00973289 | 4.23618179 |
| **Rab27b** | 0.33379481 | 1.26032412 | 7.60781314 | 3.02E-05 | 0.01130325 | 2.95381812 |
| **Hectd2** | 0.3324492 | 1.25914916 | 4.60511587 | 0.001228 | 0.0290718 | -0.5634068 |
| **Lmo3** | 0.33241527 | 1.25911955 | 3.83450181 | 0.00388177 | 0.05196265 | -1.8472236 |
| **Cntn4** | 0.33092186 | 1.25781685 | 4.89477295 | 0.00081448 | 0.02511028 | -0.2864095 |
| **Tmem121b** | 0.32894924 | 1.25609818 | 3.52819208 | 0.00627131 | 0.06605828 | -2.1339778 |
| **Nrg3** | 0.32889761 | 1.25605323 | 7.27717947 | 4.30E-05 | 0.01161523 | 2.60254512 |
| **Wscd2** | 0.3287155 | 1.25589469 | 4.38308622 | 0.00169614 | 0.03404676 | -0.9366882 |
| **Usp34** | 0.32749273 | 1.2548307 | 5.59492321 | 0.00031731 | 0.01704154 | 0.55607123 |
| **AU041133** | 0.32730331 | 1.25466596 | 3.2543402 | 0.00971878 | 0.0827347 | -2.4115641 |
| **Wnk3** | 0.32633868 | 1.25382733 | 4.14167872 | 0.00242913 | 0.04063978 | -1.3869689 |
| **Cdh6** | 0.32619545 | 1.25370286 | 7.24785624 | 4.44E-05 | 0.01161523 | 2.57840782 |
| **Lor** | 0.32606921 | 1.25359316 | 3.57142359 | 0.0058567 | 0.0640773 | -1.9826443 |
| **Myt1l** | 0.3257145 | 1.25328498 | 5.37888034 | 0.0004213 | 0.01915025 | 0.25969837 |
| **Pcdh19** | 0.32520517 | 1.2528426 | 4.85518472 | 0.00086087 | 0.0258531 | -0.3673323 |
| **Ydjc** | 0.32516777 | 1.25281012 | 3.62148651 | 0.00541221 | 0.0615452 | -1.9202924 |
| **Tusc3** | 0.32418011 | 1.25195275 | 3.73896922 | 0.00450253 | 0.0563702 | -2.1067747 |
| **Tmem91** | 0.32359179 | 1.25144232 | 3.83439535 | 0.00388241 | 0.05196265 | -1.8044087 |
| **Grin2b** | 0.32349386 | 1.25135737 | 5.7079233 | 0.00027429 | 0.01647115 | 0.73002897 |
| **Chrm2** | 0.32166023 | 1.24976794 | 6.06616818 | 0.00017481 | 0.01447525 | 1.20460383 |
| **Cdkl5** | 0.32163218 | 1.24974363 | 7.06266327 | 5.44E-05 | 0.01210215 | 2.37440478 |
| **Serpini1** | 0.32110691 | 1.2492887 | 6.32792127 | 0.00012713 | 0.01356866 | 1.48908345 |

| | | | | | | |
|---|---|---|---|---|---|---|
| **Sorcs3** | 0.32095764 | 1.24915945 | 3.94462833 | 0.00327647 | 0.04791766 | -1.6622796 |
| **Arhgap26** | 0.31914422 | 1.24759028 | 6.34911443 | 0.00012394 | 0.01356866 | 1.52681706 |
| **Stag3** | 0.31876284 | 1.24726052 | 3.61147985 | 0.00549814 | 0.06202051 | -1.9210486 |
| **Guf1** | 0.31781504 | 1.24644139 | 7.49911472 | 3.38E-05 | 0.01161523 | 2.83852894 |
| **Zdhhc2** | 0.31764435 | 1.24629392 | 4.21237383 | 0.00218474 | 0.03871493 | -1.365407 |
| **Ikzf4** | 0.31738545 | 1.24607028 | 5.01529143 | 0.00068903 | 0.02310112 | -0.0290686 |
| **Zdhhc15** | 0.31731921 | 1.24601307 | 3.67069222 | 0.0050097 | 0.05965428 | -1.8903467 |
| **Arfgef3** | 0.31690681 | 1.24565695 | 6.36816307 | 0.00012114 | 0.01356866 | 1.53798357 |
| **Nav3** | 0.31667194 | 1.24545417 | 6.34918187 | 0.00012393 | 0.01356866 | 1.55621767 |
| **Pkia** | 0.31540243 | 1.24435871 | 5.85915767 | 0.00022631 | 0.01533545 | 0.89649609 |
| **Eid2b** | 0.31496184 | 1.24397875 | 4.98359003 | 0.00071988 | 0.02366476 | -0.0893082 |
| **Cdh13** | 0.31400233 | 1.24315168 | 4.04116806 | 0.0028278 | 0.04359044 | -1.6628122 |
| **Dync2h1** | 0.31306858 | 1.24234733 | 6.22342688 | 0.00014421 | 0.01397668 | 1.43377815 |
| **Fam84a** | 0.31278102 | 1.24209973 | 4.72010075 | 0.00104178 | 0.02788671 | -0.5408637 |
| **Olfm3** | 0.31188954 | 1.24133245 | 7.05778422 | 5.47E-05 | 0.01210215 | 2.36658246 |
| **Lin28b** | 0.31044887 | 1.24009348 | 4.39993552 | 0.00165466 | 0.03374151 | -0.8636761 |
| **Dyrk2** | 0.31022973 | 1.23990512 | 3.99849191 | 0.00301754 | 0.045541 | -1.5783406 |
| **Fam196a** | 0.31000103 | 1.23970859 | 4.14045348 | 0.00243362 | 0.04064334 | -1.2263716 |
| **Cdh18** | 0.30906608 | 1.23890544 | 5.84423099 | 0.00023061 | 0.01543277 | 0.97578875 |
| **Mchr1** | 0.30842893 | 1.23835842 | 4.79812369 | 0.0009328 | 0.02667926 | -0.3145797 |
| **Hjurp** | 0.30673326 | 1.23690376 | 3.4607298 | 0.0069807 | 0.06990651 | -2.3310872 |
| **Zfp618** | 0.30665789 | 1.23683915 | 5.68720229 | 0.00028168 | 0.01649839 | 0.80972672 |
| **Rps6ka6** | 0.30607535 | 1.23633983 | 3.625422 | 0.0053788 | 0.06145488 | -1.9789531 |
| **Gprasp2** | 0.30597776 | 1.2362562 | 5.2156929 | 0.00052416 | 0.02043832 | 0.02614081 |
| **Gabrg2** | 0.30585869 | 1.23615418 | 4.21746631 | 0.00216817 | 0.03858958 | -1.4031701 |
| **Fstl4** | 0.30569182 | 1.2360112 | 4.17298895 | 0.00231749 | 0.03981954 | -1.1419163 |
| **Cacna1e** | 0.30555349 | 1.2358927 | 4.79873884 | 0.00093199 | 0.02667926 | -0.548532 |
| **Cxxc4** | 0.30521923 | 1.23560638 | 4.06310639 | 0.00273521 | 0.04291687 | -1.5761257 |
| **Peg3** | 0.30487838 | 1.23531449 | 6.66437016 | 8.55E-05 | 0.01356866 | 1.8743768 |
| **Pgr** | 0.3026716 | 1.23342637 | 5.1575898 | 0.00056708 | 0.02088383 | 0.13302292 |
| **Magi3** | 0.30225805 | 1.23307286 | 8.48093543 | 1.25E-05 | 0.00983403 | 3.80945935 |
| **Ptprz1** | 0.30184475 | 1.23271966 | 5.96111671 | 0.00019914 | 0.01473847 | 1.01130274 |
| **Garnl3** | 0.3017423 | 1.23263213 | 5.33727447 | 0.00044527 | 0.01931297 | 0.24505481 |
| **Nbea** | 0.30106395 | 1.23205269 | 6.51109186 | 0.00010225 | 0.01356866 | 1.6994595 |
| **Gprin2** | 0.3010286 | 1.2320225 | 3.36487477 | 0.00813588 | 0.07600746 | -2.4161141 |
| **Rbm3** | 0.301007 | 1.23200405 | 6.81623302 | 7.18E-05 | 0.01235543 | 2.08267903 |
| **Acsl4** | 0.3007985 | 1.23182601 | 5.90470279 | 0.00021371 | 0.01473847 | 0.97588723 |
| **Adarb2** | 0.30064305 | 1.2316933 | 3.77566966 | 0.00425251 | 0.05462771 | -1.9346471 |
| **Plekha7** | 0.3000335 | 1.231173 | 4.18599414 | 0.00227273 | 0.03935041 | -1.2900053 |

| | | | | | | |
|---|---|---|---|---|---|---|
| **Basp1** | 0.29988693 | 1.23104792 | 3.91860364 | 0.00340989 | 0.04885679 | -1.9039887 |
| **Ica1l** | 0.29975212 | 1.2309329 | 3.40973258 | 0.0075723 | 0.07336973 | -2.448885 |
| **Nlgn1** | 0.29905053 | 1.23033444 | 6.29766216 | 0.00013183 | 0.01372423 | 1.50069849 |
| **L3mbtl1** | 0.29815804 | 1.22957356 | 4.08742322 | 0.00263634 | 0.04221155 | -1.4052311 |
| **Rwdd2a** | 0.29796972 | 1.22941307 | 6.00506771 | 0.00018854 | 0.01473847 | 1.16804018 |
| **Nos1ap** | 0.29693638 | 1.2285328 | 3.96207155 | 0.00319013 | 0.04716002 | -1.5650702 |
| **Slc17a6** | 0.29677436 | 1.22839484 | 5.35886331 | 0.00043265 | 0.01931297 | 0.22146853 |
| **Cnksr2** | 0.29658212 | 1.22823117 | 4.78143396 | 0.00095504 | 0.02685043 | -0.3802297 |
| **H2afy2** | 0.29651704 | 1.22817577 | 5.75595432 | 0.00025795 | 0.0162317 | 0.89205569 |
| **Dmxl2** | 0.29649203 | 1.22815447 | 4.80980445 | 0.00091757 | 0.02649637 | -0.5469588 |
| **Kbtbd3** | 0.29614677 | 1.22786059 | 3.68665764 | 0.00488596 | 0.0586401 | -1.8568299 |
| **Zc3h6** | 0.29574713 | 1.22752051 | 3.73934778 | 0.00449987 | 0.0563702 | -1.7770648 |
| **Vstm2a** | 0.29473102 | 1.22665626 | 5.12484806 | 0.00059292 | 0.02131998 | -0.0715963 |
| **Slc4a10** | 0.29443497 | 1.22640456 | 3.84077839 | 0.00384428 | 0.05187869 | -1.9914304 |
| **Usp9x** | 0.2938111 | 1.22587434 | 4.02099389 | 0.00291587 | 0.04450062 | -1.7267255 |
| **Dcaf7** | 0.29318653 | 1.22534375 | 5.40166374 | 0.00040876 | 0.01909522 | 0.27827675 |
| **Brinp3** | 0.29245291 | 1.22472081 | 4.93916504 | 0.00076563 | 0.02433804 | -0.2262385 |
| **Pgm2l1** | 0.29230761 | 1.22459747 | 9.37104344 | 5.47E-06 | 0.00870145 | 4.61711262 |
| **Fam19a1** | 0.29194989 | 1.22429387 | 3.51364439 | 0.00641764 | 0.06707193 | -2.2211767 |
| **Galnt13** | 0.29188767 | 1.22424107 | 6.56685202 | 9.58E-05 | 0.01356866 | 1.81549282 |
| **St8sia3** | 0.29164737 | 1.22403717 | 6.99416037 | 5.87E-05 | 0.01210215 | 2.27570162 |
| **Vps13c** | 0.2910615 | 1.2235402 | 4.69019881 | 0.0010871 | 0.02836303 | -0.6977825 |
| **Herc1** | 0.29074928 | 1.22327543 | 5.30678348 | 0.00046377 | 0.01970036 | 0.15938894 |
| **Cdkl2** | 0.28991967 | 1.2225722 | 4.63964265 | 0.0011686 | 0.02863473 | -0.6785251 |
| **Dcaf12l1** | 0.28964709 | 1.22234123 | 3.68957688 | 0.00486368 | 0.05850294 | -2.0855565 |
| **Ccdc177** | 0.28927854 | 1.22202902 | 4.15492143 | 0.00238123 | 0.04041084 | -1.1785763 |
| **Gpr68** | 0.28828684 | 1.22118929 | 4.10770974 | 0.00255675 | 0.04164652 | -1.2501815 |
| **Cdo1** | 0.28811712 | 1.22104564 | 5.63525333 | 0.00030117 | 0.01691691 | 0.73490817 |
| **Rundc3b** | 0.28777833 | 1.22075893 | 5.64509349 | 0.00029737 | 0.01691691 | 0.71252978 |
| **Myo16** | 0.28742735 | 1.22046197 | 3.60338553 | 0.0055687 | 0.06252363 | -2.2926114 |
| **Homer2** | 0.28714049 | 1.22021933 | 4.22082818 | 0.00215731 | 0.03845755 | -1.2438485 |
| **Sstr3** | 0.28706401 | 1.22015464 | 3.29025069 | 0.00917214 | 0.08067212 | -2.446349 |
| **Tmx4** | 0.28463141 | 1.21809902 | 8.51574016 | 1.21E-05 | 0.00983403 | 3.84027497 |
| **Prkaa2** | 0.28375881 | 1.21736248 | 6.5992698 | 9.22E-05 | 0.01356866 | 1.87273177 |
| **Cep192** | 0.28361025 | 1.21723713 | 4.78952015 | 0.0009442 | 0.02680628 | -0.3338105 |
| **Zfp783** | 0.28152875 | 1.21548219 | 3.5224518 | 0.00632862 | 0.0664959 | -2.2195965 |
| **Pclo** | 0.28126249 | 1.21525788 | 5.4613499 | 0.0003778 | 0.01839444 | 0.37254855 |
| **Gpc6** | 0.28071662 | 1.21479815 | 3.31317131 | 0.00884001 | 0.07940008 | -2.3610827 |
| **Cdh24** | 0.2806711 | 1.21475982 | 3.5057779 | 0.00649825 | 0.0675209 | -2.3742976 |

| | | | | | | |
|---|---|---|---|---|---|---|
| **Lypd6b** | 0.28056948 | 1.21467426 | 3.38101622 | 0.00792822 | 0.07520578 | -2.3321641 |
| **Pgap1** | 0.27979492 | 1.21402229 | 4.65002661 | 0.00115134 | 0.02848064 | -0.7246632 |
| **Slc44a5** | 0.27964479 | 1.21389597 | 3.68630721 | 0.00488864 | 0.0586401 | -1.8791795 |
| **Htr2a** | 0.27962134 | 1.21387624 | 4.57986131 | 0.00127349 | 0.02958149 | -0.6413817 |
| **Atad5** | 0.27925081 | 1.21356452 | 3.38779043 | 0.00784272 | 0.07480745 | -2.2357842 |
| **Sfxn1** | 0.27856175 | 1.21298504 | 3.54741184 | 0.00608331 | 0.06544692 | -2.3645421 |
| **Oprl1** | 0.2784728 | 1.21291025 | 5.84602424 | 0.00023009 | 0.01543277 | 0.90971893 |
| **Mcph1** | 0.27840184 | 1.21285059 | 4.81424272 | 0.00091185 | 0.02640938 | -0.2886391 |
| **Zkscan2** | 0.27792631 | 1.21245088 | 5.90315953 | 0.00021412 | 0.01473847 | 0.99883171 |
| **Bhlhb9** | 0.27789931 | 1.2124282 | 4.39517971 | 0.00166626 | 0.03385518 | -1.0810031 |
| **Shisa7** | 0.27788554 | 1.21241663 | 3.48140025 | 0.00675484 | 0.06878842 | -2.5017555 |
| **Aff4** | 0.27766549 | 1.21223171 | 6.38272425 | 0.00011906 | 0.01356866 | 1.54220142 |
| **Bicd1** | 0.2776571 | 1.21222466 | 5.26853394 | 0.00048816 | 0.02001271 | 0.14606694 |
| **Nabp1** | 0.27734249 | 1.21196034 | 3.57075837 | 0.00586286 | 0.0640773 | -2.0264451 |
| **Ncam2** | 0.27688238 | 1.21157388 | 8.3294953 | 1.45E-05 | 0.00983403 | 3.6646617 |
| **Psd3** | 0.27665584 | 1.21138365 | 6.19117972 | 0.00014997 | 0.01397668 | 1.31037944 |
| **Lrrn3** | 0.27647754 | 1.21123394 | 4.39063315 | 0.00167743 | 0.03391862 | -1.0515027 |
| **Klhl15** | 0.27604305 | 1.21086922 | 4.89279038 | 0.00081673 | 0.02513254 | -0.1985511 |
| **Gria2** | 0.27583448 | 1.21069418 | 6.44728686 | 0.00011026 | 0.01356866 | 1.61769681 |
| **Grik2** | 0.27474297 | 1.20977854 | 3.47764916 | 0.00679525 | 0.06895612 | -2.5197858 |
| **Coil** | 0.2747278 | 1.20976582 | 4.29257944 | 0.00193875 | 0.03600787 | -1.0184134 |
| **Pou3f2** | 0.27387437 | 1.20905039 | 3.76014762 | 0.00435642 | 0.05533773 | -1.9624107 |
| **Sntg1** | 0.2734742 | 1.20871507 | 5.00776986 | 0.00069622 | 0.0232037 | -0.0669151 |
| **Slc1a6** | 0.27293506 | 1.20826345 | 3.94291792 | 0.00328506 | 0.04796476 | -1.4916001 |
| **Spats2l** | 0.27281472 | 1.20816267 | 3.30087358 | 0.00901662 | 0.07991985 | -2.7145262 |
| **Rnf152** | 0.27240285 | 1.20781781 | 4.27565391 | 0.00198809 | 0.03648229 | -1.2765429 |
| **Negr1** | 0.27201738 | 1.20749514 | 3.56619854 | 0.00590525 | 0.06416633 | -2.4390458 |
| **Dync2li1** | 0.27191503 | 1.20740947 | 3.47211814 | 0.0068553 | 0.06914346 | -2.1847534 |
| **Rimklb** | 0.27187096 | 1.20737259 | 7.02731056 | 5.66E-05 | 0.01210215 | 2.32782954 |
| **Lrrtm3** | 0.27078984 | 1.20646816 | 3.54290448 | 0.00612686 | 0.06556265 | -2.2876092 |
| **Htr2c** | 0.2704207 | 1.2061595 | 5.25067533 | 0.00050002 | 0.02009114 | 0.08245448 |
| **Gjd2** | 0.26885445 | 1.20485076 | 5.03084446 | 0.00067442 | 0.02284415 | -0.1056407 |
| **Flrt3** | 0.26861284 | 1.204649 | 3.25609062 | 0.00969136 | 0.08264749 | -2.6034194 |
| **Fat3** | 0.26817194 | 1.2042809 | 5.34276023 | 0.00044203 | 0.01931297 | 0.28220392 |
| **Mdga1** | 0.26809351 | 1.20421543 | 3.79383727 | 0.0041342 | 0.05391497 | -2.018749 |
| **Kcnma1** | 0.26780616 | 1.2039756 | 5.97960249 | 0.0001946 | 0.01473847 | 1.05898516 |
| **Eif1a** | 0.26743982 | 1.20366992 | 6.19483573 | 0.00014931 | 0.01397668 | 1.38878574 |
| **Rgs17** | 0.26646535 | 1.20285717 | 7.67391701 | 2.81E-05 | 0.01130325 | 3.00344235 |
| **Grm8** | 0.26627067 | 1.20269487 | 4.04206363 | 0.00282395 | 0.04359044 | -1.4243629 |

| | | | | | | |
|---|---|---|---|---|---|---|
| **N4bp2** | 0.26575338 | 1.20226371 | 5.21226948 | 0.00052659 | 0.02043832 | 0.16620383 |
| **Prr16** | 0.26473716 | 1.20141714 | 3.59306956 | 0.00565999 | 0.06307646 | -1.9947192 |
| **Syt14** | 0.26473619 | 1.20141634 | 4.32335432 | 0.00185235 | 0.03548748 | -0.9829713 |
| **Cspg5** | 0.26454549 | 1.20125755 | 3.74018945 | 0.00449397 | 0.056364 | -2.1723545 |
| **Epha5** | 0.26385419 | 1.20068207 | 6.04613931 | 0.00017918 | 0.01459793 | 1.14834619 |
| **Dpp10** | 0.26355895 | 1.20043638 | 6.094577 | 0.00016879 | 0.01435693 | 1.21646797 |
| **Dgkh** | 0.26278534 | 1.19979285 | 3.56494725 | 0.00591694 | 0.06421059 | -2.2518899 |
| **Shisa6** | 0.26265477 | 1.19968427 | 3.61484275 | 0.0054691 | 0.06181701 | -2.1237112 |
| **Chml** | 0.26192252 | 1.19907552 | 5.3568002 | 0.00043384 | 0.01931297 | 0.35367359 |
| **Mcm2** | 0.26140017 | 1.19864145 | 3.25841535 | 0.00965508 | 0.08250591 | -2.413838 |
| **Cntln** | 0.26081923 | 1.19815888 | 4.28970005 | 0.00194705 | 0.03606644 | -1.0323388 |
| **Plcxd2** | 0.26047804 | 1.19787555 | 6.45066593 | 0.00010982 | 0.01356866 | 1.62927207 |
| **Tanc2** | 0.26032402 | 1.19774768 | 5.62168923 | 0.00030649 | 0.01691691 | 0.58689854 |
