## Supplementary material for "The ultrasonic vocalization (USV) syllable profile during neonatal opioid withdrawal and a kappa opioid receptor component to increased USV emissions in female mice": Table.S3.pdf

**Table S3.** Downregulated genes ($log_2FC \leq -0.26$; $p < 0.01$) between morphine (2F, 2M) vs. saline (2F, 2M) mice.

| Gene | logFC | Fold Change | t value | p value | Adjusted p value | B |
|---|---|---|---|---|---|---|
| **Gpr143** | -3.3571378 | 0.09758899 | -4.524874 | 0.0013789 | 0.0307147 | -3.960946 |
| **Ltf** | -2.6636728 | 0.15781729 | -3.3367178 | 0.00851179 | 0.07779018 | -3.7906347 |
| **Camp** | -2.6352947 | 0.16095232 | -4.1441409 | 0.00242015 | 0.0405297 | -3.6458536 |
| **S100a9** | -1.9674669 | 0.25570161 | -3.2690685 | 0.0094906 | 0.08199257 | -3.135056 |
| **Tgm7** | -1.8681149 | 0.27393112 | -3.865874 | 0.00369815 | 0.0510717 | -3.768619 |
| **Krt15** | -1.7697924 | 0.29325094 | -3.5046482 | 0.00650991 | 0.06755883 | -3.6168594 |
| **Cyp2d26** | -1.7183377 | 0.30389868 | -3.2726329 | 0.00943622 | 0.08172269 | -3.9464428 |
| **Pax1** | -1.6146659 | 0.32654056 | -3.4565835 | 0.00702695 | 0.07012685 | -3.7570632 |
| **Ctse** | -1.5441329 | 0.34290174 | -3.4424177 | 0.00718738 | 0.07121505 | -3.5860921 |
| **Tspan10** | -1.3119704 | 0.40277042 | -3.5870704 | 0.00571381 | 0.06342905 | -3.4901338 |
| **Lcn2** | -1.2665028 | 0.41566616 | -3.5345792 | 0.00620817 | 0.06586461 | -3.2300603 |
| **Haao** | -1.2205709 | 0.42911286 | -3.6344217 | 0.00530322 | 0.06110777 | -3.2662755 |
| **Magix** | -1.1649819 | 0.44596984 | -4.2030948 | 0.00221527 | 0.03888574 | -2.463086 |
| **Etnppl** | -1.1627566 | 0.44665828 | -7.6920965 | 2.76E-05 | 0.01130325 | 2.41767338 |
| **Acsm3** | -1.1293895 | 0.45710911 | -3.7106653 | 0.00470589 | 0.05775851 | -3.1423013 |
| **Nkx2-9** | -1.0818091 | 0.47243604 | -4.7870729 | 0.00094746 | 0.02683412 | -1.6343916 |
| **Serpinb1a** | -1.0004667 | 0.49983828 | -7.0491202 | 5.52E-05 | 0.01210215 | 1.01561393 |
| **Abca4** | -0.9566639 | 0.51524701 | -4.5094375 | 0.00141014 | 0.0309643 | -0.8147917 |
| **Slc5a11** | -0.9386182 | 0.52173235 | -9.1439489 | 6.71E-06 | 0.00870145 | 3.44256478 |
| **Dao** | -0.9295681 | 0.52501548 | -4.6871318 | 0.00109187 | 0.02836303 | -0.5642777 |
| **Rec8** | -0.9250787 | 0.52665178 | -3.2775088 | 0.00936235 | 0.0814689 | -2.8593138 |
| **Galnt15** | -0.9138779 | 0.53075652 | -3.4561491 | 0.00703181 | 0.07012685 | -2.1497607 |
| **Trac** | -0.9102252 | 0.53210204 | -4.6072652 | 0.00122421 | 0.02903664 | -1.449892 |
| **Lyz2** | -0.8895949 | 0.53976567 | -8.6542889 | 1.06E-05 | 0.00983403 | 2.58234854 |
| **Adamtsl4** | -0.8667139 | 0.54839452 | -10.370411 | 2.33E-06 | 0.00870145 | 4.69316984 |
| **Tmem52** | -0.843777 | 0.55718296 | -3.3461386 | 0.00838405 | 0.07727863 | -2.5931261 |
| **Aqp6** | -0.8205953 | 0.56620827 | -4.3109983 | 0.00188654 | 0.03554702 | -1.1401644 |
| **Ccl9** | -0.8175655 | 0.56739861 | -5.4059087 | 0.00040647 | 0.01904097 | -0.5223662 |
| **Mfap4** | -0.8157006 | 0.56813253 | -5.1184415 | 0.00059813 | 0.02140612 | -0.1853815 |
| **A2m** | -0.8064813 | 0.57177471 | -5.4498021 | 0.00038359 | 0.01842957 | 0.51601639 |
| **Abca8a** | -0.7910806 | 0.57791107 | -4.8559102 | 0.00085999 | 0.0258531 | -0.699545 |
| **Mybpc3** | -0.7587873 | 0.59099292 | -3.3130234 | 0.00884211 | 0.07940008 | -2.9493192 |
| **Aspg** | -0.7563667 | 0.59198531 | -6.5043882 | 0.00010306 | 0.01356866 | 1.35071093 |
| **AC125101.1** | -0.7559012 | 0.59217637 | -5.7296902 | 0.00026675 | 0.016358 | -0.1911361 |
| **Atoh8** | -0.7444669 | 0.59688839 | -4.7144966 | 0.00105012 | 0.02793253 | -1.1719424 |
| **Enpp3** | -0.7213825 | 0.60651595 | -4.6722126 | 0.00111537 | 0.02836303 | -1.0765744 |

| | | | | | | |
|---|---|---|---|---|---|---|
| **Calr4** | -0.7209498 | 0.60669788 | -3.3088902 | 0.00890108 | 0.07949593 | -2.4531409 |
| **Lag3** | -0.7191496 | 0.60745542 | -7.2507754 | 4.42E-05 | 0.01161523 | 1.4933655 |
| **Ndrg1** | -0.714231 | 0.60952996 | -9.7543909 | 3.91E-06 | 0.00870145 | 4.93589763 |
| **Serping1** | -0.7065836 | 0.61276949 | -6.0647915 | 0.0001751 | 0.01447525 | 1.03693473 |
| **Ifitm1** | -0.6970516 | 0.61683153 | -4.7236113 | 0.0010366 | 0.02788671 | -1.0704551 |
| **Pkn3** | -0.6821817 | 0.62322212 | -4.3878058 | 0.00168441 | 0.03396369 | -1.481505 |
| **H2-T24** | -0.6777094 | 0.62515706 | -4.478652 | 0.00147472 | 0.03163717 | -1.2302955 |
| **Kctd11** | -0.6699974 | 0.62850783 | -5.0997935 | 0.00061355 | 0.021705 | -0.4176866 |
| **Slc6a12** | -0.6583744 | 0.63359182 | -3.6562219 | 0.0051247 | 0.060329 | -2.1701569 |
| **Cdh3** | -0.6580511 | 0.63373382 | -3.8009527 | 0.00408881 | 0.05370223 | -1.8380581 |
| **Gfap** | -0.6539012 | 0.63555938 | -4.8692609 | 0.00084406 | 0.02561018 | -0.4731314 |
| **Pkd2l1** | -0.6471793 | 0.63852751 | -5.0295263 | 0.00067565 | 0.02284415 | -0.3353686 |
| **Plscr2** | -0.6455507 | 0.63924873 | -3.2436698 | 0.00988763 | 0.08366535 | -2.5695234 |
| **Arl5c** | -0.6438513 | 0.64000217 | -4.8954795 | 0.00081367 | 0.02511028 | -0.6500771 |
| **Sp100** | -0.6331984 | 0.64474544 | -4.2509945 | 0.00206239 | 0.03723777 | -1.3272971 |
| **Pgm5** | -0.6313975 | 0.6455508 | -4.4879972 | 0.00145479 | 0.03152146 | -0.7386864 |
| **Lcat** | -0.623989 | 0.64887434 | -7.0731961 | 5.38E-05 | 0.01210215 | 2.39477431 |
| **Col3a1** | -0.6194297 | 0.65092817 | -4.3935636 | 0.00167022 | 0.03387861 | -0.976894 |
| **Kcnk13** | -0.6178214 | 0.65165423 | -8.3228959 | 1.46E-05 | 0.00983403 | 3.59823815 |
| **Mt2** | -0.6108311 | 0.65481935 | -5.4370947 | 0.00039006 | 0.01863469 | 0.50167793 |
| **Slc15a3** | -0.6071819 | 0.65647779 | -3.7957222 | 0.00412212 | 0.05388796 | -2.0233063 |
| **Trf** | -0.6029902 | 0.65838792 | -7.4249529 | 3.66E-05 | 0.01161523 | 2.7602067 |
| **Dcn** | -0.6018603 | 0.65890378 | -5.3822297 | 0.00041943 | 0.01915025 | 0.43576668 |
| **Emilin2** | -0.6017204 | 0.65896767 | -3.8099749 | 0.00403201 | 0.05333006 | -1.7791912 |
| **Itgb4** | -0.5977077 | 0.66080306 | -6.4819312 | 0.00010583 | 0.01356866 | 1.70547245 |
| **Ppp1r3g** | -0.5955114 | 0.66180981 | -4.145438 | 0.00241543 | 0.0405297 | -1.4661711 |
| **Fxyd1** | -0.5884914 | 0.66503796 | -5.2603213 | 0.00049358 | 0.02001271 | 0.24334124 |
| **Llgl2** | -0.5874593 | 0.66551391 | -3.3678854 | 0.00809673 | 0.07573112 | -2.545337 |
| **Cacna2d4** | -0.5844464 | 0.6669052 | -4.5696613 | 0.00129237 | 0.02975173 | -0.6170106 |
| **Trp63** | -0.5831489 | 0.66750528 | -3.7407852 | 0.0044898 | 0.056364 | -1.922535 |
| **Dock5** | -0.5810811 | 0.66846268 | -6.1972098 | 0.00014888 | 0.01397668 | 1.3986465 |
| **Tbx1** | -0.5772812 | 0.67022567 | -4.8353519 | 0.00088517 | 0.02600619 | -0.7420657 |
| **Irgm2** | -0.5764388 | 0.67061711 | -3.2641383 | 0.00956636 | 0.08222779 | -2.4934092 |
| **C4b** | -0.5757101 | 0.67095591 | -5.9135156 | 0.00021136 | 0.01473847 | 1.04328554 |
| **Neat1** | -0.5575413 | 0.67945914 | -5.8587316 | 0.00022643 | 0.01533545 | 0.94668002 |
| **Enpp1** | -0.5542935 | 0.68099047 | -6.6251187 | 8.95E-05 | 0.01356866 | 1.82624168 |
| **Flnc** | -0.5522157 | 0.68197195 | -3.7470692 | 0.00444605 | 0.05598994 | -1.7331909 |
| **Ninj2** | -0.5509858 | 0.68255359 | -3.6451286 | 0.00521474 | 0.06076284 | -1.8860715 |
| **Gstp2** | -0.5502882 | 0.68288371 | -3.6179866 | 0.00544211 | 0.0615944 | -2.0225172 |

| | | | | | | |
|---|---|---|---|---|---|---|
| **Mmp2** | -0.5440683 | 0.68583419 | -5.3204991 | 0.00045535 | 0.01939138 | 0.33381839 |
| **Sp7** | -0.5402037 | 0.68767379 | -6.0194595 | 0.0001852 | 0.01468439 | 1.1650627 |
| **Shroom1** | -0.5355173 | 0.68991124 | -7.6122856 | 3.00E-05 | 0.01130325 | 2.61192901 |
| **Cryab** | -0.5351299 | 0.69009654 | -7.3838152 | 3.83E-05 | 0.01161523 | 2.69449461 |
| **Dct** | -0.5263877 | 0.69429095 | -3.4148933 | 0.00751013 | 0.0730818 | -2.4422189 |
| **Cd14** | -0.5256557 | 0.69464331 | -3.5625237 | 0.00593965 | 0.06441556 | -2.2194805 |
| **Anln** | -0.5180145 | 0.69833226 | -5.4903907 | 0.00036366 | 0.01832533 | 0.42561669 |
| **Prelp** | -0.5160976 | 0.69926073 | -3.4804749 | 0.00676479 | 0.0688066 | -2.1056693 |
| **Npas2** | -0.5146843 | 0.6999461 | -6.4879959 | 0.00010508 | 0.01356866 | 1.47211185 |
| **Efhd1** | -0.5059651 | 0.70418913 | -7.3369961 | 4.03E-05 | 0.01161523 | 2.67363583 |
| **Ppfibp2** | -0.5048716 | 0.70472308 | -7.6652791 | 2.84E-05 | 0.01130325 | 2.94300189 |
| **Rasal3** | -0.5014699 | 0.70638672 | -3.8353215 | 0.00387685 | 0.05196265 | -1.7745172 |
| **Csf3r** | -0.5001889 | 0.70701421 | -5.0864754 | 0.00062483 | 0.02191995 | -0.0167257 |
| **Loxl2** | -0.4983186 | 0.70793137 | -5.059957 | 0.00064796 | 0.02230057 | -0.2043813 |
| **Gjb6** | -0.4940248 | 0.71004147 | -8.1018488 | 1.82E-05 | 0.01057257 | 3.44628804 |
| **Cdkn1c** | -0.4937758 | 0.71016405 | -5.3382897 | 0.00044467 | 0.01931297 | 0.37444124 |
| **Npsr1** | -0.4906597 | 0.71169958 | -3.8657648 | 0.00369878 | 0.0510717 | -1.7360186 |
| **Ddit4l** | -0.4903749 | 0.71184007 | -3.3817062 | 0.00791947 | 0.07520578 | -2.3030041 |
| **Cfh** | -0.4887478 | 0.71264336 | -6.3189766 | 0.0001285 | 0.01362891 | 1.55768678 |
| **Tec** | -0.483416 | 0.71528198 | -3.4833986 | 0.00673341 | 0.06875199 | -2.1678152 |
| **Tmprss7** | -0.4827476 | 0.71561345 | -3.5991009 | 0.00560643 | 0.06278007 | -2.1455482 |
| **Ttyh2** | -0.4811119 | 0.71642525 | -6.0473407 | 0.00017892 | 0.01459793 | 1.14226353 |
| **Cd82** | -0.4789244 | 0.71751235 | -7.0572206 | 5.47E-05 | 0.01210215 | 2.35622061 |
| **Fbln1** | -0.4766877 | 0.71862563 | -4.1867976 | 0.00226999 | 0.03934345 | -1.1741865 |
| **Enpp2** | -0.4764909 | 0.71872365 | -4.6761609 | 0.0011091 | 0.02836303 | -0.7521617 |
| **Chad** | -0.4752582 | 0.71933806 | -3.2625756 | 0.0095905 | 0.0823328 | -2.5796108 |
| **Olfml3** | -0.4738676 | 0.72003175 | -4.2334516 | 0.00211703 | 0.03797837 | -1.0603309 |
| **Hmgcs2** | -0.4728789 | 0.72052534 | -5.2001441 | 0.0005353 | 0.0204724 | 0.20625253 |
| **Vstm4** | -0.4721279 | 0.7209005 | -5.2864007 | 0.0004766 | 0.01999711 | 0.30611101 |
| **Plin3** | -0.4715295 | 0.72119958 | -5.3792669 | 0.00042108 | 0.01915025 | 0.40130477 |
| **Col1a2** | -0.4707692 | 0.72157979 | -8.4451562 | 1.29E-05 | 0.00983403 | 3.77090209 |
| **Apod** | -0.4706378 | 0.7216455 | -6.9033319 | 6.50E-05 | 0.01210215 | 2.15286084 |
| **Loxl3** | -0.4701156 | 0.72190673 | -4.4305774 | 0.00158198 | 0.03303559 | -0.8297589 |
| **Bag3** | -0.4697032 | 0.72211313 | -4.3410714 | 0.00180448 | 0.03509886 | -1.0123734 |
| **Col1a1** | -0.4685841 | 0.72267349 | -3.8088805 | 0.00403886 | 0.05333693 | -1.7736072 |
| **Car14** | -0.4675905 | 0.72317141 | -6.5309654 | 9.99E-05 | 0.01356866 | 1.78923375 |
| **Mpeg1** | -0.4668346 | 0.72355037 | -9.4823016 | 4.95E-06 | 0.00870145 | 4.59555435 |
| **Arhgef37** | -0.4659189 | 0.72400978 | -3.3327717 | 0.0085659 | 0.07795758 | -2.3841905 |
| **Bgn** | -0.4646937 | 0.7246249 | -5.423011 | 0.00039738 | 0.01882439 | 0.47524706 |

| | | | | | | |
|---|---|---|---|---|---|---|
| **Snx33** | -0.4646799 | 0.72463184 | -6.2684561 | 0.00013656 | 0.01376723 | 1.49237259 |
| **Col2a1** | -0.4641045 | 0.72492092 | -5.2138496 | 0.00052547 | 0.02043832 | 0.08765739 |
| **Fzd10** | -0.4636937 | 0.72512734 | -3.4851529 | 0.00671466 | 0.06868376 | -2.1299795 |
| **Stat5a** | -0.4635126 | 0.72521839 | -4.2154545 | 0.0021747 | 0.03864502 | -1.1317025 |
| **Rhbdf2** | -0.4622188 | 0.72586904 | -3.6639905 | 0.00506262 | 0.05995508 | -2.0028398 |
| **Itgb2** | -0.4605302 | 0.72671912 | -3.3433403 | 0.00842179 | 0.07739782 | -2.2939657 |
| **Scd1** | -0.4590086 | 0.72748601 | -5.9035788 | 0.00021401 | 0.01473847 | 0.93705553 |
| **Mt1** | -0.4541913 | 0.72991924 | -9.8602274 | 3.57E-06 | 0.00870145 | 5.02793948 |
| **Cebpa** | -0.4527544 | 0.73064654 | -4.5906991 | 0.00125375 | 0.0294065 | -0.6059939 |
| **Exoc3l4** | -0.4519858 | 0.73103592 | -3.801194 | 0.00408728 | 0.05370223 | -1.6876738 |
| **Slpi** | -0.4499185 | 0.73208419 | -4.9597374 | 0.00074406 | 0.02403811 | -0.2061219 |
| **Litaf** | -0.4498624 | 0.73211265 | -7.618936 | 2.98E-05 | 0.01130325 | 2.95720472 |
| **Ppp1r14a** | -0.4449478 | 0.7346109 | -6.7175807 | 8.04E-05 | 0.01316315 | 1.98679372 |
| **Cntn2** | -0.4431314 | 0.7355364 | -6.1400946 | 0.00015963 | 0.01435693 | 1.24377834 |
| **Smtnl2** | -0.443129 | 0.7355376 | -4.6452559 | 0.00115923 | 0.02849755 | -0.5210042 |
| **Rbpms2** | -0.4422718 | 0.73597474 | -3.6009643 | 0.00558999 | 0.0627209 | -1.9434469 |
| **Pde8a** | -0.4421367 | 0.73604367 | -5.8482267 | 0.00022945 | 0.01543277 | 0.9628028 |
| **Hapln2** | -0.4418919 | 0.73616861 | -4.4217727 | 0.00160251 | 0.03303559 | -1.0926164 |
| **Lpar1** | -0.440216 | 0.73702427 | -10.221733 | 2.63E-06 | 0.00870145 | 5.31716666 |
| **Tmem63a** | -0.4336983 | 0.74036148 | -5.3813094 | 0.00041994 | 0.01915025 | 0.28385397 |
| **Aspa** | -0.4316507 | 0.74141298 | -4.6662446 | 0.00112493 | 0.02836303 | -0.7062273 |
| **C1qb** | -0.4309667 | 0.7417646 | -5.2540598 | 0.00049775 | 0.02009114 | 0.27277381 |
| **Pls1** | -0.4285836 | 0.74299086 | -5.3306871 | 0.0004492 | 0.01931297 | 0.37163804 |
| **Tcn2** | -0.4278471 | 0.74337025 | -7.5521565 | 3.20E-05 | 0.01161523 | 2.87989922 |
| **Gadd45g** | -0.4271917 | 0.74370803 | -5.4599572 | 0.00037849 | 0.01839444 | 0.52945121 |
| **Jup** | -0.4253917 | 0.74463653 | -10.580754 | 1.96E-06 | 0.00870145 | 5.52020489 |
| **Gng8** | -0.424026 | 0.74534174 | -4.5235824 | 0.00138148 | 0.0307147 | -0.7257596 |
| **Fam83d** | -0.4238787 | 0.74541785 | -4.5840021 | 0.00126591 | 0.02951498 | -0.6012145 |
| **Fa2h** | -0.4236101 | 0.74555668 | -6.7724243 | 7.55E-05 | 0.0127301 | 2.00132457 |
| **AC132307.1** | -0.4225761 | 0.7460912 | -4.7208353 | 0.0010407 | 0.02788671 | -0.5200291 |
| **Plekhb1** | -0.4224452 | 0.74615892 | -6.4496177 | 0.00010995 | 0.01356866 | 1.61756786 |
| **Tgm2** | -0.4198394 | 0.74750785 | -4.6263382 | 0.00119111 | 0.028843 | -0.5851108 |
| **Gjb1** | -0.4190488 | 0.74791758 | -5.9092167 | 0.0002125 | 0.01473847 | 0.97031872 |
| **Lats2** | -0.4184503 | 0.74822792 | -7.6961959 | 2.75E-05 | 0.01130325 | 2.89042467 |
| **Pdk4** | -0.4183342 | 0.74828811 | -4.6907657 | 0.00108622 | 0.02836303 | -0.450919 |
| **Icosl** | -0.4178845 | 0.7485214 | -3.7858511 | 0.00418577 | 0.05429914 | -1.7321114 |
| **Cpm** | -0.4176236 | 0.74865678 | -4.324611 | 0.00184891 | 0.03548748 | -1.188248 |
| **Unc5b** | -0.4171104 | 0.74892314 | -7.8544046 | 2.33E-05 | 0.01130325 | 3.19273212 |
| **D7Ertd443e** | -0.416309 | 0.74933929 | -4.0526144 | 0.00277908 | 0.04328868 | -1.305229 |

| | | | | | | |
|---|---|---|---|---|---|---|
| **Mal** | -0.414523 | 0.75026753 | -4.5554265 | 0.00131923 | 0.0301048 | -0.9287145 |
| **Emcn** | -0.4140623 | 0.75050716 | -4.1695475 | 0.00232949 | 0.03992326 | -1.1676427 |
| **Rnf43** | -0.412309 | 0.75141976 | -3.4143924 | 0.00751614 | 0.07309397 | -2.1997517 |
| **Mboat1** | -0.4117132 | 0.75173018 | -4.3665505 | 0.00173793 | 0.03443997 | -0.8809414 |
| **Arhgap30** | -0.4111354 | 0.75203127 | -3.8108295 | 0.00402667 | 0.05330128 | -1.6812846 |
| **Tbx3** | -0.4105868 | 0.75231732 | -3.4393334 | 0.00722282 | 0.07144022 | -2.1464565 |
| **Rtkn2** | -0.4079925 | 0.75367136 | -4.979071 | 0.00072439 | 0.0237669 | -0.094268 |
| **Lpcat2** | -0.4070541 | 0.75416177 | -3.4826759 | 0.00674115 | 0.06875495 | -2.1591959 |
| **Mbp** | -0.4053071 | 0.75507554 | -7.237913 | 4.48E-05 | 0.01161523 | 2.55327557 |
| **Srd5a1** | -0.4046609 | 0.75541384 | -5.9420059 | 0.00020395 | 0.01473847 | 1.09584285 |
| **Pygm** | -0.4043053 | 0.75560005 | -6.6482747 | 8.71E-05 | 0.01356866 | 1.87139082 |
| **Lamc3** | -0.4021594 | 0.75672477 | -4.5754676 | 0.00128159 | 0.02961326 | -0.6029664 |
| **Tshb** | -0.3989359 | 0.75841748 | -5.529664 | 0.00034544 | 0.0180354 | 0.55953974 |
| **Slc8b1** | -0.3971202 | 0.75937259 | -5.1996894 | 0.00053562 | 0.0204724 | 0.11799252 |
| **Fxyd5** | -0.3969536 | 0.75946028 | -4.1726365 | 0.00231871 | 0.03981954 | -1.1429516 |
| **AC134576.3** | -0.3958252 | 0.76005453 | -4.4678377 | 0.00149815 | 0.03190009 | -0.7683323 |
| **Sod3** | -0.3941253 | 0.76095061 | -6.6959288 | 8.24E-05 | 0.01336627 | 1.92657389 |
| **Arhgap45** | -0.3902822 | 0.76298036 | -5.1141178 | 0.00060167 | 0.02140612 | 0.04054603 |
| **Ecm2** | -0.3891167 | 0.76359696 | -3.4859978 | 0.00670565 | 0.06866063 | -2.1250826 |
| **Cavin1** | -0.3888568 | 0.76373456 | -5.1327519 | 0.00058657 | 0.02122735 | 0.06149923 |
| **Prr5l** | -0.3885436 | 0.76390038 | -5.1585623 | 0.00056633 | 0.02088383 | 0.01872549 |
| **Slc52a3** | -0.3884562 | 0.76394663 | -3.8778204 | 0.00363066 | 0.05053222 | -1.6084848 |
| **Pla2g16** | -0.38719 | 0.76461745 | -5.452623 | 0.00038216 | 0.01842957 | 0.42343108 |
| **Fam107a** | -0.3869803 | 0.7647286 | -4.7362572 | 0.00101814 | 0.02779648 | -0.5994877 |
| **Mmp28** | -0.3862015 | 0.7651415 | -3.5370516 | 0.0061839 | 0.06583672 | -2.0663275 |
| **Col20a1** | -0.384426 | 0.76608371 | -5.2436941 | 0.00050474 | 0.02009387 | 0.26187319 |
| **Rgl3** | -0.3837611 | 0.7664369 | -4.688803 | 0.00108927 | 0.02836303 | -0.5079495 |
| **Plp1** | -0.3831042 | 0.76678596 | -5.5863112 | 0.00032087 | 0.01717835 | 0.5651335 |
| **Tgfb3** | -0.381568 | 0.76760285 | -4.1024436 | 0.00257716 | 0.04182984 | -1.2374667 |
| **Cyp27a1** | -0.381271 | 0.76776089 | -5.1574736 | 0.00056717 | 0.02088383 | 0.1371129 |
| **Klhdc7a** | -0.3803687 | 0.76824123 | -4.2339017 | 0.00211561 | 0.03797837 | -1.2690578 |
| **Megf10** | -0.3801622 | 0.76835122 | -5.5217444 | 0.00034903 | 0.01811086 | 0.53081795 |
| **Kcnj10** | -0.3794683 | 0.76872086 | -4.6512351 | 0.00114935 | 0.02848064 | -0.788057 |
| **Ccdc163** | -0.379411 | 0.76875138 | -3.7401144 | 0.0044945 | 0.056364 | -1.7375939 |
| **Msn** | -0.3786642 | 0.7691494 | -5.526565 | 0.00034684 | 0.01805267 | 0.49949081 |
| **Tesk2** | -0.3779692 | 0.76952002 | -6.3356952 | 0.00012595 | 0.01356866 | 1.55515607 |
| **Btd** | -0.3779156 | 0.76954861 | -4.7936437 | 0.00093872 | 0.02674075 | -0.3295789 |
| **Nfatc1** | -0.3770937 | 0.76998715 | -4.9113833 | 0.00079582 | 0.02485304 | -0.1816535 |
| **Ctss** | -0.3761127 | 0.77051091 | -8.1814122 | 1.68E-05 | 0.01047278 | 3.51896042 |

| | | | | | | |
|---|---|---|---|---|---|---|
| **Aplnr** | -0.3755288 | 0.77082284 | -4.8758707 | 0.00083628 | 0.02554907 | -0.2633115 |
| **Pcp4l1** | -0.3731143 | 0.77211397 | -4.1345296 | 0.00245542 | 0.04083645 | -1.5512793 |
| **Rin3** | -0.370154 | 0.77369988 | -3.2459007 | 0.00985208 | 0.08348522 | -2.4186054 |
| **Cyp2d22** | -0.369781 | 0.77389997 | -5.9405787 | 0.00020431 | 0.01473847 | 1.05233168 |
| **Thbs2** | -0.3694206 | 0.77409331 | -6.0851098 | 0.00017077 | 0.01439958 | 1.28268344 |
| **Car2** | -0.3693786 | 0.77411588 | -6.9197152 | 6.38E-05 | 0.01210215 | 2.17342372 |
| **Fam234a** | -0.369288 | 0.77416449 | -7.0515608 | 5.51E-05 | 0.01210215 | 2.36618073 |
| **Abca7** | -0.3659854 | 0.77593868 | -4.154696 | 0.00238204 | 0.04041084 | -1.4395691 |
| **Dpy19l1** | -0.3658538 | 0.77600948 | -6.9243249 | 6.35E-05 | 0.01210215 | 2.18017256 |
| **Itgb3** | -0.3633066 | 0.77738082 | -3.4726003 | 0.00685005 | 0.06913178 | -2.2146345 |
| **Gpd1** | -0.3626583 | 0.7777302 | -3.9728549 | 0.00313797 | 0.04670671 | -1.8042819 |
| **Josd2** | -0.3610818 | 0.77858055 | -8.522755 | 1.20E-05 | 0.00983403 | 3.8498852 |
| **Myo1e** | -0.3596364 | 0.77936099 | -5.1463716 | 0.0005758 | 0.02108837 | 0.02483653 |
| **Tmem88b** | -0.3592665 | 0.77956081 | -5.9738491 | 0.000196 | 0.01473847 | 1.03132106 |
| **Cd59a** | -0.3584067 | 0.78002558 | -6.1018406 | 0.00016729 | 0.01435693 | 1.27413611 |
| **Arhgef5** | -0.3561247 | 0.78126035 | -4.4592076 | 0.00151714 | 0.03222294 | -0.7578755 |
| **Rhpn2** | -0.3557781 | 0.78144804 | -3.8359476 | 0.0038731 | 0.05196265 | -1.6192207 |
| **Ldlr** | -0.3542463 | 0.7822782 | -4.9384976 | 0.00076634 | 0.02433804 | -0.3430931 |
| **Cldn11** | -0.3539883 | 0.78241812 | -6.5382793 | 9.90E-05 | 0.01356866 | 1.72658764 |
| **Rhog** | -0.3535083 | 0.78267851 | -6.8900711 | 6.60E-05 | 0.01210215 | 2.16287443 |
| **Hcn2** | -0.3529817 | 0.78296422 | -6.1325319 | 0.00016111 | 0.01435693 | 1.23136024 |
| **Cyp4f13** | -0.3526072 | 0.78316751 | -4.1266943 | 0.00248457 | 0.04103802 | -1.2078385 |
| **Pdzrn3** | -0.3524794 | 0.7832369 | -4.813726 | 0.00091251 | 0.02640938 | -0.2960181 |
| **Jam3** | -0.351335 | 0.78385843 | -7.4497256 | 3.57E-05 | 0.01161523 | 2.78939882 |
| **Foxc1** | -0.3503968 | 0.78436836 | -4.1505124 | 0.00239707 | 0.04041084 | -1.176445 |
| **Foxo4** | -0.3490686 | 0.78509077 | -7.6616348 | 2.85E-05 | 0.01130325 | 2.99427766 |
| **Foxo1** | -0.3481435 | 0.78559436 | -6.3795054 | 0.00011951 | 0.01356866 | 1.61943662 |
| **Fth1** | -0.347286 | 0.78606146 | -8.1004139 | 1.82E-05 | 0.01057257 | 3.43471191 |
| **Ly6c1** | -0.3472031 | 0.78610662 | -7.3534993 | 3.96E-05 | 0.01161523 | 2.67980897 |
| **Prodh** | -0.3460355 | 0.78674309 | -6.3766608 | 0.00011992 | 0.01356866 | 1.62225436 |
| **Zic2** | -0.3452975 | 0.78714563 | -6.4482919 | 0.00011013 | 0.01356866 | 1.6932429 |
| **Rasgrp3** | -0.3449403 | 0.78734055 | -5.9543017 | 0.00020084 | 0.01473847 | 1.10902281 |
| **Parp4** | -0.3446745 | 0.78748561 | -4.0158724 | 0.00293868 | 0.0448083 | -1.3883425 |
| **Ermn** | -0.34453 | 0.78756452 | -4.5499113 | 0.00132979 | 0.03026397 | -0.9321435 |
| **Sec14l5** | -0.3439645 | 0.7878733 | -4.0527314 | 0.00277859 | 0.04328868 | -1.6589665 |
| **Ntsr2** | -0.3424764 | 0.78868636 | -9.1464496 | 6.69E-06 | 0.00870145 | 4.42040114 |
| **Spsb1** | -0.3395197 | 0.79030439 | -4.3118757 | 0.00188409 | 0.03554058 | -1.2089196 |
| **Anxa4** | -0.3393447 | 0.79040022 | -3.3415688 | 0.00844577 | 0.07745614 | -2.3310237 |
| **Trip10** | -0.3392313 | 0.79046236 | -3.5548218 | 0.00601242 | 0.06503883 | -1.9925201 |

| | | | | | | |
|---|---|---|---|---|---|---|
| **Phactr4** | -0.3390853 | 0.79054239 | -5.9828136 | 0.00019383 | 0.01473847 | 1.11110807 |
| **Tubb4a** | -0.3388563 | 0.79066786 | -8.589455 | 1.13E-05 | 0.00983403 | 3.90989618 |
| **Tpm2** | -0.338588 | 0.79081491 | -3.7698279 | 0.00429131 | 0.05482825 | -1.774654 |
| **Irf8** | -0.3381393 | 0.79106089 | -3.3633866 | 0.00815531 | 0.0760261 | -2.2503754 |
| **Myh14** | -0.3380741 | 0.79109668 | -5.9376124 | 0.00020507 | 0.01473847 | 1.01093556 |
| **Inf2** | -0.3373371 | 0.79150092 | -5.7074452 | 0.00027445 | 0.01647115 | 0.7019003 |
| **Tmbim1** | -0.3366681 | 0.79186799 | -7.3079363 | 4.16E-05 | 0.01161523 | 2.6171519 |
| **Acrbp** | -0.3353204 | 0.7926081 | -3.3921685 | 0.00778797 | 0.07467945 | -2.2167277 |
| **Ldlrad3** | -0.3348381 | 0.79287309 | -6.1734388 | 0.00015325 | 0.01404592 | 1.37889002 |
| **Myrf** | -0.3344366 | 0.79309377 | -5.607496 | 0.00031218 | 0.01704154 | 0.55284088 |
| **Aass** | -0.3339742 | 0.79334801 | -4.1913026 | 0.00225472 | 0.03919965 | -1.12651 |
| **Plxnb3** | -0.3326803 | 0.79405989 | -6.4059769 | 0.0001158 | 0.01356866 | 1.57323647 |
| **Htra1** | -0.3326302 | 0.79408747 | -6.0292903 | 0.00018296 | 0.01462292 | 1.1101512 |
| **Ugt8a** | -0.3324295 | 0.7941979 | -4.5432873 | 0.0013426 | 0.03043199 | -0.9476421 |
| **Lgals9** | -0.3320643 | 0.79439898 | -4.5233774 | 0.0013819 | 0.0307147 | -0.6714949 |
| **Boc** | -0.331703 | 0.79459795 | -5.7332902 | 0.00026552 | 0.016358 | 0.86524808 |
| **Cpne9** | -0.3315849 | 0.794663 | -3.8560343 | 0.00375474 | 0.05135432 | -1.8697035 |
| **Carns1** | -0.3305406 | 0.79523841 | -4.7845963 | 0.00095078 | 0.02685043 | -0.4548271 |
| **Nkain2** | -0.3304156 | 0.79530734 | -7.8492335 | 2.35E-05 | 0.01130325 | 3.19603691 |
| **Matn2** | -0.3300655 | 0.79550038 | -4.2288372 | 0.00213166 | 0.03808088 | -1.0719322 |
| **Rhou** | -0.3299182 | 0.79558157 | -5.3734783 | 0.00042433 | 0.01918467 | 0.26835571 |
| **Lrp10** | -0.3298579 | 0.79561483 | -6.9309816 | 6.30E-05 | 0.01210215 | 2.23482653 |
| **Lrrc1** | -0.3276459 | 0.79683566 | -4.3124782 | 0.00188241 | 0.03554058 | -0.9888554 |
| **S100b** | -0.3273719 | 0.79698702 | -4.9561809 | 0.00074774 | 0.02406481 | -0.3469688 |
| **Erbb3** | -0.32729 | 0.79703225 | -3.609295 | 0.00551709 | 0.06219268 | -2.3499256 |
| **Gng11** | -0.3267736 | 0.79731759 | -4.1812225 | 0.00228904 | 0.03951812 | -1.3299134 |
| **Gjc2** | -0.3266572 | 0.7973819 | -5.1152403 | 0.00060075 | 0.02140612 | -0.0968047 |
| **Cntfr** | -0.3262125 | 0.79762772 | -7.8008325 | 2.47E-05 | 0.01130325 | 3.13822483 |
| **Tjap1** | -0.3262044 | 0.79763224 | -7.3101591 | 4.15E-05 | 0.01161523 | 2.6177951 |
| **Eva1b** | -0.3260517 | 0.79771667 | -3.3185873 | 0.00876337 | 0.07899051 | -2.3117024 |
| **Gja1** | -0.3240336 | 0.79883333 | -6.4061459 | 0.00011578 | 0.01356866 | 1.56608984 |
| **Hr** | -0.3235877 | 0.79908023 | -4.6633846 | 0.00112954 | 0.02836303 | -0.7325281 |
| **Rapgef3** | -0.3224553 | 0.79970774 | -7.8359288 | 2.38E-05 | 0.01130325 | 3.18423994 |
| **Ccdc191** | -0.3224549 | 0.79970792 | -3.6321259 | 0.00532239 | 0.06114223 | -1.8881608 |
| **Evi2a** | -0.3223683 | 0.79975596 | -4.9440665 | 0.00076043 | 0.02429026 | -0.2519276 |
| **Pnpla2** | -0.3219516 | 0.79998697 | -5.1819392 | 0.00054865 | 0.02057468 | 0.13294623 |
| **Fam222a** | -0.3215697 | 0.80019876 | -4.0630246 | 0.00273555 | 0.04291687 | -1.4281106 |
| **Ctsz** | -0.3213411 | 0.80032554 | -7.1298454 | 5.05E-05 | 0.01210215 | 2.4473313 |
| **Mobp** | -0.3212932 | 0.80035214 | -6.2173253 | 0.00014528 | 0.01397668 | 1.34906615 |

| | | | | | | |
|---|---|---|---|---|---|---|
| **Plekhh1** | -0.3212833 | 0.80035765 | -3.9954933 | 0.00303137 | 0.0455502 | -1.766435 |
| **Sh3tc2** | -0.3208637 | 0.80059044 | -4.1242667 | 0.00249368 | 0.04104942 | -1.2599357 |
| **Rabep2** | -0.319844 | 0.80115652 | -4.9057044 | 0.00080214 | 0.02500434 | -0.1879537 |
| **Nid1** | -0.3195564 | 0.80131624 | -7.414205 | 3.71E-05 | 0.01161523 | 2.73858687 |
| **Tmem98** | -0.3193493 | 0.80143125 | -5.4257245 | 0.00039596 | 0.01880988 | 0.44448303 |
| **Mfsd2a** | -0.3192661 | 0.8014775 | -6.1040346 | 0.00016684 | 0.01435693 | 1.21573499 |
| **Dio2** | -0.318989 | 0.80163145 | -5.9274535 | 0.0002077 | 0.01473847 | 1.01553961 |
| **Il17ra** | -0.3184371 | 0.80193814 | -5.3915579 | 0.00041427 | 0.01915025 | 0.43621969 |
| **Lrp4** | -0.3183536 | 0.80198455 | -3.7755635 | 0.00425321 | 0.05462771 | -1.9551254 |
| **Adamts2** | -0.3183523 | 0.80198532 | -6.2925269 | 0.00013265 | 0.01372423 | 1.52281012 |
| **Ctsa** | -0.3183403 | 0.80199199 | -5.6256979 | 0.00030491 | 0.01691691 | 0.59928221 |
| **Stard8** | -0.3180969 | 0.80212727 | -5.1967084 | 0.00053779 | 0.0204724 | 0.19505704 |
| **Unc93b1** | -0.3176093 | 0.80239846 | -5.3488219 | 0.00043847 | 0.01931297 | 0.38073847 |
| **Rida** | -0.3173371 | 0.80254982 | -6.6760573 | 8.43E-05 | 0.0135466 | 1.95830199 |
| **Scd3** | -0.3172017 | 0.80262516 | -4.1318743 | 0.00246526 | 0.04087914 | -1.1975045 |
| **Tmem125** | -0.3170384 | 0.80271603 | -5.5140975 | 0.00035254 | 0.0181571 | 0.50907624 |
| **Atp2a3** | -0.3169654 | 0.80275666 | -4.3129595 | 0.00188107 | 0.03554058 | -1.0081659 |
| **Spag5** | -0.3169059 | 0.80278973 | -4.0524008 | 0.00277998 | 0.04328868 | -1.4668571 |
| **Padi2** | -0.3164409 | 0.80304855 | -4.0988783 | 0.00259108 | 0.04193466 | -1.5942902 |
| **Sdc4** | -0.3162621 | 0.80314807 | -6.3340597 | 0.00012619 | 0.01356866 | 1.51676475 |
| **Hspb6** | -0.3155196 | 0.80356154 | -5.0469544 | 0.00065964 | 0.02256429 | 0.0075933 |
| **Daam2** | -0.3152959 | 0.80368613 | -7.427773 | 3.65E-05 | 0.01161523 | 2.73990237 |
| **Man2b1** | -0.3150802 | 0.80380629 | -6.5414449 | 9.87E-05 | 0.01356866 | 1.78259752 |
| **Cpt2** | -0.3149266 | 0.80389187 | -4.3159142 | 0.00187286 | 0.03548748 | -0.9876423 |
| **Fgfr2** | -0.3148487 | 0.80393529 | -5.7590734 | 0.00025693 | 0.0162317 | 0.78219064 |
| **Spp1** | -0.3146153 | 0.80406538 | -3.3783237 | 0.00796247 | 0.075291 | -2.6475702 |
| **Atp1a2** | -0.3145269 | 0.80411465 | -4.8203601 | 0.00090403 | 0.0263309 | -0.5359221 |
| **Prrg1** | -0.3141902 | 0.80430231 | -3.3755873 | 0.00799744 | 0.075291 | -2.5000549 |
| **Tprn** | -0.3135542 | 0.80465698 | -5.7439152 | 0.00026194 | 0.016358 | 0.78500185 |
| **Opalin** | -0.3114117 | 0.80585284 | -4.5776692 | 0.00127752 | 0.02959364 | -0.8503384 |
| **Lpar6** | -0.3108011 | 0.80619397 | -4.2836514 | 0.00196462 | 0.03624867 | -0.992253 |
| **Trp53inp2** | -0.3103148 | 0.80646578 | -7.0209128 | 5.70E-05 | 0.01210215 | 2.28521982 |
| **Nod1** | -0.3101908 | 0.80653511 | -5.690235 | 0.00028058 | 0.01649839 | 0.80795811 |
| **Pltp** | -0.3085337 | 0.80746202 | -5.9958168 | 0.00019072 | 0.01473847 | 1.12913462 |
| **Prr18** | -0.3085127 | 0.80747379 | -5.2905768 | 0.00047394 | 0.01999711 | 0.138067 |
| **Ifi27** | -0.3082744 | 0.80760718 | -3.3641995 | 0.00814469 | 0.07601114 | -2.3335436 |
| **Slc38a2** | -0.3081506 | 0.80767645 | -7.2254949 | 4.55E-05 | 0.01161523 | 2.51871247 |
| **Eva1a** | -0.3080495 | 0.80773309 | -4.506278 | 0.00141663 | 0.03098571 | -0.7204987 |
| **Mag** | -0.3079452 | 0.80779145 | -5.6966719 | 0.00027827 | 0.01649839 | 0.67189744 |

| | | | | | | |
|---|---|---|---|---|---|---|
| **Spg20** | -0.3079051 | 0.80781389 | -4.4353591 | 0.00157095 | 0.03295078 | -1.0608399 |
| **Phldb1** | -0.3071191 | 0.80825416 | -5.4980859 | 0.00036001 | 0.01825396 | 0.40365195 |
| **Gprc5b** | -0.3069997 | 0.80832104 | -5.1803951 | 0.0005498 | 0.02057468 | -0.0273293 |
| **Wnt7a** | -0.3064508 | 0.80862863 | -3.2800689 | 0.00932381 | 0.08144096 | -2.5829212 |
| **Lims2** | -0.3062457 | 0.80874362 | -7.0906073 | 5.27E-05 | 0.01210215 | 2.41343381 |
| **Frmd8** | -0.3057257 | 0.80903517 | -4.842229 | 0.00087666 | 0.0259922 | -0.4312774 |
| **Nkd1** | -0.3055492 | 0.80913414 | -4.9568946 | 0.000747 | 0.02406481 | -0.2392717 |
| **Slc4a2** | -0.3055413 | 0.80913859 | -4.6290629 | 0.00118647 | 0.02883077 | -0.7877496 |
| **Srpk3** | -0.3050941 | 0.80938943 | -3.9016526 | 0.00349988 | 0.04951509 | -1.6258142 |
| **Fcgrt** | -0.3049839 | 0.80945124 | -5.2815909 | 0.00047968 | 0.02001271 | 0.29631657 |
| **Eya2** | -0.3047047 | 0.80960793 | -4.1208279 | 0.00250664 | 0.04120077 | -1.242394 |
| **Aacs** | -0.304268 | 0.80985303 | -6.4746545 | 0.00010674 | 0.01356866 | 1.66537252 |
| **Pip4k2a** | -0.3041075 | 0.80994311 | -4.5442801 | 0.00134067 | 0.03042944 | -0.928095 |
| **Vim** | -0.303727 | 0.81015676 | -4.7591134 | 0.00098568 | 0.02729468 | -0.5467138 |
| **Cldn5** | -0.3033628 | 0.8103613 | -6.0182928 | 0.00018547 | 0.01468439 | 1.13661781 |
| **Olfml1** | -0.3032852 | 0.8104049 | -3.4501386 | 0.00709947 | 0.07071795 | -2.1730711 |
| **Acot11** | -0.3024171 | 0.8108927 | -4.4468245 | 0.00154483 | 0.03260571 | -1.0231524 |
| **Adi1** | -0.3019644 | 0.81114719 | -6.4277159 | 0.00011285 | 0.01356866 | 1.62420316 |
| **Ucp2** | -0.3018531 | 0.81120978 | -6.2143215 | 0.00014581 | 0.01397668 | 1.40650668 |
| **Myo1d** | -0.3016642 | 0.811316 | -3.3745929 | 0.00801019 | 0.075291 | -2.6765752 |
| **S1pr5** | -0.3014033 | 0.81146273 | -5.6288997 | 0.00030365 | 0.01691691 | 0.62622622 |
| **Cdh5** | -0.3011352 | 0.81161354 | -5.3976763 | 0.00041093 | 0.01914329 | 0.41867184 |
| **Pla2g7** | -0.2999487 | 0.81228129 | -4.6624998 | 0.00113097 | 0.02836303 | -0.7604744 |
| **Qdpr** | -0.2996317 | 0.81245978 | -5.70968 | 0.00027367 | 0.01647115 | 0.7333026 |
| **Heg1** | -0.2996315 | 0.81245992 | -5.5012292 | 0.00035853 | 0.01825396 | 0.45089046 |
| **Tlr3** | -0.2994502 | 0.81256202 | -3.3303158 | 0.00859976 | 0.07811765 | -2.5747346 |
| **Kif13b** | -0.2986678 | 0.81300277 | -3.5408671 | 0.00614666 | 0.06566202 | -2.4507603 |
| **P4ha1** | -0.2975596 | 0.81362756 | -5.5104429 | 0.00035423 | 0.0181571 | 0.50919208 |
| **Galnt6** | -0.2973988 | 0.81371824 | -3.7893757 | 0.00416293 | 0.05416945 | -1.9331114 |
| **Mir5125** | -0.297124 | 0.81387322 | -3.5284593 | 0.00626865 | 0.06605828 | -2.0991355 |
| **Cmtm5** | -0.2967458 | 0.81408661 | -6.0750634 | 0.0001729 | 0.01447525 | 1.19182834 |
| **Mob3b** | -0.2954896 | 0.81479577 | -6.2648878 | 0.00013715 | 0.01376723 | 1.42830834 |
| **Cdc42ep1** | -0.2954633 | 0.81481063 | -5.409203 | 0.0004047 | 0.01904097 | 0.33656514 |
| **Cdk18** | -0.2952883 | 0.81490948 | -6.9594651 | 6.11E-05 | 0.01210215 | 2.23537682 |
| **Gna12** | -0.2949176 | 0.81511886 | -6.3502347 | 0.00012377 | 0.01356866 | 1.50693238 |
| **Nkx6-2** | -0.294587 | 0.81530568 | -4.1634282 | 0.00235099 | 0.04012864 | -1.4138824 |
| **Usp54** | -0.2943168 | 0.81545842 | -4.8508132 | 0.00086616 | 0.0258531 | -0.4831181 |
| **Rffl** | -0.2940395 | 0.81561518 | -3.9515579 | 0.00324188 | 0.04755229 | -1.7616233 |
| **Pold4** | -0.2936978 | 0.81580838 | -3.4288279 | 0.00734489 | 0.07223097 | -2.1780638 |

| | | | | | | |
|---|---|---|---|---|---|---|
| **Acss2** | -0.293161 | 0.81611198 | -5.2430128 | 0.00050521 | 0.02009387 | 0.0888306 |
| **Lgr6** | -0.2930876 | 0.81615351 | -3.3884779 | 0.00783409 | 0.07476748 | -2.3237741 |
| **Glul** | -0.2922578 | 0.81662304 | -6.2248719 | 0.00014396 | 0.01397668 | 1.34222589 |
| **Slc22a4** | -0.2922312 | 0.81663809 | -4.1520457 | 0.00239155 | 0.04041084 | -1.2326028 |
| **Sgk3** | -0.2921292 | 0.81669583 | -4.9852504 | 0.00071823 | 0.02365658 | -0.0721996 |
| **Agpat4** | -0.2915503 | 0.8170236 | -7.0136197 | 5.75E-05 | 0.01210215 | 2.29669271 |
| **Hey2** | -0.2909072 | 0.8173879 | -3.3121767 | 0.00885416 | 0.07944569 | -2.3487659 |
| **Chst3** | -0.289863 | 0.81797975 | -4.587928 | 0.00125877 | 0.02948313 | -0.7074899 |
| **Zbtb7b** | -0.2896107 | 0.81812281 | -5.9132168 | 0.00021144 | 0.01473847 | 1.02294391 |
| **Sh3bp2** | -0.2889863 | 0.81847696 | -3.3116737 | 0.00886132 | 0.07944569 | -2.3383003 |
| **Bmp2** | -0.2886513 | 0.81866703 | -3.9177171 | 0.00341454 | 0.04888182 | -1.4920566 |
| **Cdkn1a** | -0.287673 | 0.81922234 | -3.9766318 | 0.00311991 | 0.04647897 | -1.4860577 |
| **Selenop** | -0.2876209 | 0.81925194 | -8.2415889 | 1.58E-05 | 0.010245 | 3.5737879 |
| **Smad7** | -0.2875668 | 0.81928266 | -6.435732 | 0.00011178 | 0.01356866 | 1.65225522 |
| **Mid1ip1** | -0.2871661 | 0.81951025 | -6.5651056 | 9.60E-05 | 0.01356866 | 1.77399587 |
| **Jph1** | -0.2869461 | 0.81963522 | -3.3186581 | 0.00876237 | 0.07899051 | -2.3803957 |
| **Phlda1** | -0.286079 | 0.820128 | -6.0304035 | 0.00018271 | 0.01462292 | 1.2172246 |
| **Gdpd5** | -0.2859591 | 0.82019616 | -6.1248769 | 0.00016263 | 0.01435693 | 1.30732244 |
| **Smox** | -0.285753 | 0.82031335 | -6.6413178 | 8.78E-05 | 0.01356866 | 1.87240743 |
| **Cdr2** | -0.2853518 | 0.82054151 | -3.5483715 | 0.00607408 | 0.06544692 | -2.3988597 |
| **Sox10** | -0.2853433 | 0.82054632 | -5.3484754 | 0.00043867 | 0.01931297 | 0.20918348 |
| **Card10** | -0.2852035 | 0.82062585 | -3.3202506 | 0.00873997 | 0.07899051 | -2.598371 |
| **Slc44a1** | -0.2851358 | 0.82066438 | -4.4110312 | 0.00162794 | 0.03335792 | -1.1427793 |
| **Ccdc80** | -0.2851128 | 0.82067747 | -5.2175295 | 0.00052287 | 0.02043832 | 0.22877882 |
| **Slco3a1** | -0.2850976 | 0.82068608 | -5.779978 | 0.00025018 | 0.01600944 | 0.80958092 |
| **Svil** | -0.2847638 | 0.82087599 | -5.9504817 | 0.0002018 | 0.01473847 | 1.12190687 |
| **Rftn1** | -0.2844906 | 0.82103148 | -3.9792637 | 0.00310739 | 0.04637442 | -1.5776519 |
| **Thrsp** | -0.2844476 | 0.82105593 | -3.799766 | 0.00409634 | 0.05375931 | -1.8276664 |
| **Pdgfd** | -0.2832602 | 0.82173195 | -3.5825567 | 0.00575465 | 0.06342905 | -2.07948 |
| **Nfe2l2** | -0.2831667 | 0.82178523 | -5.2657921 | 0.00048996 | 0.02001271 | 0.22474109 |
| **Aplp1** | -0.2829817 | 0.8218906 | -7.7385785 | 2.63E-05 | 0.01130325 | 3.06525476 |
| **Ushbp1** | -0.2829064 | 0.82193351 | -4.4283854 | 0.00158706 | 0.03303559 | -0.8340208 |
| **Natd1** | -0.2826059 | 0.82210475 | -6.2955645 | 0.00013217 | 0.01372423 | 1.48844646 |
| **Mettl7a1** | -0.2818831 | 0.82251669 | -4.6459896 | 0.00115802 | 0.02849755 | -0.5968965 |
| **Cnn2** | -0.2810527 | 0.82299029 | -4.4308202 | 0.00158142 | 0.03303559 | -0.7982702 |
| **Eng** | -0.2809109 | 0.82307117 | -5.2862682 | 0.00047669 | 0.01999711 | 0.20847993 |
| **H2afj** | -0.2806471 | 0.82322167 | -4.7714877 | 0.00096857 | 0.02699816 | -0.3875453 |
| **Agt** | -0.2805347 | 0.82328583 | -4.9228352 | 0.00078322 | 0.02459629 | -0.37313 |
| **Cgnl1** | -0.2803321 | 0.82340144 | -3.7591713 | 0.00436305 | 0.05536375 | -2.0968907 |

| | | | | | | |
|---|---|---|---|---|---|---|
| **Fah** | -0.2798067 | 0.82370137 | -5.324086 | 0.00045317 | 0.01934759 | 0.24290811 |
| **P3h4** | -0.279285 | 0.8239993 | -3.8776345 | 0.0036317 | 0.05053222 | -1.6471438 |
| **Etv1** | -0.2792796 | 0.8240024 | -4.2067619 | 0.00220315 | 0.03883375 | -1.3330017 |
| **Serpinh1** | -0.2780063 | 0.82472995 | -4.7800607 | 0.0009569 | 0.02685043 | -0.5051907 |
| **Osmr** | -0.2779205 | 0.82477898 | -4.2121338 | 0.00218552 | 0.03871493 | -1.0887822 |
| **Enpep** | -0.2777209 | 0.82489313 | -3.8544058 | 0.00376419 | 0.05141269 | -1.5775149 |
| **Irak4** | -0.2776718 | 0.82492117 | -3.2896503 | 0.00918101 | 0.0806819 | -2.3685916 |
| **Magt1** | -0.2775636 | 0.82498305 | -5.4190077 | 0.00039949 | 0.01887122 | 0.35287414 |
| **Gab1** | -0.2772632 | 0.82515487 | -4.9991196 | 0.00070459 | 0.02334411 | -0.2606237 |
| **Tradd** | -0.2770368 | 0.82528435 | -3.9962707 | 0.00302778 | 0.0455502 | -1.390364 |
| **Depdc1b** | -0.2767493 | 0.82544884 | -3.69445 | 0.00482673 | 0.05834729 | -1.854006 |
| **Afap1l1** | -0.2765516 | 0.82556194 | -5.9113588 | 0.00021193 | 0.01473847 | 1.03773986 |
| **Acp6** | -0.2764767 | 0.8256048 | -4.1640501 | 0.0023488 | 0.04012864 | -1.2423405 |
| **Acbd4** | -0.2759007 | 0.82593454 | -4.4432723 | 0.00155287 | 0.03269361 | -0.8310892 |
| **Piga** | -0.2755091 | 0.82615871 | -4.6388606 | 0.00116991 | 0.02863473 | -0.6984531 |
| **Laptm5** | -0.2751208 | 0.82638112 | -5.2693405 | 0.00048764 | 0.02001271 | 0.25620963 |
| **Sall1** | -0.2746952 | 0.82662495 | -4.425965 | 0.0015927 | 0.03303559 | -1.0503288 |
| **Aif1l** | -0.2740909 | 0.82697129 | -4.2801813 | 0.00197477 | 0.0363564 | -1.0190746 |
| **Igfbp7** | -0.2739091 | 0.82707545 | -6.8662252 | 6.78E-05 | 0.01229995 | 2.16365276 |
| **Itgb5** | -0.2731816 | 0.82749263 | -6.9185445 | 6.39E-05 | 0.01210215 | 2.22289869 |
| **Rpl13a** | -0.2713647 | 0.82853542 | -4.2590202 | 0.00203789 | 0.0369537 | -1.0377615 |
| **Mog** | -0.2711037 | 0.82868533 | -4.2784939 | 0.00197972 | 0.0364079 | -1.3498991 |
| **Col11a2** | -0.2703837 | 0.82909901 | -5.1407349 | 0.00058023 | 0.02108837 | -0.0660225 |
| **Igsf11** | -0.2700972 | 0.82926369 | -4.8523038 | 0.00086436 | 0.0258531 | -0.433678 |
| **Sipa1** | -0.2695916 | 0.82955434 | -5.3335386 | 0.00044749 | 0.01931297 | 0.33277775 |
| **Slain1** | -0.2695641 | 0.82957016 | -4.6481622 | 0.00115442 | 0.02848064 | -0.7593398 |
| **Scd2** | -0.2692541 | 0.82974845 | -5.2002767 | 0.0005352 | 0.0204724 | 0.01488458 |
| **Elovl1** | -0.2685425 | 0.83015782 | -4.9435302 | 0.000761 | 0.02429026 | -0.3394646 |
| **Stat2** | -0.2683696 | 0.83025732 | -4.1494026 | 0.00240107 | 0.04041084 | -1.3794399 |
| **Suox** | -0.2683304 | 0.83027984 | -4.8687329 | 0.00084468 | 0.02561018 | -0.2315098 |
| **Mxra8** | -0.2682944 | 0.83030058 | -3.9697347 | 0.00315297 | 0.04680604 | -1.453517 |
| **Plekha2** | -0.2682302 | 0.83033754 | -4.5953033 | 0.00124546 | 0.02929361 | -0.635094 |
| **Vamp3** | -0.2674215 | 0.83080309 | -4.9686271 | 0.00073495 | 0.02383497 | -0.2838845 |
| **Pnpla7** | -0.2667514 | 0.83118906 | -4.4873917 | 0.00145607 | 0.03152146 | -0.7485543 |
| **Mlc1** | -0.2665361 | 0.83131312 | -7.7359373 | 2.64E-05 | 0.01130325 | 3.06916949 |
| **Stard3** | -0.2664204 | 0.83137979 | -5.1717094 | 0.00055632 | 0.02061914 | 0.09894684 |
| **Tgfb1** | -0.2663289 | 0.83143255 | -3.648242 | 0.0051893 | 0.06064616 | -1.8981561 |
| **Rom1** | -0.2662281 | 0.83149063 | -3.3505064 | 0.0083255 | 0.07693221 | -2.2823861 |
| **Ldlrap1** | -0.2661578 | 0.83153113 | -3.8750352 | 0.00364628 | 0.05065849 | -1.8349468 |

| | | | | | | |
|---|---|---|---|---|---|---|
| **Rab7b** | -0.2661332 | 0.83154532 | -3.4820863 | 0.00674748 | 0.06875495 | -2.3440662 |
| **Mt3** | -0.2655204 | 0.83189861 | -7.1005214 | 5.22E-05 | 0.01210215 | 2.38234959 |
| **Pacsin3** | -0.2647637 | 0.83233505 | -4.9691652 | 0.0007344 | 0.02383497 | -0.2092236 |
| **Pdgfrb** | -0.2645189 | 0.83247631 | -4.4336886 | 0.00157479 | 0.03299039 | -0.9709559 |
| **Gstm7** | -0.264145 | 0.8326921 | -5.5636108 | 0.00033048 | 0.01747093 | 0.63132294 |
| **Adssl1** | -0.2639597 | 0.83279902 | -4.9339784 | 0.00077117 | 0.02442873 | -0.2866224 |
| **Glns-ps1** | -0.2631339 | 0.83327586 | -4.0395694 | 0.00283467 | 0.04365652 | -1.433384 |
| **Setd7** | -0.2625395 | 0.83361925 | -3.4437669 | 0.00717194 | 0.07114562 | -2.6536571 |
| **Arhgap29** | -0.262382 | 0.83371026 | -5.1979836 | 0.00053686 | 0.0204724 | 0.11381032 |
| **Cdc42bpg** | -0.2622253 | 0.83380085 | -3.559369 | 0.00596934 | 0.06465445 | -2.0449762 |
| **Pbxip1** | -0.2612676 | 0.83435453 | -6.3116765 | 0.00012963 | 0.01366306 | 1.47378259 |
| **Pacs2** | -0.2609763 | 0.834523 | -4.6172368 | 0.00120679 | 0.02894912 | -0.8411106 |
| **Tstd2** | -0.2602516 | 0.8349423 | -3.9106689 | 0.00345171 | 0.04921948 | -1.5855492 |
| **Clu** | -0.2600014 | 0.83508708 | -4.7183987 | 0.00104431 | 0.02790999 | -0.6917428 |
